# Chemical tools to expand the ligandable proteome: diversity-oriented synthesis-based photoreactive stereoprobes

**DOI:** 10.1101/2024.02.27.582206

**Authors:** Daisuke Ogasawara, David B. Konrad, Zher Yin Tan, Kimberly L. Carey, Jessica Luo, Sang Joon Won, Haoxin Li, Trever Carter, Kristen E. DeMeester, Evert Njomen, Stuart L. Schreiber, Ramnik J. Xavier, Bruno Melillo, Benjamin F. Cravatt

**Affiliations:** Department of Chemistry, Scripps Research, La Jolla, CA 92037, USA; Immunology Program, Broad Institute of MIT and Harvard, Cambridge, MA 02142, USA; Department of Molecular Biology, Massachusetts General Hospital, Harvard Medical School, Boston, MA 02114, USA; Center for Computational and Integrative Biology, Massachusetts General Hospital, Harvard Medical School, Boston, MA 02114, USA; Chemical Biology and Therapeutics Science Program, Broad Institute, Cambridge, MA 02142, USA; Department of Chemistry and Chemical Biology, Harvard University, Cambridge, MA 02138, USA

## Abstract

Chemical proteomics enables the global assessment of small molecule-protein interactions in native biological systems and has emerged as a versatile approach for ligand discovery. The range of small molecules explored by chemical proteomics has, however, been limited. Here, we describe a diversity-oriented synthesis (DOS)-inspired library of stereochemically-defined compounds bearing diazirine and alkyne units for UV light-induced covalent modification and click chemistry enrichment of interacting proteins, respectively. We find that these ‘photo-stereoprobes’ interact in a stereoselective manner with hundreds of proteins from various structural and functional classes in human cells and demonstrate that these interactions can form the basis for high-throughput screening-compatible nanoBRET assays. Integrated phenotypic analysis and chemical proteomics identified photo-stereoprobes that modulate autophagy by engaging the mitochondrial serine protease CLPP. Our findings show the utility of photo-stereoprobes for expanding the ligandable proteome, furnishing target engagement assays, and discovering and characterizing bioactive small molecules by cell-based screening.

## Introduction

Chemical probes offer powerful tools to acutely and dynamically perturb protein activities in biological systems, often in a bidirectional manner (losses or gains in function), and can even confer neo-functional outcomes^1-4^. Small molecules also represent a major category of drugs for treating human disease. Still, most human proteins lack selective chemical probes^5,6^, and this challenge has stimulated extensive innovation in the synthesis of small molecule libraries and ways to assay these compounds for interactions with proteins.

The process of chemical probe discovery often involves high-throughput screening (HTS) of large compound libraries against a protein of interest (target-based) or in a more complex biological system (phenotype-based)^7,8^. Target-based approaches frequently implement specialized functional assays that can be difficult to develop for poorly characterized proteins lacking well-defined biochemical activities. Alternative methods such as fragment-based ligand discovery^9,10^ and DNA-encoded library screening^11,12^ offer more general assays for identifying small molecules that bind to proteins, but still rely mostly on purified proteins. Phenotype-based screening, on the other hand, can identify chemical probes for proteins that are problematic to study in purified form (e.g., proteins that are components of large or dynamic complexes) or with biochemical functions that manifest mainly in cellular systems^13,14^. However, identifying the targets of phenotypic screening hits poses a significant hurdle^13,14^.

More recently, chemical proteomics has emerged as a versatile strategy for performing small-molecule probe discovery directly in native biological systems^15-19^. When combined with compound libraries that produce covalent interactions with proteins, either by intrinsic (e.g., electrophilic^20-23^) or induced (e.g., photoreactive^24-26^) mechanisms, chemical proteomics can identify, through mass spectrometry (MS) analysis, small molecules that bind to a wide range of proteins in cells often with clear structure-activity relationships (e.g., stereoselective and chemoselective binding). Chemical proteomics can also be integrated with phenotypic screening by using, for instance, ‘fully functionalized’ compounds possessing both (photo)reactive and latent affinity (e.g., alkyne) groups for the enrichment and identification of protein targets of bioactive small molecules in cells^24,27^. Attributes of chemical proteomics thus include in-depth maps of small molecule-protein interactions afforded by the analytical power of MS and a capacity to discover and optimize chemical probes targeting proteins in their endogenous cellular environment. Nonetheless, the throughput of chemical proteomic experiments is limited and more suitable for the analysis of focused (vs large) compound libraries.

The high-content, but lower throughput characteristics of chemical proteomics place emphasis on compound library design, and we and others have shown how properties such as stereochemistry are particularly well-suited to endow small sets of compounds with information-rich protein interaction profiles^21,23,25,27-29^. The design of stereochemically defined small-molecule libraries has further benefited from advances in synthetic organic chemistry, such as diversity-oriented synthesis (DOS), that provide access to architecturally complex cores with multiple stereocenters and sp^3^-carbon-rich rigid ring systems^30^. Compound libraries derived from DOS, when combined with phenotypic screening or chemical proteomics, have been productive sources of small-molecule probes that stereoselectively regulate protein function by atypical (e.g., allosteric) mechanisms^21,27,31-34^.

Chemical proteomic studies of DOS libraries have, so far, mainly focused on electrophilic compounds, such as acrylamides, that covalently bind to cysteine residues^21,27,28^. Given the expectation that many ligandable pockets throughout the proteome lack proximal cysteines, we sought herein to generate and globally map the protein binding profiles of a stereochemically defined, DOS-inspired photoreactive compounds (photo-stereoprobes). These compounds were designed to contain – 1) multiple structurally distinct cores, each possessing two stereocenters that present variable binding groups in different three-dimensional orientations; and 2) a common appendage bearing diazirine and alkyne groups for UV light-induced covalent binding and click chemistry-enabled enrichment of proteins from human cells, respectively. The photo-stereoprobes were found to stereoselectively enrich >200 structurally and functionally diverse proteins in human cancer cells, including those lacking chemical probes, such as adaptor/scaffolding and transcriptional regulatory proteins. The photo-stereoprobe-protein interactions displayed variable dependency on cellular environment, with some occurring only in situ, but not in vitro, and others not being recapitulated with recombinantly expressed protein, a property that correlated with membership in large protein complexes. Competition studies indicated that the photo-stereoprobes generally bound proteins with low affinity, but could nonetheless serve as the basis for target engagement assays compatible with high-throughput compound screening. Finally, we demonstrated the utility of photo-stereoprobes in phenotypic assays and report the discovery of compounds that stereoselectively modulate autophagy through engaging the mitochondrial protease CLPP.

## Results

### Design and initial profiling of DOS-inspired photo-stereoprobes

We selected tryptoline, azetidine, and pyrrolidine-based cores for the design of photo-stereoprobes, as these cores have relatively high skeletal and stereochemical complexity that can be accessed through DOS from inexpensive commercially available starting materials^35-37^ (**Figure 1A**). Analysis of the photo-stereoprobes by Plane of Best Fit (PBF), a chemical structural descriptor indicating the de-flatness of small molecules^38^, revealed significantly higher PBF scores compared to fragment-based photoreactive probes reported in other studies^24,25^ (**Figure S1A**), consistent with the DOS photo-stereoprobes having a higher degree of three-dimensionality. Tryptoline and azetidine acrylamides have also been extensively studied by cysteine-directed activity-based protein profiling^21,27,28^, which would thus allow for a comparison of proteomic interactions of electrophilic and photo-stereoprobes sharing common cores. We prepared a focused set of photo-stereoprobes consisting of four stereoisomers per core (4 stereoisomers × 3 cores = 12 photo-stereoprobes in total) (**Figure 1A**) such that chemical proteomic experiments performed in multiplexed format (see below) could instantly identify photo-stereoprobe-protein interactions displaying stereoselectivity, which is a prominent feature of small-molecule binding events occurring at defined pockets in proteins^39^.

**Figure 1.**
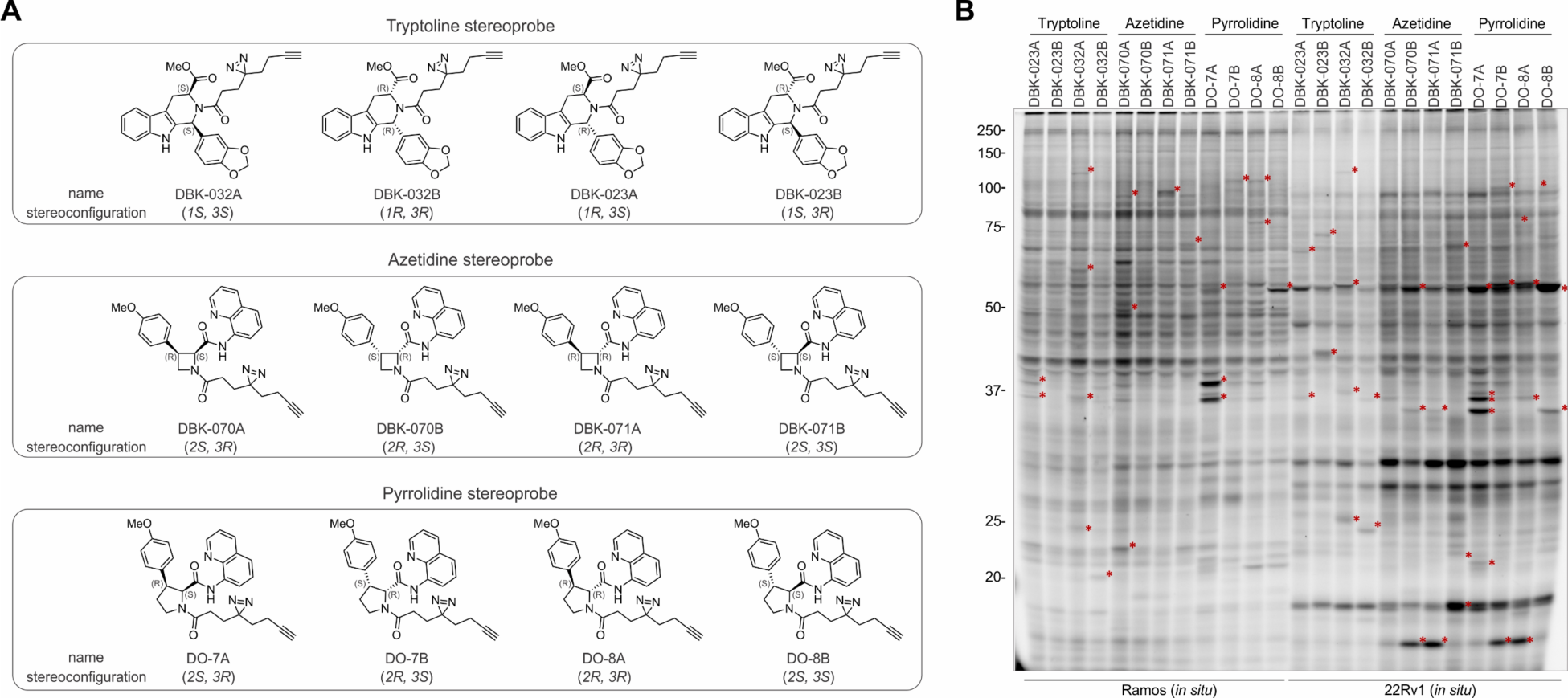
Photo-stereoprobes for mapping reversible small molecule-protein interactions in human cells. (**A**) Structures of photo-stereoprobes used in this study. (**B**) Gel profiling of stereoprobe-protein interactions in human cells. Ramos or 22Rv1 cells were incubated with stereoprobes (20 µM) for 30 min followed by UV cross-linking for 10 min. Stereoprobe-modified proteins in the soluble fraction were conjugated to a tetramethylrhodamine-azide (TAMRA-N_3_) reporter tag by CuAAC and analyzed by SDS-PAGE and in-gel fluorescence scanning. Red asterisks mark proteins showing stereoselective labeling by stereoprobes.

We first surveyed the proteomic interactions of photo-stereoprobes by gel-based profiling where the human Ramos (B lymphocyte) and 22Rv1 (prostate) cancer cell lines were treated with the photo-stereoprobes (20 µM, 30 min), followed by exposure to UV light (365 nm, 10 min), cell lysis, and stereoprobe-labeled proteins visualized by conjugation to a tetramethylrhodamine (TAMRA)-azide tag using Cu(I)-catalyzed azide–alkyne 1,3-dipolar cycloaddition (CuAAC) chemistry^40,41^, SDS-PAGE, and in-gel fluorescence scanning^42^. These experiments revealed similar overall degrees of proteomic engagement for the three photo-stereoprobe cores with clear stereoselective interactions observed with each stereoisomeric compound (**Figures 1B** and **S1B**). Virtually all of the proteomic labeling by the photo-stereoprobes was dependent on UV light (**Figure S1C**, **D**), supporting that these interactions reflect reversible binding events covalently trapped by photocrosslinking. Encouraged by the stereoselective protein binding events observed for the photo-stereoprobes in human cancer cells, we next performed an in-depth analysis of these interactions by quantitative MS-based proteomics.

### Global mapping of photo-stereoprobe-protein interactions in human cells

We analyzed the four stereoisomers of each photo-stereoprobe core in replicate experiments combined into a single multiplexed (16plex or 10plex) TMT (tandem mass tagging) proteomic experiment. These TMT experiments were performed separately in Ramos and 22Rv1 cells with 3–4 replicates each to yield 6–8 total replicates for each photo-stereoprobe in each cell line. In brief, cells were treated with photo-stereoprobes (20 µM, 1 h), UV cross-linked, lysed, and stereoprobe-modified proteins conjugated to biotin-azide by CuAAC reaction, enriched with streptavidin beads, trypsinized, and the tryptic peptides labeled with TMT reagents and identified (MS1/MS2 analysis) and quantified (MS3 analysis) using Orbitrap Fusion Tribid MS instruments (**Figure 2A**).

**Figure 2.**
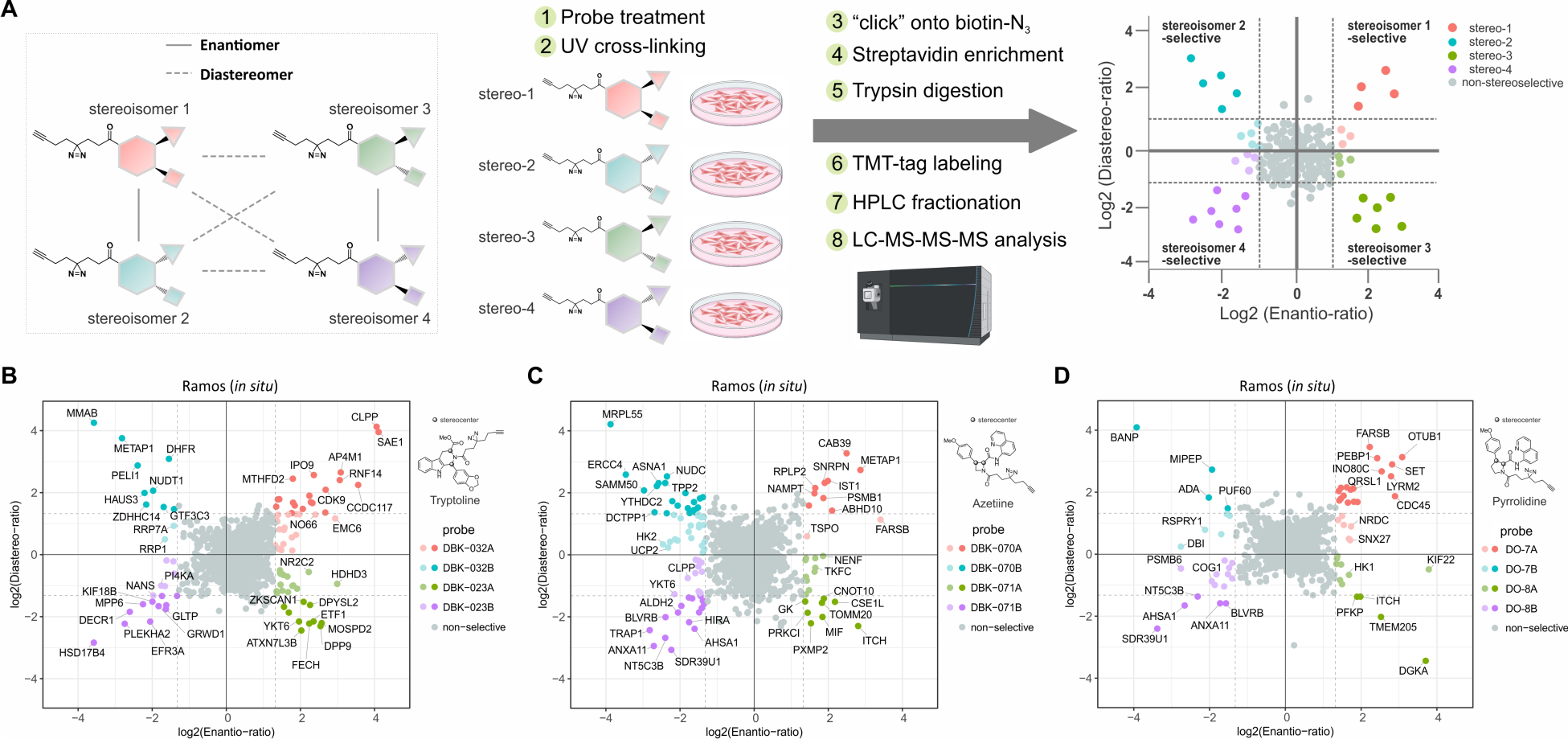
Mass spectrometry (MS)-based proteomic profiling of photo-stereoprobe-protein interactions in human cells. (**A**) Workflow for the MS-based analysis of photo-stereoprobe-protein interactions in human cells where relative protein enrichment across a full set of stereoisomeric probes is quantified by multiplexing (tandem mass tag (TMT)). (**B**–**D**) Quadrant plots displaying stereoselectively liganded proteins by tryptoline (**B**), azetidine (**C**), and pyrrolidine (**D**) photo-stereoprobes in Ramos cells. Enantio-ratio (x-axis) is the fold-enrichment ratio for the stereoisomer showing maximum engagement of a protein (probe^max^) over its enantiomer. Diastereo-ratio (y-axis) is the fold-enrichment ratio of probe^max^ over the diastereomer with higher relative engagement. Proteins that show an enantio-ratio >= 2.5 are colored as follows. Dark color: proteins that show enantio-ratio >= 2.5 and diastereo-ratio >= 2.5. Light color: proteins that show enantio-ratio >= 2.5 and diastereo-ratio < 2.5. Data represents mean values from at least two independent experiments.

We were principally interested in identifying proteins exhibiting high stereoselective engagement by the photo-stereoprobes (> 2.5-fold protein enrichment by one stereoisomer over its enantiomer), which we considered indicative of specific small molecule-protein interaction (or “liganding”) events (**Figure 2A**). Each photo-stereoprobe engaged a distinct set of proteins with high stereoselectivity, as visualized in quadrant plots where the positions of proteins on the x- and y-axes reflect enantioselective and diastereoselective liganding, respectively (**Figure 2B–D, Figure S2**, and **Supplementary Dataset 1**). We also generated in vitro photo-stereoprobe-protein interaction maps in Ramos cell lysates (**Figure S2D**–**F**; and see below).

In total, we mapped 392 stereoselective photo-stereoprobe-protein interactions that collectively corresponded to 253 stereoselectively liganded proteins in cancer cells, with >85% of these proteins showing preferential interactions with a single photo-stereoprobe (**Figure 3A** and **Supplementary Dataset 1**). Each photo-stereoprobe core stereoselectively liganded a similar number of proteins – 101, 105 and 83 proteins for the tryptoline, azetidine, and pyrrolidine probe sets, respectively (**Figure 3B**) – and stereoselective probe-protein interactions were observed for all four stereoisomers of each core set (**Figure 3C**). The majority of the stereoselectively liganded proteins lack established chemical probes, as evaluated using the Probe Miner database (**Figures 3B** and **S3A**), suggesting that the DOS photo-stereoprobes were engaging heretofore unliganded proteins. Also consistent with this conclusion, the stereoselectively liganded proteins were from a wide range of structural and functional classes, including those historically considered challenging to target with small molecules (e.g., transcriptional regulators and adaptors/scaffolding protein) (**Figure 3D**). Notably, we observed only a limited bias towards druggable protein classes like enzymes in the photo-stereoprobe-protein interaction map when compared to the global proteomes of Ramos and 22Rv1 cells (**Figure S3B**), supporting the broad ligandability potential of diverse protein classes, at least when assessed for small molecule interactions in native cellular environments. Of potential additional relevance to this interpretation, we noted that approximately half of the stereoselectively liganded proteins showed preferential interactions with photo-stereoprobes in living cells (*in situ*) compared to cell lysates (*in vitro*), while very few proteins showed the opposite profile (preferential liganding in lysates) (**Figure 3E-G**). Finally, a substantial subset of the stereoselectively liganded proteins (∼50%) are encoded by genes assigned as Common Essential or Strongly Selective in the Cancer Dependency Map^43^ (**Figure S3C**), indicating the importance of these proteins for general and specialized states of cell growth.

**Figure 3.**
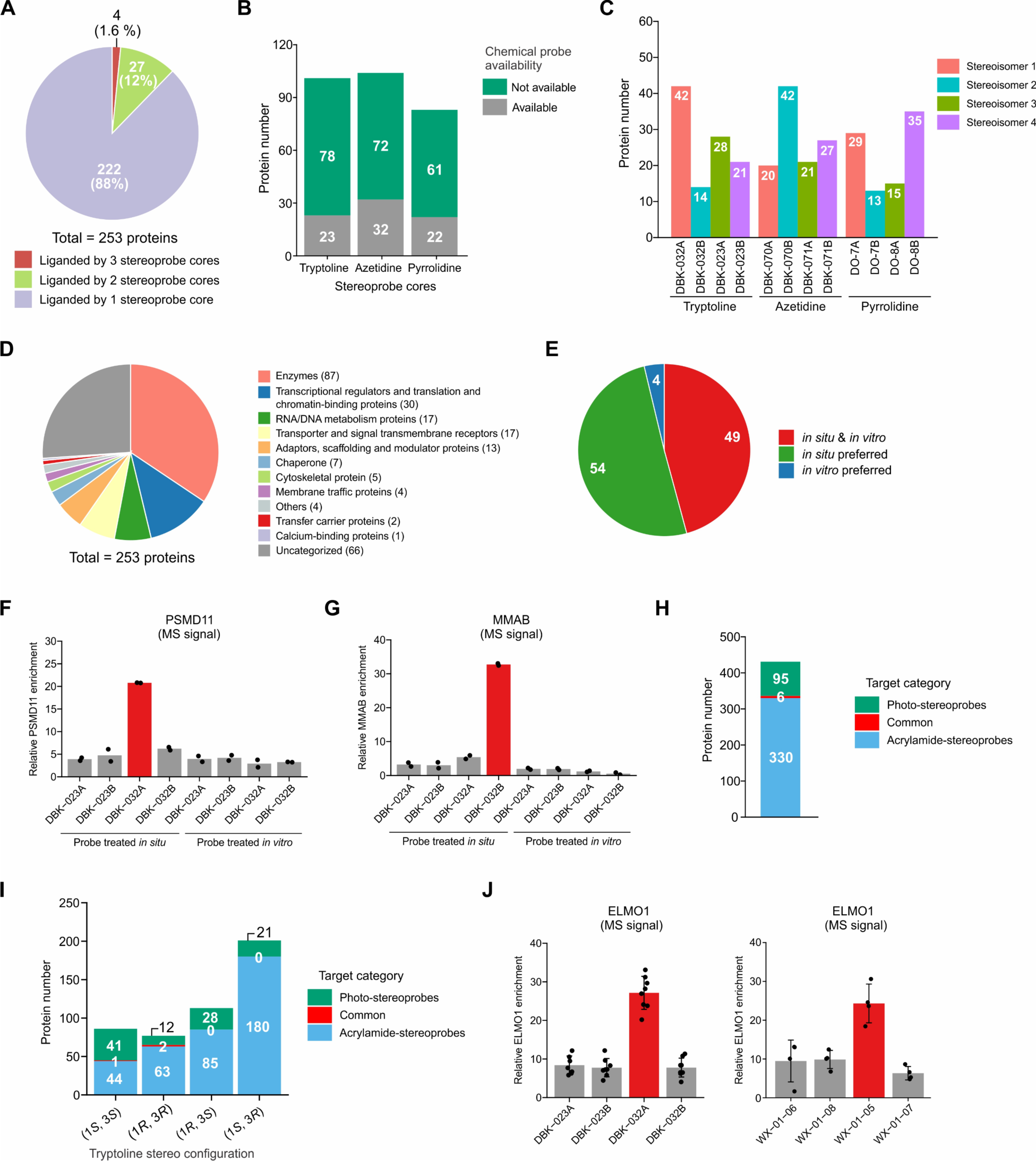
Features of proteins stereoselectively liganded by photo-stereoprobes. (**A**) Pie chart showing categorization of stereoselectively liganded proteins based on number of photo-stereoprobe cores targeting each protein. (**B**) Bar graph showing the proportion of stereoselectively liganded proteins with chemical probes as assigned by the Probe Miner database. (**C**) Bar graph showing numbers of stereoselectively liganded proteins by each stereoisomer of each photo-stereoprobe cores. (**D**) Pie chart showing functional classification of proteins stereoselectively liganded by photo-stereoprobes as assigned by the Panther database. (**E**) Pie chart showing proportion of proteins stereoselectively liganded by photo-stereoprobes *in situ* and/or *in vitro*. *In situ* & *in vitro*: proteins showing > 2.5-fold enantioselective enrichment in both *in situ* and *in vitro* data sets. *In situ* preferred: proteins showing > 2.5-fold enantioselective enrichment in *in situ* data sets and < 1.5-fold enantioselective enrichment in *in vitro* data sets. *In vitro* preferred: proteins showing > 2.5-fold enantioselective enrichment in *in vitro* data sets and < 1.5-fold enantioselective enrichment *in situ* data sets. (**F, G**) Comparison of *in situ* and *in vitro* enrichment profiles for (**F**) PSMD11 and (**G**) MMAB by tryptoline photo-stereoprobes in Ramos cells (*in situ*) or Ramos cell lysate (*in vitro*). Data represents mean values of two experiments per group. (**H, I**) Bar graphs showing proportion of proteins stereoselectively engaged by tryptoline photo-stereoprobes, tryptoline acrylamide stereoprobes, or both sets of tryptoline stereoprobes. Common targets were assigned as the proteins stereoselectively engaged by both sets of tryptoline stereoprobes, regardless of their stereo configurations (**H**), or with identical stereo configurations (**I**). Data for tryptoline acrylamide stereoprobes derived from ref 44. (**J**) Comparison of ELMO1 enrichment profiles by tryptoline photo-stereoprobes (left panel) and acrylamide stereoprobes (right panel) in Ramos cells. Data represents mean values of 4–8 experiments per group. For (**A**–**E**), data are combined for Ramos and 22Rv1 cells. For (**A**–**E**, **H**, **I**), proteins showing > 2.5-fold enrichment by one stereoprobe compared to its enantiomer were assigned as stereoselectively liganded proteins.

We next compared the ligandability maps of photo-stereoprobes to those of acrylamide stereoprobes constructed from the same core. Here, we focused on the tryptoline core, as we have recently performed a deep analysis of proteins liganded by tryptoline acrylamides using cysteine- and protein-directed ABPP^44^. The tryptoline photo- and acrylamide stereoprobes showed very little overlap in their respective ligandability maps (**Figure 3H**), and, interestingly, among the six shared proteins, three (ELMO1, GSTO1 and MRPL10) were engaged by photo- and acrylamide stereoprobes of the same stereoconfiguration (**Figure 3I, J**). Finally, we also found that the proteins stereoselectively interacting with DOS photo-stereoprobes were largely distinct from those stereoselectively enriched by fragment enantioprobes^25^ (**Figure S3D**).

Taken together, our chemical proteomic results indicate that photo-stereoprobes stereoselectively interact with a broad and distinctive (i.e., as-of-yet unliganded) array of proteins in human cells. We next selected a subset of these photo-stereoprobe-protein interactions for further characterization.

### Assessment of photo-stereoprobe interactions with recombinant proteins

We next sought to confirm the stereoselective interaction of photo-stereoprobes with representative proteins. For this analysis, we recombinantly expressed proteins from various structural and functional classes with a Flag-epitope tag in HEK293T cells by transient transfection, followed by treatment with photo-stereoprobes (20 µM unless indicated otherwise) for 30 min, UV cross-linking, and gel-based analysis of stereoprobe-protein interactions as described in **Figure 1**. We analyzed a total of 36 photo-stereoprobe targets, of which 22 showed the expected stereoselective interaction observed for endogenous forms of the proteins (**Figures 4** and **S4A**–**Y**). Examples include the E3 ubiquitin-protein ligase RNF14, the synaptobrevin homolog YKT6, and the transcriptional regulatory protein SET, which were consistently liganded as endogenous and recombinant proteins by DBK-032A, DBK-071B, and DO-7A respectively (**Figure 4A–C**).

**Figure 4.**
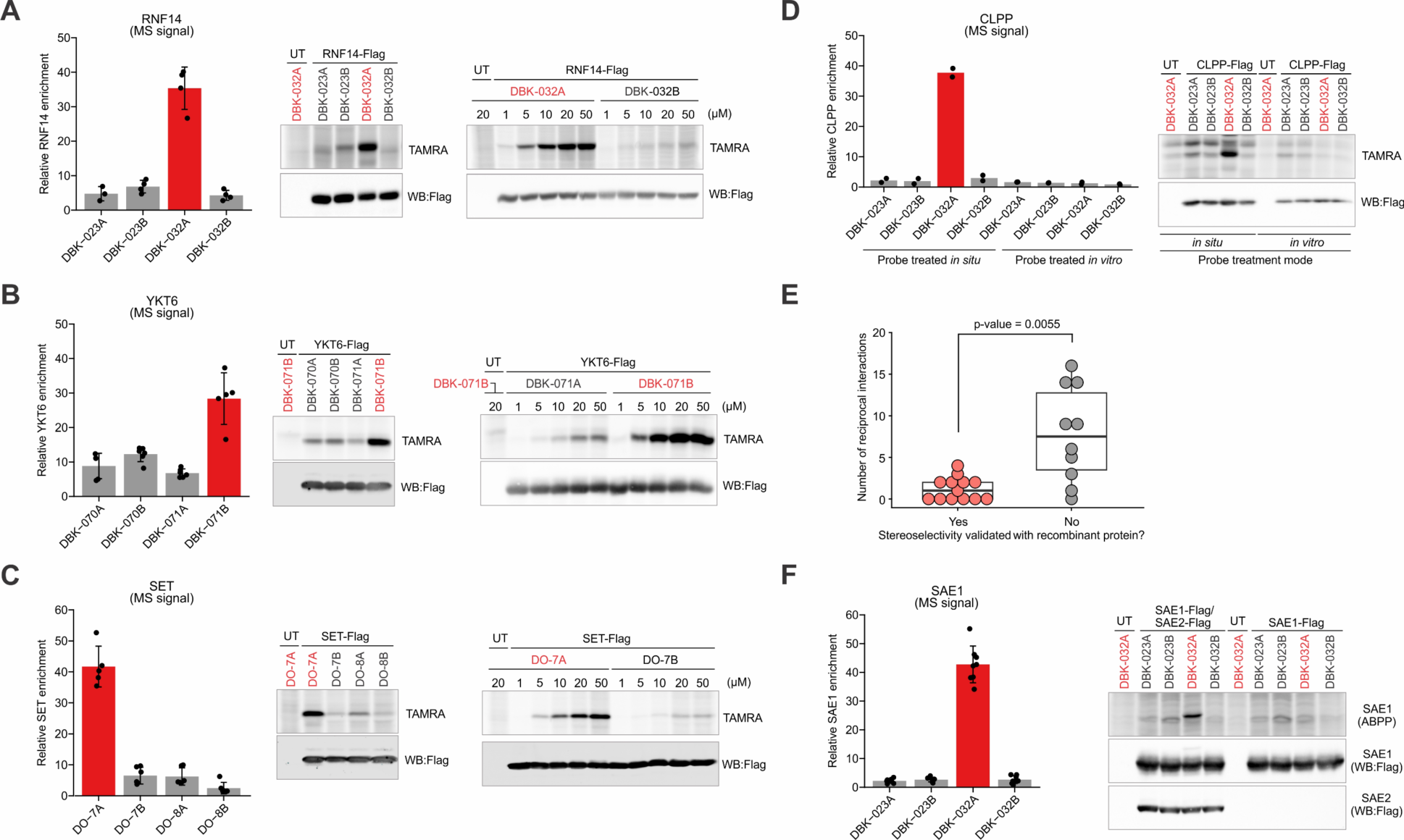
Characterization of proteins stereoselectively liganded by photo-stereoprobes. (**A**–**C**) Confirmation of *in situ* stereoselective engagement of RNF14 (**A**), YKT6 (**B**), SET (**C**) by indicated photo-stereoprobes. (**D**) Stereoselective engagement of CLPP by photo-stereoprobe DBK-032A *in situ*, but not *in vitro*. (**E**) Proteins that failed to display stereoselective engagement by photo-stereoprobes when tested in recombinant form displayed a higher number of reciprocal interactions with other proteins in the BioPlex interactome database. A protein-protein interaction detected as “Bait”-“Prey” in a bidirectional manner was considered as a reciprocal interaction. BioPlex 3.0 (HEK293T) was used as a reference database. (**F**) Stereoselective engagement of SAE1 by photo-stereoprobe DBK-032A requires presence of SAE1-interacting protein SAE2 (or UBA2). For (**A**–**D, F**), Left: relative enrichment profiles with the indicated photo-stereoprobes (20 µM) for RNF14 (**A**), YKT6 (**B**), SET (**C**), CLPP (**D**) and SAE1 (**F**) as determined by quantitative MS-based proteomics for Ramos cells (**A**–**D, F**) and/or Ramos cell lysates (**D**). Right: gel-based confirmation of stereoprobe interactions for proteins expressed recombinantly in HEK293T cells. RNF14 (**A**), YKT6 (**B**), SET (**C**), CLPP (**D**) and SAE1/SAE2 (**F**) were expressed with Flag epitope tags by transient transfection in HEK293T cells, after which the cells (**A**–**D, F**) and/or the lysates of transfected cells (**D**) were treated with the indicated stereoprobes (at 20 µM except where indicated) followed by UV cross-linking for 10 min. Probe-modified proteins were conjugated to TAMRA-N_3_ by CuAAC and analyzed by SDS-PAGE followed by in-gel fluorescence scanning. For (**A**–**C**) and (**F**), bar graph data represents mean values ± SD for at least four independent experiments. For (**D**), bar graph data represents mean values of two independent experiments per group.

In addition to stereoselectivity, the chemoselectivity of representative photo-stereoprobe-protein interactions was also confirmed with recombinant proteins. For example, Annexin A11 (ANXA11) was identified as a shared stereoselective target of both azetidine (DBK-071B) and pyrrolidine (DO-8B) probes of the same stereoconfiguration (**Figure S4A**, left bar graph), while SET stereoselectively engaged only the pyrrolidine probe, DO-7A, but not its stereochemically matched azetidine analogue, DBK-070A (**Figure S4B**, left bar graph), and recombinant ANXA11 and SET recapitulated these core selectivity profiles (**Figure S4A**–**B**, right panels). Photo-steroprobe-protein interactions showing consistent *in situ* and *in vitro* profiles were also verified with recombinant proteins (**Figure S4A**–**F**), as well as those showing divergent *in situ* and *in vitro* profiles like CLPP (**Figure 4D**). CLPP is a mitochondrial protease that exists as a homo-tetradecamer in association with an ATP-dependent unfoldase subunit CLPX to form a fully active protease complex^45-48^. While we do not yet understand why the stereoselective engagement of CLPP by DBK-032A was robustly observed *in situ*, but not *in vitro*, for both endogenous and recombinant forms of the protein (**Figure 4D**), it is possible that the oligomeric state of this protein or its interactions with CLPX are disrupted in cell lysates leading to a loss of photo-stereoprobe engagement.

Despite our generally high success rate in confirming photo-stereoprobe interactions with recombinant proteins, several exceptions were noted (**Figure S4F**–**H**). In most of these cases, we observed clear expression of the recombinant protein in HEK293T cells by anti-FLAG western blotting (**Figure S4F**–**H**), indicating that the failure to observe stereoselective engagement by the photo-stereoprobes was not related to poor protein expression. We next evaluated the profiles of photo-stereoprobe targets in the Bioplex database of protein-protein interactions^49^, which revealed that proteins failing in recombinant form to display the expected stereoprobe engagement were much more likely to be members of large multisubunit complexes (**Figure 4E**). This result suggested that some photo-stereoprobe-protein interactions may depend on an intact protein complex that is not properly assembled when one of the complex members is independently expressed. Consistent with this hypothesis, we found that the stereoselective engagement of SUMO-activating enzyme subunit 1 (SAE1) by DBK-032A was only recapitulated when this protein was co-expressed with its cognate binding partner SUMO-activating enzyme subunit 2 (SAE2) (**Figure 4F**).

Our analysis of recombinant protein targets thus supported the general veracity of liganding events for photo-stereoprobes, while also calling attention to a subset of these interactions that appear to require intact protein complexes, underscoring the value of performing small-molecule binding assays with endogenous proteins in their native cellular environments.

### Assessing the stoichiometry of photo-stereoprobe-protein interactions

Our assignment of proteins liganded by photo-stereoprobes was based on stereoselective enrichment, which does not address the affinity of the observed small molecule-protein interactions. As we and others have shown previously, the affinity of small molecule-protein interactions can be assessed in native biological systems with photoreactive compounds by competition experiments, wherein a non-photoreactive small molecule is tested for blockade of photoreactive probe binding to proteins^24,25,50^ (**Figures 5A** and **S5A**). We accordingly synthesized non-photoreactive propanamide analogs of representative tryptoline photo-stereoprobes for competition experiments (**Figure 5B**), where we would expect to observe preferential blockade of photo-stereoprobe enrichment by the corresponding stereoisomeric competitor (**Figure 5A** and **S5A**).

**Figure 5.**
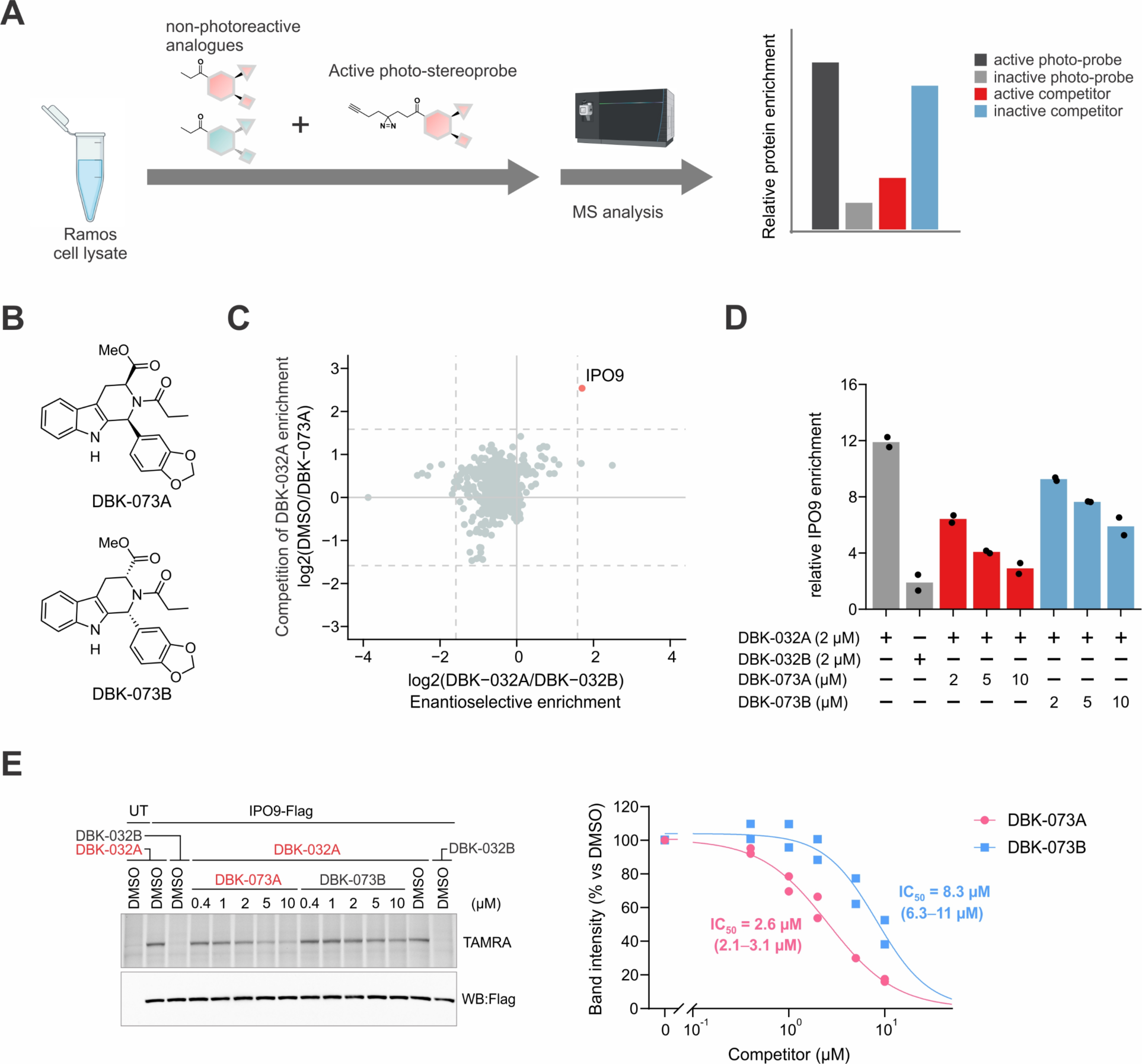
Assessing the stoichiometry of stereoprobe-protein interactions in cell lysates. (**A**) Workflow for the MS-based analysis of the stoichiometry of stereoprobe-protein interactions based on comparative quantification of protein enrichments from cell lysates co-incubated with DMSO or 4X (40 µM) non-photoreactive competitor stereoprobes and photo-stereoprobes (10 µM) of the same stereochemistry for 20 min. (**B**) Structures of non-photoreactive competitor stereoprobes DBK-073A/B. (**C**) Stereoselective enrichment of IPO9 by DBK-032A (x-axis) is blocked by competitor stereoprobe DBK-073A (y-axis). x-axis: log2 fold-enrichment ratio of DBK-032A (10 µM) over DBK-032B (10 µM). y-axis: log2 competition ratio for proteins enriched by DBK-032A (10 µM) in Ramos cell lysate pre-treated with DMSO or DBK-073A (40 µM). Dashed lines mark ratio cutoffs of > 3 for stereoselective enrichment (DBK-032A/DBK-032B) and competition of DBK-032A enrichment (DMSO/DBK-073A). (**D**) Concentration-dependent stereoselective blockade of DBK-032A enrichment of IPO9 by DBK-073A in comparison to enantiomer DKB-073B in Ramos cell lysates as determined by following the workflow shown in panel (**A**). Data represents mean values for two independent experiments per group. (**E**) Concentration-dependent stereoselective blockade of DBK-032A interactions with recombinant Flag-tagged IPO9 in lysates from transfected HEK293T cells as determined by gel-based analysis. Data represents mean values for two independent experiments per group.

Attempts to perform experiments *in situ* with excess concentration of propanamide competitors (e.g., 40 µM or above vs 10 µM photo-stereoprobe) encountered a technical challenge where the excess competitor caused a generally suppressive effect on photo-stereoprobe-protein interactions in cells (**Figure S5B**). We interpreted this result to indicate that the cellular uptake of the photo-stereoprobes was perturbed by high concentrations of competitors. We therefore performed initial competition experiments in Ramos cell lysates co-treated with DMSO or propanamide competitor (DBK-073A or DBK-073B (40 µM)) and corresponding photo-stereoprobes (DBK-032A or DBK-032B (10 µM)) for 20 min followed by UV light exposure (10 min) photocrosslinking and quantification of relative protein enrichment for each treatment condition by MS-based proteomics. The stereoselective enrichment of most protein targets of DBK-032A/B was generally unperturbed by competitor treatment (< 50% change), suggesting that these interactions are low affinity (**Figure 5C** and **Supplementary Dataset 1**). One exception was IPO9, for which the stereoselective enrichment by DKB-032A was substantially blocked by DBK-073A (**Figure 5C**). The competition displayed by DBK-073A was greater than that observed for the enantiomer DBK-073B across a concentration range of 2-10 µM (**Figure 5D**), yielding an IC_50_ value for stereoselectively blockade of DBK-032A interactions with recombinant IPO9 of 2.6 µM (2.1-3.1 µM) (**Figure 5E**). We also confirmed stereoselective *in situ* competition of DBK-032A interactions with endogenous and recombinant IPO9 by DBK-073A (**Figure S5C, D**).

Taken together, our initial competition studies indicate that most of the photo-stereoprobe-protein interactions mapped in our chemical proteomic studies are low affinity with rare exceptions like IPO9, where high stoichiometry engagement displays the expected stereoselectivity.

### Adapting photo-stereoprobe-protein interactions for high-throughput screening

In assessing the output of our initial chemical proteomic studies of photo-stereoprobes, we inferred that these experiments were successful at illuminating novel ligandable pockets on diverse proteins in cancer cells; however, the majority of the photo-stereoprobe-protein interactions also appeared to be low affinity. We therefore contemplated potential ways to convert global maps of stereoselective small molecule-binding into a general workflow for high-throughput screening of larger compound libraries against stereoprobe targets of interest. We pursued this objective using NanoBRET^51,52^, which, combined with our overall success at recapitulating photo-stereoprobe interactions with recombinant proteins expressed in HEK293T cells, offered a potential way to screen small molecules for binding to photo-stereoprobe targets in cell lysates. In this NanoBRET assay, a fluorescent version of the photo-stereoprobe serves as a BRET (bioluminescence resonance energy transfer) tracer for quantitatively assessing engagement to a target protein fused with nanoLuciferase (NLuc) (**Figure 6A**). Importantly, the quality of the assay can be assessed by comparing enantiomeric photo-stereoprobe pairs, where a “specific BRET signal” is defined as the difference in BRET intensity between the preferred versus nonpreferred enantiomeric tracer (**Figure 6A**). The binding of small molecules to the NLuc-fused protein target is then measured by suppression of the specific BRET signal (**Figure S6A**).

**Figure 6.**
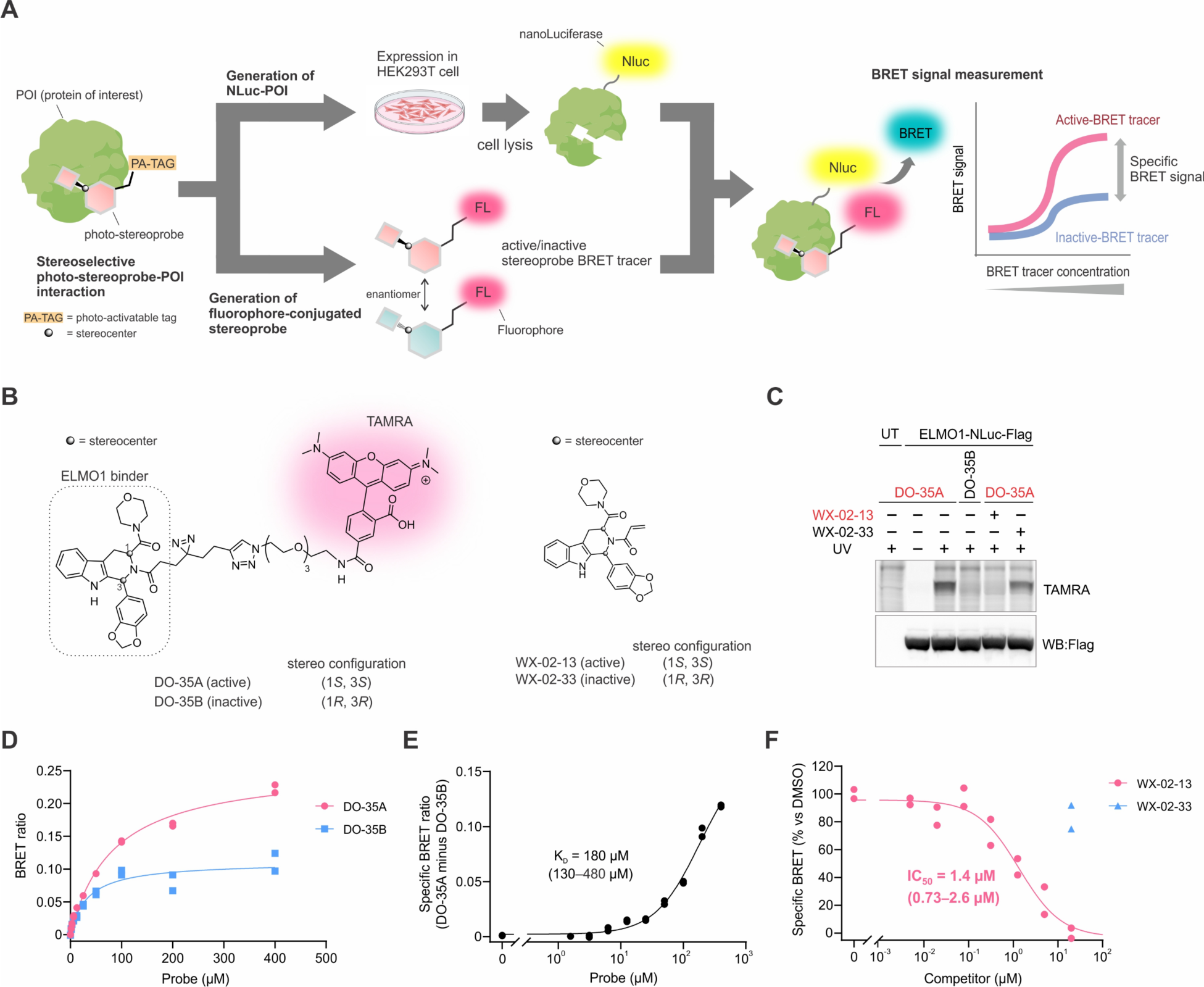
Converting photo-stereoprobe-protein interactions into a high-throughput screening-compatible nanoBRET assay. (**A**) Schematic workflow for development of a nanoBRET assay based on a photo-stereoprobe that stereoselectively engages a protein of interest (POI). (**B**) Structures of BRET tracer DO-35A for ELMO1, inactive enantiomer DO-35B, and electrophilic stereoprobe competitors WX-02-13 (active enantiomer) and WX-02-33 (inactive enantiomer). (**C**) Stereoselective engagement of ELMO1 by DO-35A. Lysate of HEK293T cells recombinantly expressing an ELMO1-NLuc-Flag fusion protein was incubated with DMSO, WX-02-13, or WX-02-33 (20 µM compounds, 1 h) followed by DO-35A or DO-35B (100 µM, 20 min) and UV cross-linking (10 min). The samples were then analyzed by SDS-PAGE followed by in-gel fluorescence scanning. (**D**) Concentration-dependent and stereoselective BRET signals for ELMO1-NLuc-Flag-expressing HEK293T cell lysates treated with DO-35A or DO-35B. BRET signals were measured after 20 min incubation. Data are from two independent experiments. (**E**) Specific BRET signal for DO-35A interaction with ELMO1-NLuc-Flag as determined by subtracting the BRET signal generated with DO-35B from the BRET signal generated with DO-35A in (**D**). (**F**) WX-02-13, but not WX-02-33 blocked the BRET signal for DO-35A interaction with ELMO1-NLuc-Flag in a concentration-dependent manner. BRET tracers (100 µM) and indicated concentrations of WX-02-13 and WX-02-33 were incubated with ELMO1-NLuc-Flag-expressing HEK293T cell lysates as described in (**C**). Data are from two independent experiments.

As a model protein for NanoBRET assay development, we selected the scaffolding protein ELMO1 (Engulfment and cell motility protein 1)^53^ as it was one of the few proteins stereoselectively liganded by both photoreactive and electrophilic tryptoline stereoprobes (**Figure 3J**), which allowed for the use of electrophilic stereoprobes as additional tools for NanoBRET assay development and validation. Previous cysteine-directed ABPP experiments identified the morpholine amide-substituted tryptoline acrylamide WX-02-13 (**Figure 6B**, right) as a stereoselective covalent ligand targeting C438 of ELMO1 in Ramos cells^34^, and we confirmed this interaction with recombinant ELMO1 expressed in HEK293T cells (**Figure S6B**). We then found that the corresponding morpholine amide tryptoline photo-stereoprobe (WX-02-18) stereoselectively engaged recombinant ELMO1 to a greater extent than the original photo-stereoprobe DBK-032A (**Figure S6C, D**). The engagement of both endogenous and recombinant ELMO1 by WX-02-18 was blocked by pre-treatment with WX-02-13, but not the inactive enantiomer WX-02-33 (**Figure S6B**, **E, F**), indicating that the electrophilic and photoreactive tryptoline stereoprobes bind the same pocket of ELMO1. Also consistent with this premise, a C438A-ELMO1 mutant failed to react with either electrophilic or photoreactive tryptoline stereoprobes (**Figure S6B, E**). Docking studies identified a potential small molecule binding pocket in proximity to C438 of ELMO1 that rationalized the preferential binding of (*S*, *S*) tryptoline compounds (e.g., WX-02-13) compared to the (*R*, *R*) enantiomers (e.g., WX-02-33) (**Figure S6G**).

Having identified both photoreactive and electrophilic stereoprobes that react stereoselectively and site-specifically with ELMO1, we synthesized fluorescent photo-stereoprobes DO-35A and DO-35B where a TAMRA fluorophore was conjugated by CuAAC to the alkyne group of WX-02-18/38 (**Figure 6B**), as this conjugation was predicted based on structural modeling to allow the linker and TAMRA group to exit the protein toward solvent (**Figure S6G**). We confirmed that DO-35A stereoselectively engaged a NLuc-tagged ELMO1 (NLuc-ELMO1) in transfected HEK293T cell lysates and that this interaction was UV light-dependent manner and blocked by WX-02-13 but not by WX-02-33 (**Figures 6C** and **S6H**). We next performed a NanoBRET assay in NLuc-ELMO1-transfected HEK293T cell lysates aliquoted into a 384-well format and found that DO-35A produced a concentration-dependent BRET signal that exceeded the signal of the enantiomer DO-35B (**Figure 6D**). We quantified specific BRET signals by subtracting the values for DO-35B from DO-35A at each concentration, from which we then calculated an estimated K_D_ value of 180 µM (95% confidence interval (CI) of 130–480 µM) for DO-35A binding to NLuc-ELMO1 (**Figure 6E**). No difference in BRET signals was observed for DO-35A and DO-35B in HEK293T cell lysates expressing a NLuc-only construct lacking ELMO1 (**Figure S6I**). Gel filtration eliminated the BRET signal for DO-35A with NLuc-ELMO1 (**Figure S6J**), supporting the reversibility of this interaction. Introducing a photocrosslinking step prior to gel filtration preserved a minor fraction of the total BRET signal that was much stronger for DO-35A compared to DO-35B (**Figure S6J**).

We next performed a competitive NanoBRET assay, which revealed the concentration-dependent blockade of specific BRET signal for NLuc-ELMO1 by WX-02-13, but not WX-02-33 (pre-treatment of 60 min), with an IC_50_ value of 1.4 µM (95% CI of 0.73-2.6 µM) (**Figure 6F**) that was similar to the IC_50_ value measured by gel-based analysis (**Figure S6K**). Similar competitive NanoBRET and gel-based assays were performed with propanamide analogs DO-40A and DO-40B, which revealed stereoselective blockade by enantiomer DO-40B with an IC_50_ value of 70 µM (95% CI of 50-110 µM) (**Figure S6L, M**). Finally, we validated the NanoBRET assay with a second protein IPO9, for which BRET tracers based on DBK-032A/B (DO-39A/B; **Figure S6N**) showed the expected stereoselective engagement of NLuc-IPO9 (**Figure S6O, P**) with an estimated K_D_ value for DO-39A of 13 µM (95% CI of 7.9-32 µM) calculated from the specific BRET signal (**Figure S6Q, R**) and the stereoselective blockade of this interaction by DBK-073A compared to the enantiomer DBK-073B (**Figure S6S, T**).

Taken together, these data indicate that photo-stereoprobe-protein interactions, despite showing generally low affinity (K_D_ values > 10 µM), can be converted into nanoBRET assays compatible with (at least) 384-well assay formats to measure the binding of small molecules to recombinant NLuc fusion proteins expressed in mammalian cell lysates.

### Phenotypic screening of photo-stereoprobes identifies autophagy-modulating compounds

Previous studies have shown that phenotypic screening of photoreactive compound libraries can enable target identification by chemical proteomics without requiring synthetic modification of hit compounds^24,54^. For such studies, structurally related inactive control compounds are critical for illuminating proteins that selectively interact with the hit compounds. We surmised that DOS photo-stereoprobes might prove useful for phenotypic screening because hit compounds that produce stereoselective biological effects are accompanied by physicochemically matched inactive enantiomeric control stereoprobes for subsequent target identification experiments^21,27^.

We screened photo-stereoprobes for effects on the cellular process of autophagy using an LC3 puncta induction assay as an indicator of autophagosome formation^55,56^. Autophagy is an essential intracellular degradation pathway for toxic macromolecules such as protein aggregates and invading pathogens^57-59^. To date, only a limited number of small-molecule modulators of autophagy have been identified, many of which also produce broader cellular effects (e.g., MTOR inhibitors)^60-65^. We treated HeLa cells stably expressing GFP-tagged LC3 with photo-stereoprobes for 4 h, and then LC3 puncta formation was quantified by fluorescence microscopy (**Figure 7A**). A single compound– the tryptoline DBK-032A – was identified as a hit that stereoselectively promoted LC3 puncta formation in a concentration-dependent manner with EC_50_ of 3.7 µM (95% CI of 3.3–4.2 µM) (**Figures 7B** and **S7A**). The effect of DBK-032A on autophagic flux was further confirmed by an LC3-II immunoblotting assay showing a stereoselective increase in LC3-I to LC3-II conversion (**Figure 7C**). An additional focused structure-activity relationship (SAR) analysis identified analogues with largely preserved (WX-02-22, WX-03-93) attenuated (WX-03-97), or substantial reduction in (WX-02-18) LC3 puncta activity (**Figure 7D**). Each active analogue maintained stereoselectivity (**Figure 7D**). We next performed MS-based proteomics to identify proteins preferentially enriched by DBK-032A compared to the inactive enantiomer DBK-032B and morpholino analogues WX-02-18 and WX-02-38. Five proteins (CLPP, IPO9, SAE1, EMC6, and PIR) were specifically enriched by DBK-032A (**Figure 7E** and **Supplementary Dataset 1**). Additional chemical proteomic studies with other photo-stereoprobe analogs (WX-02-22, WX-03-93, WX-03-97, and their respective enantiomers) revealed an SAR for protein enrichment that correlated best with the LC3 puncta assay for CLPP and IPO9 (**Figure 7F** and **Supplementary Dataset 1**). Other candidate targets such as SAE1 and EMC6 showed lower correlation in SAR because they were not engaged by WX-02-22 (**Figure S7B**), a compound displaying activity in the LC3 puncta formation assay (**Figure 7D**).

**Figure 7.**
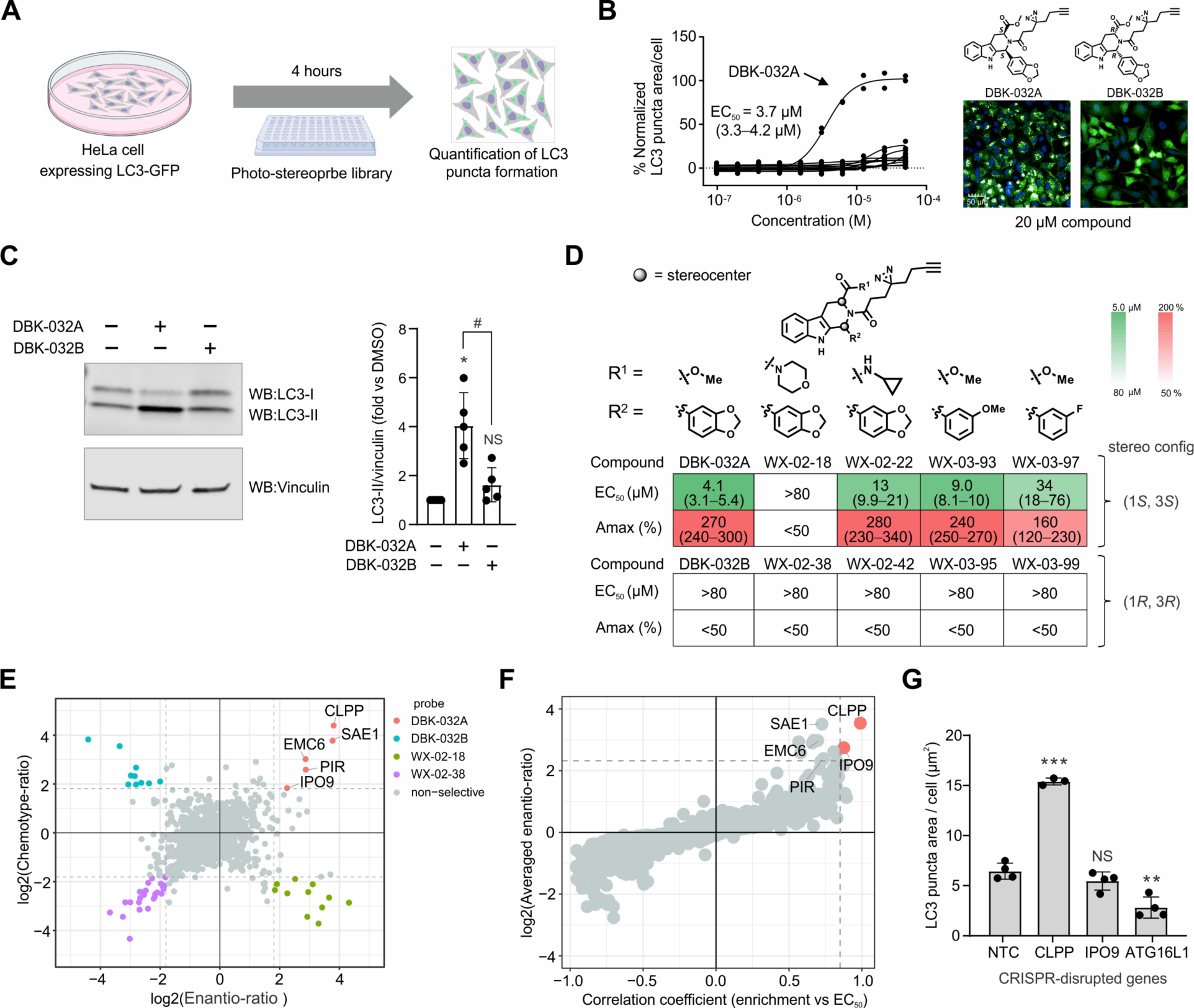
Integrated phenotypic screening and chemical proteomics identifies photo-stereoprobes that modulate autophagy by engaging CLPP. (**A**) Schematic for phenotypic screen to identify small-molecule modulators of autophagy. HeLa cells stably expressing LC3-GFP were treated with a photo-stereoprobe library for 4 h, after which LC3 puncta formation was measured by fluorescence microscopy. (**B**) DBK-032A stereoselectively promoted LC3 puncta formation with an EC_50_ value of 3.7 µM. Data are baseline-corrected with DMSO-treated samples and expressed as the % LC3 puncta area/cell relative to the cells treated with the mTOR inhibitor PI-103 (5 µM). Data represent mean values for two independent experiments per group. (**C**) DBK-032A stereoselectively promoted autophagic flux as assessed by LC3-II immunoblotting. Left gel image: a representative gel image for LC3 and vinculin (loading control) quantification. Right bar graph: normalized LC3-II levels quantified by immunoblotting. Bar graph data represents average values ± SD (n = 5 per group). *P < 0.05 (one sample Student’s t-test performed relative to DMSO control group); ^#^P < 0.05 (two samples Student’s t-test performed for DBK-032A and DBK-032B groups). NS: not significant. (**D**) Structures and LC3 puncta formation activities of DBK-032A analogues. EC_50_ and Amax (% maximum activation relative to the cells treated with 5 µM PI-103) represent average values (95% confidence interval) for at least four independent experiments for active stereoprobes. (**E**) Quadrant plots highlighting proteins showing greatest stereoselective enrichment by DBK-032A versus inactive analogues DBK-032B, WX-02-18, and WX-02-38. (**F**) CLPP and IPO9 showed the closest SAR correlation of stereoprobe engagement with LC3 puncta formation activity. x-axis: Pearson correlation coefficient of relative protein engagement in MS-based experiment and EC_50_ values from panel **D** (for Pearson correlation coefficient calculation, EC_50_ of the inactive probes were regarded as 80 µM). y-axis: Averaged fold-protein enrichment ratio between (*S*, *S*)- and (*R*, *R*)-isomers of each probe chemotype. For (**E**) and (**F**), HeLa cells stably expressing LC3-GFP were treated with corresponding photo-stereoprobes (20 µM, 1 h). Relative protein enrichment from whole cell lysate (**D**) or soluble fraction (**E**) was quantified following the workflow described in Figure 2. (**G**) Effect of CRISPR-mediated disruption of indicated genes on basal LC3 puncta formation in HeLa cells. ATG16L1 was included as a control protein required for autophagy induction. NTC: non-targeting control. Data represent average values ± SD (n = 3–4 per group). **P < 0.01; ***P < 0.001 (two-sided Student’s t-test performed relative to NTC control group). NS: not significant.

We next performed genetic studies to assess the biological relevance of CLPP and IPO9 to autophagy. Both siRNA-mediated knockdown and CRISPR-mediated knockout of CLPP, but not IPO9, significantly promoted the basal LC3 puncta formation activities in HeLa cells (**Figures 7G** and **S7C**, **D**) without affecting cell viability (**Figure S7E**). We further confirmed that siRNA-mediated knockdown of CLPP or CLPX, but not IPO9, significantly attenuated the LC3 puncta formation activity of DBK-032A (**Figure S7G**). These findings support that CLPP is the likely functional target for the autophagy-inducing effects of DBK-032A and are consistent with previous literature also showing that siRNA-mediated knockdown of this protein promotes autophagy^66,67^, which may relate to the role that this protein plays in mitochondrial protein quality control^68-70^.

## Discussion

Original chemical proteomic studies attempting to map the ligandabilty of protein in human cells mainly explored focused libraries of small-molecule fragments^20,24,71,72^. The limited SAR afforded by such experiments can frustrate confident assignment of small molecule binding events occurring at ‘actionable’ druggable pockets on proteins. This challenge has more recently been addressed by incorporation of defined stereocenters^21,23,25^, as well as more elaborated (DOS-inspired) structural cores^21,28^, into small molecule library designs for chemical proteomic investigation. These efforts have identified a large and diverse array of stereoselective small molecule-protein interactions in human cells, including many that occur on protein classes considered challenging for chemical probe development (e.g., DNA/RNA-binding proteins, adaptor proteins^27,34,44^). Nonetheless, much of the focus of chemical proteomics has been directed toward electrophilic compound libraries that covalently modify proteins. Our efforts here to unite DOS-inspired scaffolds with photoreactive chemistry offer a complementary way to enrich our understanding of the ligandable proteome.

We are encouraged that even with a small, focused library of DOS-inspired photo-stereoprobes, many stereoselective small molecule-protein interactions were uncovered, and most of these events occurred on proteins that have not yet been liganded by electrophilic or fragment-based photo-stereoprobes (or small molecules more generally). Our chemical proteomic results are consistent with previous assessments of DOS libraries as a source of compounds that show distinctive protein-binding and biological activities^73,74^. A clear difference, however, in the outputs of chemical proteomic studies of electrophilic and photo-stereoprobes relates to the stoichiometry of observed protein interactions, as electrophilic stereoprobes have been found to stereoselectively engage many cysteines on proteins with high stoichiometry^21,27,28,44^, while our data indicate that photo-stereoprobe-protein interactions are generally low occupancy events. This is perhaps unsurprising because the photo-stereoprobe binding events depend entirely on reversible affinity for protein pockets, and it may be unrealistic to expect frequent high affinity interactions to emerge from chemical proteomic screens of very small libraries of photo-stereoprobes. We nonetheless believe that the low stoichiometry interactions identified herein, in particular, those showing clear stereoselectivity, are illuminating ligandable pockets on the target proteins. And, our findings further support that the photo-stereoprobes can themselves serve as the basis for higher-throughput screens of these pockets. One potential challenge with converting photo-stereoprobes to fluorescent analogs compatible with NanoBRET screens is identifying a compatible exit vector on the stereoprobe for appending the fluorophore. Depending on how a photo-stereoprobe binds to a specific protein pocket, multiple exit vectors may need to be tested to generate a fluorescently tagged analogue that maintains protein binding.

Our finding that a subset of photo-stereoprobe-protein interactions were not recapitulated with recombinantly expressed proteins and that these cases were enriched for proteins that are parts of large complexes emphasizes the value of performing small molecule binding assays with endogenous proteins in native biological systems, where post-translational modes of regulating protein structure and function are taken into account. This is arguably one of the strongest attributes of chemical proteomics. Nonetheless, we acknowledge that optimizing reversibly binding ligands for proteins that can only be assayed (to date) in endogenous form remains challenging, especially considering the limited throughput of chemical proteomic experiments. Efforts are being made to increase the pace of chemical proteomics^75^, and we expect further advances in this area to facilitate larger library screening in native biological settings.

The photo-stereoprobe-protein interactions identified by chemical proteomics are maps of small-molecule binding, and it is therefore important to consider ways of prioritizing interactions with functional potential. Recent advances in mapping the sites of photoreactive small-molecule binding on a proteome-wide scale should help^76^, as these sites can then be mapped onto established or predicted protein structures to determine their proximity to possible functional pockets (e.g., active sites or at biomolecular interfaces). We have also shown how phenotypic screening can provide a ‘function-first’ assay for photo-stereoprobes. In these screens, the stereoselective activity of hit compounds should facilitate target identification by chemical proteomics, as we have shown for the autophagy modulating stereoprobes engaging the mitochondrial protease CLPP. Considering that genetic disruption of CLPP recapitulated the autophagy modulatory effects of the photo-stereoprobes, we speculate that these compounds act as CLPP inhibitors. Several small molecule and macrocyclic peptide activators of bacterial and mammalian CLPP have been reported^77-79^, as have inhibitors of bacterial CLPP^80,81^, but we are not aware of cell-active inhibitors of mammalian CLPP. How DBK-032A and related photo-stereoprobes may inhibit CLPP will require further investigation and overcoming the current technical challenge of loss of binding of these compounds following cell lysis.

Projecting forward, we anticipate that further efforts to synthesize and evaluate structurally diverse photo-stereoprobes will continue to enrich our understanding of the ligandability of the human proteome. Indeed, the very limited overlap in proteins stereoselectively enriched by the trypoline, azetidine, and pyrrolidine cores examined herein suggests we are far from saturating the full scope of proteins that can be liganded by DOS-inspired stereoprobes in human cells. Our limited SAR analysis of tryptoline photo-stereoprobes related to the autophagy-modulating compounds further points to the value of appendage modifications as an additional source of library diversity. Our current datasets contain many proteins that were enriched by the photo-stereoprobes without stereoselectivity, and we have so far assumed that most of these interactions are non-specific. But, it remains possible that some stereoprobe-proteins interactions lacking stereoselectivity are occurring at defined binding pockets, and a sufficiently deep chemoselective SAR through chemical proteomic analysis of larger sets of photo-stereoprobes may clarify such events. We are also encouraged by the wide range of structural and functional classes of proteins that were stereoselectively engaged by photo-stereoprobes, which extended far beyond classically ligandable proteins like enzymes to include adaptor/scaffolding proteins and transcriptional factors. We acknowledge that understanding the functional potential of such liganding events will require improvements in stereoprobe affinity and selectivity or the use of stereoprobes as target engagement assays to identify additional chemistries binding the newly discovered ligandable pockets. Stereoprobe ligands may also form the basis for constructing heterobifunctional small molecules capable of degrading or redirecting the functions of target proteins^82,83^. Finally, future studies that explore stereoprobe-protein interactions across a broader array of cell types and states may uncover liganding events that occur on, for instance, dynamically regulated proteoforms, which represent a rich and understudied source of protein diversification^84^.

## Methods

### Cell lines/Cell culture

Ramos (ATCC, CRL-1596), 22Rv1 (ATCC, CRL-2505), and HEK293T (ATCC, CRL-3216) were grown and maintained in RPMI-1640 (Gibco) (Ramos and 22Rv1) or high-glucose DMEM (Corning) (HEK293T) supplemented with 10% fetal bovine serum (Omega Scientific), 2 mM GlutaMAX (Gibco), penicillin (100 U/ml), and streptomycin (100 µg/ml) at 37 °C in a humidified 5% CO2 atmosphere. HeLa (ATCC CCL2) cells stably expressing GFP-LC3 were cultured in Dulbecco’s Modified Eagle Medium (DMEM) supplemented with 10% Fetal bovine serum (FBS; Sigma; F0392), 1X Glutamax (Invitrogen; 35050-061), 0.375% sodium bicarbonate at 37 °C in a humidified 5% CO2 atmosphere. Frozen cells were passaged at least twice before usage.

### Cloning

Open reading frames (ORF) of genes of interest were obtained from the Human ORFome V8.1 Library (Dharmacon) or transOMIC (list of sources shown in **Supplementary Table 1**) and cloned into a pRK5-derived plasmid with a C- or N-terminal FLAG tag using the gateway vector conversion system (Invitrogen, cat # 11828029). All gene constructs were verified by DNA sequencing.

For generating N-term-His-ELMO1 construct, N-term strep-His tag was first inserted into phCMV3 vector (Genlantis, cat # P003300) using Q5® Site-Directed Mutagenesis Kit (New England BioLabs, cat # E0554S) with the following primers.

Forward primer: 5’-ggatcgggaggttcagcgtggagccacccgcagttcgagaaaggtggaCACCATCATCATCATCATGGAGGAG-3’

Reverse primer: 3’-accgccagaacctccacctttttcgaactgcgggtggctccaagcgctTCCCATGGTGGCCTCGAG-5’

ELMO1 ORF was then cloned into the phCMV3-derived vector obtained above using the following primers.

Forward primer: 5’-tttGAATTCAGCGGACATCGTCAAGGTG-3’ Reverse primer: 3’-tttGGATCCTCAGTTACAGTCATAGACGAAG-5’

N-term-His-ELMO1C438A construct was generated from N-term-His-ELMO1 construct obtained above using Q5® Site-Directed Mutagenesis Kit (New England BioLabs, cat # E0554S), with the following primers.

Forward primer: 5’-GAGTTGCCTAGTGAGACCGCCAACGACTTCCACCCG-3’

Reverse primer: 3’-CGGGTGGAAGTCGTTGGCGGTCTCACTAGGCAACTC-5’

For generating N-term-NLuc-ELMO1 construct, the PCR products of the Strep-Flag-tag and NLuc (Promega, cat # N1091) were inserted into the pHLmMBP-8 vector (Addgene, cat # 72347)^85^ using restriction enzymes. Strep-Flag-tag and NLuc were cloned using the following primers.

Strep-Flag-tag forward primer: 5’-tttgaattcgccaccatggctagctg-3’

Strep-Flag-tag reverse primer: 3’-tttgcggccgccttgtcatcgtcatccttg-5’

NLuc forward primer: 5’-tttgcggccgcgagcGTCTTCACACTCGAAGATTTC-3’

NLuc reverse primer: 3’-tttgagctcccgcttccgcttccCGCCAGAATGCGTTCG-5’

ELMO1 ORF was then cloned into the pHLmMBP-8-derived vector obtained above using the following primers.

Forward primer: 5’-tttgagctcaggaggaggaGCGGACATCGTCAAGGTG-3’

Reverse primer: 3’-tttacgcgtTCAGTTACAGTCATAGACGAAG-5’

For generating C-term-NLuc-IPO9 construct, IPO9 ORF was cloned into the pFC32A Nluc CMV-neo Flexi^®^ vector (Promega, cat # N1331) using Flexi^®^ Cloning System (Promega, cat # C8640) with the following primers.

Forward primer: 5’-ttttgcgatcgccATGGCTGCAGCTGCAGCTGCTGGAG-3’

Reverse primer: 3’-ttttgtttaAACGATGCCGATGGTCTGCAGCACTCTACGT-5’

### Transient transfection of epitope-tagged proteins of interest in HEK293T cells

HEK293T cells were plated in a 6-well plate (0.45 × 10^6^ cells in 2 mL media/well) and grown for 24 hours. To serum-free DMEM (200 µL), plasmid DNA (1 – 2 µg) and polyethylenimine (PEI, 1 mg/mL) were added using a 1:3 ratio of DNA (μg) to PEI (μL) and incubated for 30 min. Transfection was initiated by adding 200 µL of the transfection mix to each well and incubated for 48 hours.

### Preparation of proteome for gel- and MS-based analysis

Cells were harvested by scraping and/or centrifuging (1,400 × g, 3 min, 4 °C). Harvested cells were washed with cold DPBS (2×), and the pellets were kept frozen at - 80 °C until use. Cell pellets were lysed by sonication (8 pulses × 3, 10% power) in ice-cold PBS using a Branson Ultrasonics Sonifier S-250A cell disruptor to provide whole cell lysates. For the separation of soluble and particulate fractions, the whole cell lysates were centrifuged (100,000 × g, 20 min, 4 °C) to provide soluble (supernatant) and particulate (pellet) fractions. Particulate fractions were resuspended in cold DPBS with sonication (3 pulses ×2, 10% power). Protein concentration of the lysate was determined by DC protein concentration assay (Bio-Rad) followed by absorbance measurements using a CLARIOstar microplate reader (BMG LABTECH). Protein concentration was adjusted to 1 – 1.5 mg/mL (gel-based analysis) or 2 mg/mL (MS-based analysis) with cold DPBS.

### In situ and in vitro labeling of proteome using photo-stereoprobes

*For gel-based analysis (in situ)*: Cells were grown at ∼80 – 100% confluency (22Rv1 and HEK293T cells) or 2 × 10^6^ cells/mL (Ramos cells). Old media was replaced with fresh complete media (containing 10% FBS) pre-mixed with a photo-stereoprobe (1000× stock in DMSO), and incubated at 37 °C for 30 min. For competition experiments with a reversible competitor, cells were incubated with fresh complete media (containing 10% FBS) pre-mixed with a photo-stereoprobe and a reversible competitor together (1000× stock in DMSO respectively) at 37 °C for 30 min. For competition experiments with a covalent competitor, cells were incubated with fresh complete media (containing 10% FBS) pre-mixed with a covalent competitor (1000× stock in DMSO) at 37 °C for 60 min. The media was then replaced with fresh complete media (containing 10% FBS) pre-mixed with a photo-stereoprobe (1000× stock in DMSO), and incubated at 37 °C for 30 min. After probe incubation, cells were exposed to UV light (365 nm) in a UV cross-inker (Stratagene, UV Stratalinker 1800) placed in cold room for 10 min. For no-UV control experiments, cells were placed at 4 °C in the cold room for 10 min under ambient light. Following UV cross-linking, cells were harvested by scraping and/or centrifuging (1,400 × g, 3 min, 4 °C). Harvested cells were washed with cold DPBS (2×), and the pellets were kept frozen at - 80 °C until use. The cell pellets were lysed using the standard cell lysis protocol described above and the protein concentration was adjusted to 1 – 1.5 mg/mL.

*For gel-based analysis (in vitro)*: Cell lysates (50 µL, 1 – 1.5 mg/mL) and a photo-stereoprobe (and a competitor if applicable) in DMSO (50× stock) were incubated at room temperature for 20 min in a clear 96-well plate, followed by exposure to UV light (365 nm) in a UV cross-inker (Stratagene, UV Stratalinker 1800) placed in a cold room for 10 min. For no-UV control experiments, cell lysates and a photo-stereoprobe mixture were placed at 4 °C in the cold room for 10 min under ambient light.

*For MS-based analysis (in situ)*: Cells were grown at ∼80 – 100% confluency in 10 or 15 cm plate (22Rv1 and HeLa cells) or 2 × 10^6^ cells/mL (Ramos cells). Old media was replaced with fresh complete media (containing 10% FBS) pre-mixed with a photo-stereoprobe (1000× stock in DMSO), and incubated at 37 °C for 60 min. For competition experiments with a reversible competitor, cells were incubated with fresh complete media (containing 10% FBS) pre-mixed with a photo-stereoprobe and a reversible competitor together (1000× stock in DMSO respectively) at 37 °C for 60 min. For competition experiments with a covalent competitor, cells were incubated with fresh complete media (containing 10% FBS) pre-mixed with a covalent competitor (1000× stock in DMSO) at 37 °C for 60 min. The media was then replaced with fresh complete media (containing 10% FBS) pre-mixed with a photo-stereoprobe (1000× stock in DMSO), and incubated at 37 °C for 60 min. After probe incubation, cells were exposed to UV light (365 nm) in a UV cross-inker (Stratagene, UV Stratalinker 1800) placed in a cold room for 10 min. For Ramos cells, cells were harvested by centrifuging (1,400 × g, 3 min, 4 °C) and the harvested cells were washed with cold DPBS (3×). For adherent cells, cells were washed with cold DPBS (2×) and scraped with cold DPBS. The cell pellets were kept frozen at - 80 °C until use. The cell pellets were lysed using the standard cell lysis protocol described above and the protein concentration was adjusted to 2 mg/mL.

*For MS-based analysis (in vitro)*: Cell lysates (500 µL, 2 mg/mL) and a photo-stereoprobe (and a competitor if applicable) in DMSO (50× stock) were incubated at room temperature for 20 min in a clear 24-well plate, followed by exposure to UV light (365 nm) in a UV cross-inker (Stratagene, UV Stratalinker 1800) placed in a cold room for 10 min. For no-UV control experiments, cell lysates and a photo-stereoprobe mixture were placed at 4 °C in the cold room for 10 min under ambient light.

### Gel-based analysis of probe-labeled proteins in cells/cell lysates

To each sample (50 μL, 1 – 1.5 mg/mL), 6 μL of freshly prepared ‘click’ mixture (1 μL of 1.25 mM TAMRA-N_3_ in DMSO, 1 μL of freshly prepared 50 mM TCEP in DPBS, 3 μL of 1.7 mM TBTA in t-BuOH:DMSO (4:1, v/v), and 1 μL of 50 mM CuSO_4_ in H_2_O) was added. The reaction mixture was mixed by tapping or gentle vortex and incubated at room temperature for 1 hour before quenching with 4× SDS gel loading buffer (18 μL). Proteins (17 – 25 μg total protein loaded per gel lane) were resolved by SDS-PAGE (10% acrylamide) made in-house and visualized by in-gel fluorescence on a ChemiDoc MP Imaging System (Bio-Rad). The images were processed using Image Lab software (version 6.1.0).

### Western blot analysis of epitope-tagged proteins

After resolving by SDS-PAGE, proteins were transferred to a nitrocellulose membrane (Amersham Protran, cat# 10600011) in Towbin buffer, the membrane was blocked for 1 hour at room temperature with 5% nonfat dry milk (w/v) in Tris-buffered saline with Tween 20 (TBST) and incubated with primary antibodies in the same solution overnight at 4 °C or 1 hour at room temperature. Blots were washed (3×, 5 min, TBST), and incubated with secondary antibodies (IRDye 800CW) in milk for 3 hours at room temperature, washed (3×, 5 min, TBST), rinsed in water and scanned with a ChemiDoc MP Imaging System (Bio-Rad). For samples containing IPO9-NLuc, the blotted proteins were developed using Nano-Glo® In-Gel Detection System (Promega) and scanned with a ChemiDoc MP Imaging System (Bio-Rad).

### MS-based analysis of photo-stereoprobe-labeled proteins

Probe-treated samples were normalized to 2 mg/mL (500 μL) and treated with 55 μL of “click” mix (5μL of 10 mM Biotin-PEG4-N_3_ in DMSO, 10 μL of freshly prepared 50 mM TCEP in DPBS, 30 μL of 1.7 mM TBTA in t-BuOH:DMSO (4:1, v/v), and 10 μL of 50 mM CuSO_4_ in H_2_O) for 1 h at room temperature with gentle vortexing every 20 min. Proteins were precipitated out of solution by the addition of chilled methanol (500 μL), chloroform (200 μL) and water (100 μL), followed by vigorous vortexing and centrifugation at 16,000 × g for 10 min, to create a disk. Without disrupting the protein disk, both top and bottom layers were aspirated, and the protein disk resonicated in 500 µL of methanol washed and centrifuged at 16,000 × g for 10 min. The protein pellets were resuspended in 500 µL of freshly made 6 M urea in DPBS, followed by the addition of 10 µL of 10% SDS and probe-sonicated to clarity. Samples were reduced with 50 μL 1:1 mixture of TCEP (200 mM in DPBS) and K_2_CO_3_ (600 mM in DPBS) at 37 °C for 30 min, followed by alkylation with 70 µL of 400 mM Iodoacetamide at room temperature protected from light for 30 min. Samples were quenched with 130 µL of 10% SDS, transferred to 15 mL tube and the total volume was brought up to 6 mL with DPBS (0.2% final SDS). Washed streptavidin beads (Thermo cat # 20353; 100 µL 50% slurry/sample) was then added and probed labeled proteins were enriched for 1.5 h at room temperature with rotation. After incubation, beads were pelleted (2 min x 800 g) and washed with 0.2% SDS in DPBS (2 x 5 mL), DPBS (1 x 10 mL), milliQ water (1 x 5 mL) then transferred to protein low-bind Eppendorf safe-lock tube. Beads were further washed with milliQ water (1 x 1 mL), and 200 mM EPPS (1x 1 mL). Enriched proteins were digested on-bead overnight with 200 μL of trypsin mix (2 M urea, 1 mM CaCl_2_, 10 μg/mL trypsin, 200 mM EPPS, pH 8.0). Beads were spun down and the supernatant was collected using gel-loading pipet tips. Acetonitrile was added to the supernatant to 30% final volume, followed by 6 μL of 20 µg/µL (in dry acetonitrile) of the corresponding TMT tag (TMT10Plex or TMT16Plex) for 1.5 h at room temperature. TMT labeling was quenched with the addition of hydroxylamine (6 μL 5% solution in H_2_O) and incubated for 15 min at room temperature. Samples were then acidified with 20 μL 100% formic acid, combined and SpeedVac to dryness. Samples were then desalted with Sep-Pak column (Waters).

### HPLC fractionation

Desalted samples were resuspended in 500 μL buffer A (5% acetonitrile, 0.1% formic acid in milliQ water) and fractionated with Agilent HPLC into a 96 deep-well plate containing 20 μL of 20% formic acid to acidify the eluting peptides, as previously reported^86^. The peptides were eluted onto a capillary column (ZORBAX 300Extend-C18, 3.5 μm) and separated at a flow rate of 0.5 mL/min using the following gradient: 100% buffer A from 0-2 min, 0%–13% buffer B from 2-3 min, 13%–42% buffer B from 3-60 min, 42%–100% buffer B from 60-61 min, 100% buffer B from 61-65 min, 100%–0% buffer B from 65-66 min, 100% buffer A from 66-75 min, 0%–13% buffer B from 75-78 min, 13%–80% buffer B from 78-80 min, 80% buffer B from 80-85 min, 100% buffer A from 86-91 min, 0%–13% buffer B from 91-94 min, 13%–80% buffer B from 94-96 min, 80% buffer B from 96-101 min, and 80%–0% buffer B from 101-102 min (buffer A: 10 mM aqueous NH_4_HCO_3_; buffer B: acetonitrile). The plates were evaporated to dryness using SpeedVac and peptides resuspended in 80% acetonitrile, with 0.1% formic acid and combined to a total of 12 fractions (e.g., fraction1= well 1A+ 1B…1H, fraction 2= well 2A+2B….2H) (3x300 μL/column).Samples were SpeedVac to dryness and the resulting 12 fractions were re-suspended in buffer A (5% acetonitrile, 0.1% formic acid) and analyzed by mass spectrometry.

### TMT liquid chromatography-mass-spectrometry (LC-MS) analysis

Samples were analyzed by liquid chromatography tandem mass-spectrometry using an Orbitrap Fusion mass spectrometer (Thermo Scientific) coupled to an UltiMate 3000 Series Rapid Separation LC system and autosampler (Thermo Scientific Dionex), as previously reported^21,87^. The peptides were eluted onto a capillary column (75 μm inner diameter fused silica, packed with C18 (Waters, Acquity BEH C18, 1.7 μm, 25 cm)) or an EASY-Spray HPLC column (Thermo ES902, ES903) using an Acclaim PepMap 100 (Thermo 164535) loading column, and separated at a flow rate of 0.25 μL/min. Data was acquired using an MS3-based TMT method on Orbitrap Fusion or Orbitrap Eclipse Tribrid Mass Spectrometers. Briefly, the scan sequence began with an MS1 master scan (Orbitrap analysis, resolution 120,000, 400−1700 m/z, RF lens 60%, automatic gain control [AGC] target 2E5, maximum injection time 50 ms, centroid mode) with dynamic exclusion enabled (repeat count 1, duration 15 s). The top ten precursors were then selected for MS2/MS3 analysis. MS2 analysis consisted of: quadrupole isolation (isolation window 0.7) of precursor ion followed by collision-induced dissociation (CID) in the ion trap (AGC 1.8E4, normalized collision energy 35%, maximum injection time 120 ms). Following the acquisition of each MS2 spectrum, synchronous precursor selection (SPS) enabled the selection of up to 10 MS2 fragment ions for MS3 analysis. MS3 precursors were fragmented by HCD and analyzed using the Orbitrap (collision energy 55%, AGC 1.5E5, maximum injection time 120 ms, resolution was 50,000). For MS3 analysis, we used charge state–dependent isolation windows. For charge state z = 2, the MS isolation window was set at 1.2; for z = 3-6, the MS isolation window was set at 0.7. Raw files were uploaded to Integrated Proteomics Pipeline (IP2) available at (http://ip2.scripps.edu/ip2/mainMenu.html) and MS2 and MS3 files extracted from the raw files using RAW Converter and searched using the ProLuCID algorithm using a reverse concatenated, non-redundant variant of the Human UniProt database (release 2016-07). Cysteine residues were searched with a static modification for carboxyamidomethylation (+57.02146 Da). N-termini and lysine residues were also searched with a static modification corresponding to the TMT tag (+229.1629 Da for 10plex and +304.2071 Da for 16plex). Peptides were required to be at least 6 amino acids long. ProLuCID data was filtered through DTASelect (version 2.0) to achieve a peptide false-positive rate below 1%. The MS3-based peptide quantification was performed with reporter ion mass tolerance set to 20 ppm with Integrated Proteomics Pipeline (IP2).

### Data processing

Enrichment ratios (probe vs probe) were calculated for each peptide-spectra match by dividing each TMT reporter ion intensity by the sum intensity for all the channels. Peptide-spectra matches were then grouped based on protein ID and, excluding peptides with summed reporter ion intensities < 10,000, coefficient of variation of > 0.5, and < 2 distinct peptides. Replicate channels were grouped across each experiment, and average values were computed for each protein. A variability metric was also computed across replicate channels, which equaled the ratio of median absolute deviation to average and was expressed in percentage. A protein was considered stereoselectively liganded if the average enrichment (of at least 2 biological replicates) by a photo-stereoprobe was > 2.5-fold that of its enantiomer and the variability corresponding to the photo-stereoprobe leading to the highest enrichment did not exceed 50%. The following additional filtering criteria were applied for removing low-quality enrichment events. 1) Enrichment events of an enantiomer pair showing signals > 3-fold lower than the larger signal of their diastereomers were omitted. 2) Enrichment events of an enantiomer pair with their lower signal < 1 were omitted. 3) Enrichment events with diastereo-ratio (min), calculated as follows <= 2 were omitted unless the liganded protein was enantioselectively engaged by both enantiomer pairs of the photo-stereoprobes. Diastereo-ratio (min): The highest signal (across four stereoisomers) divided by the lower signal of its diastereomers.

### Meta-analysis of stereoselectively liganded proteins

The following database/web tools were used for performing meta-analyses of stereoselectively liganded proteins.

*Chemical probe availability*: Probe Miner^88^ (ver. probeminer_datadump_2021-06-20).

*Function class analysis*: Panther Classification System^89^ (ver. PANTHER 18.0). Some of the function classes were combined as follows for data visualization purposes. Metabolite interconversion enzyme + Protein modifying enzyme = Enzymes; Scaffold/adaptor protein + Protein-binding activity modulators = Adaptors, scaffolding, modulators; RNA metabolism protein + DNA metabolism protein = RNA/DNA metabolism proteins; Transporter + Transmembrane signal receptor = Transporter and signal transmembrane receptors; Gene-specific transcriptional regulators + Translational protein + Chromatin-binding or -regulatory protein = Transcriptional regulators, translation and chromatin-binding proteins; Defense immunity protein + Storage protein + Viral or transposable element protein + Cell adhesion molecule + Cell junction protein + Extracellular matrix protein + Intercellular signal molecule + Structural protein = others

*Protein interactome analysis*: BIOPLEX^49^ (ver. BioPlex 3.0 HEK293T cells).

*DepMap analysis*: DepMap^43^ (ver. 22Q2).

*FDA-approved drug target analysis*: DrugBank^90^ (ver. 5.1.10 released on 2023-01-04).

### NanoBRET assay

NanoLuc-tagged protein of interest was recombinantly expressed in HEK293T cells by transient transfection following the standard transfection protocol for epitope-tagged proteins. Cells were harvested, lysed using cold DBPS, and the protein concentration was adjusted to 0.5 mg/mL. 50 µL of the lysate was co-incubated with a BRET tracer (25× stock in DMSO) and a competitor (50× stock in DMSO) at room temperature for 20 min. The mixtures were transferred into a white 384-well plate (Greiner Bio-One) (20 µL per well, duplicate per condition) followed by the addition of Nano-Glo® Substrate (Promega) (5 µL per well) diluted with DBPS (1:100) and mixing by pipetting. After incubating for 2 – 3 min, the filtered luminescence was measured on a CLARIOstar microplate reader (BMG LABTECH) equipped with a 450 nm BP filter (donor) and a 610 nm LP filter (acceptor), using 0.5 s integration time with gain settings of 2,500 – 2,800 and 3,600, respectively. The BRET ratio of each sample was calculated as follows:

BRET = ((Acceptor_sample_/Donor_sample_) – (Acceptor_control_/Donor_control_)) where sample = signal from lysates from the HEK293T cells transfected with NanoLuc-tagged POI, control = signal from lysates from the HEK293T cells (untransfected).

The specific BRET ratio was calculated as follows:

Specific BRET = (BRET_active_ – BRET_inactive_) where active = BRET ratio from a sample treated with active stereoisomer BRET tracer (stereoisomer showing higher binding activity), inactive = inactive stereoisomer BRET tracer (stereoisomer showing lower binding activity).

For the competition assay, specific BRET signals were normalized to DMSO control, and the baseline was corrected to 0 (in the case of the lowest signal value < 0). The curve fitting to determine K_d_ and IC_50_ was performed in GraphPad PRISM v.9.5 (non-linear four-parameter curve fit).

### Computational docking study

Non-covalent docking of WX-02-13 and WX-02-33 was performed on previously published cryo-EM structures of ELMO1:DOCK2:RAC1 complexes in the open ELMO1 conformation (PDB: 6TGC). All calculations were performed on Schrödinger Maestro (MMshare version 4.9.012, release 2020-1, platform Windows-x64). Docking was performed using Glide (v8.6.012) allowing for up to 3 poses per compound, and docking poses were rescored using Prime (v5.9.012, MM-GBSA v3.000). Images were generated using UCSF ChimeraX^91^ (v1.7).

### Generation of GFP-LC3 expressing HeLa cells

HeLa cells stably expressing GFP-LC3 were produced by lentiviral transduction. Briefly, lentiviral plasmids were constructed using a pLX317 backbone in competent Escherichia coli (New England Biolabs, C3050H). Lenti-XTM 293T (Takara, 632810) cells were transfected GFP-LC3 plasmid, VSV-G, and PAX2 plasmids using Lipofectamine3000 (ThermoFisher, L3000001). 24 hours after transfection, cells were given virus production media (DMEM supplemented with 20mM HEPES (Gibco; 15630080) and 30% FBS). The virus was collected the following day. Low passage HeLa cells (ATCC, CCL-2) were transduced with the lentivirus in media containing 8ug/mL polybrene (Sigma Aldrich, TR-1003-G) for 48 hours. HeLa cells were then cultured under selection with 3 μg/mL puromycin (InvivoGen, ant-pr-1) for 48hr and then expanded before storing stocks in liquid nitrogen.

### GFP-LC3 puncta assay

HeLa cells stably expressing GFP-LC3 were plated in 96- or 384-well plates (Perkin Elmer Cell Carrier Ultra; 6057308) at 10000 or 3000 cells per well in supplemented DMEM. The following day, compounds were added by pin transfer using CyBi-Well Vario (Analytik-Jena) into assay plates to the desired concentration. After 4hrs in a 37°C, 5% CO_2_ incubator, cells were washed twice with DPBS and fixed with DPBS containing 4% (v/v) paraformaldehyde (EM Sciences; 100496-496) for 15min at RT. Following fixation, HeLa cells expressing GFP-LC3 were washed twice with PBS, and nuclei were stained with DPBS containing 1 µg/mL of Hoechst-33342 (Sigma; B2261) for 10min at room temperature. After additional PBS wash, cells were imaged at 20x using an automated, high-throughput confocal microscope (Perkin Elmer; Opera Phenix). LC3 (488nm) and DNA (405nm) images were captured from 15 or 5 fields per well and the number of punctate per cell were quantitated using high-content imaging and analysis software (Perkin Elmer; Harmony).

### LC3 flux immunoblot

HeLa cells (ATCC; CCL2) were plated in 3000 µL supplemented DMEM in 6-well TC-treated dishes at 300,000 cells/well. The following day, cells were treated with 10 µM photo-stereoprobes for 4hrs at 37°C. Cells were washed with 500 µL ice-cold DPBS and lifted with a cell scraper. Resuspended cells were transferred to Eppendorf tubes and spun down at 16,000 × g for 10min at 4°C. The cells were lysed with RIPA buffer (Millipore; 20-188) containing protease inhibitor cocktail (Roche; 11-873-580-001) and phosphatase inhibitor (Roche; 4-906-837-001) on ice for 30 min. The lysate was spun down at 16,000 × g for 10 min at 4°C. Prior to SDS-PAGE, protein concentrations were determined by BCA assay (Thermo Fisher; PI23225). Equal amounts of protein samples were loaded and separated by 4-20% bis-tris SDS-PAGE gel (BioRad), transferred onto PVDF membranes (Millipore; Immobilon-FL) and blocked in Odyssey blocking buffer (LI-COR; 927-40000) for 1hr at room temperature. After overnight incubation at 4°C in Odyssey blocking buffer containing primary antibodies [(1:1,000 Anti-LC3B (CST 3868) and 1:20,000 Anti-vinculin (CST 13901S (Rb))], blots were washed with Tris-buffered saline containing 0.1% Tween 20 (TBS-T) and incubated for 1hr at room temperature in secondary antibody solution [1:1 dilution of LI-COR blocking buffer/TBS-T supplemented with 0.02% SDS; 1:10,000 IR-Dye 680LT goat anti-Rb (LI-COR; 925-68021); and 1:10,000 IR-Dye 800CW goat anti-Mo (LI-COR; 925-32210)]. Blots were imaged and quantitated using the LI-COR Odyssey infrared imaging system.

### siRNA-mediated gene knockdown

siRNA-mediated knockdown was performed based on Ambion’s Silencer select RNAi transfection protocol. GFP-LC3 expressing HeLa cells were lifted and seeded in 96-well plates (7000 cells per well) prior to transfection. 1.5 µL Lipofectamine RNAiMAX reagent (Thermofisher: 13778075) was diluted in 25 µL OptiMEM (Thermofisher: 31985070) and combined with 5 pmol of siRNA in 25 µL OptiMEM media. The resulting solution was incubated at room temperature for 10 min, and 10 µL of this solution was added to each well containing 90 µL media. The media was changed after 8 hours, and the cells were subjected to compound treatment 72 hours post-transfection. A list of siRNAs used is listed in **Supplementary Table 1**.

### Ribonucleoprotein (RNP) nucleofection-mediated CRISPR Knockout

1 µL of 63 µM Alt-R™ S.p.Cas9 V3 (IDT: 10007807) was diluted with 18 µL of SE Cell Line Nucleofector® Solution (Lonza V4XC-1032), followed by the addition of 1.8 µL of 100 µM sgRNA (3:1 sgRNA:Cas9). The resulting solution was incubated at room temperature for 10 min. 20 µL of this solution was then used to resuspend 200,000 GFP-LC3 expressing HeLa cells in an Eppendorf tube before being transferred to Lonza Nucleocuvette strips. Nucleofection was performed using program CN-114 on a Lonza-4D Nucleofector X unit. After transfection, 180 µL of pre-warmed DMEM media was added and the cells were allowed to recover for 20 min. 20 or 50 µL of the suspension to 96 or 24-well plates and the cells were treated with compounds or directly harvested 72 hours after nucleofection. A list of sgRNA used are listed in **Supplementary Table 1**.

### Measurement of cell viability with CellTiter-Glo

Cell viability was measured using CellTiter-Glo (Promega)’s protocol. Briefly, 100 µL of CellTiter-Glo was added to transfected or nucleofected cells in 100 µL media in a 96-well plate. Each plate was shaken and incubated at room temperature for 10 min. Cell viability was determined based on real luminescence units following an ATP chemi-luminescent reaction. ATP concentrations were used as a proxy for cell viability, which is expressed as real luminescence units (RLUs) measured on a BMG Labtech PHERAstar FSX.

### Plane of Best Fit (PBF) analysis of probe scaffolds

Plane of Best Fit (PBF)^38^ scores were compared for FFF probes^24^ (14 compounds), fragment enantioprobes^25^ (16 compounds = 8 pairs of enantiomers), elaborated enantioprobes (18 compounds = 9 pairs of enantiomers), and DOS photo-stereoprobes (12 compounds = 6 pairs of enantiomers) (see **Supplementary Table 2** for full list). Prior to performing PBF score calculations, (1) photo-affinity tags were replaced in chemical structures with propanoyl groups, (2) because enantiomers have identical PBF scores, only one compound was selected from each pair of enantiomers. For each considered compound, 200 conformers were generated using RDKit ETKDG version 3^92^, followed by structure refinement using Merck molecular force field (MMFF). After performing RMS pruning (RMSD cutoff = 1.0), PBF scores were calculated for the top 3 (maximum) most stable conformers using the built-in function in RDKit. Note: for some of the compounds, the numbers of the conformers that met RMSD >= 1.0 were less than 3.

### Data and Code availability

The mass spectrometry proteomics data have been deposited to the ProteomeXchange Consortium via the PRIDE^93^ partner repository with the dataset identifier PXD050096. Processed proteomic data are provided in **Supplementary Dataset 1**. All biological assay data are included in **Supplementary Table 2**. All other data that support the findings of this study and codes are available from the corresponding authors upon reasonable request.

## Supporting information

Supplementary Chemistry Information

## Acknowledgments

This work was supported by the NIH (U19 AI142784, R35 CA231991), Bristol Myers Squibb, JSPS overseas research fellowship (D.O.), HHMI Hanna H Gray Fellowship (E.N., GT15176), and Jane Coffin Childs Memorial Fellowship (K.E.D.). The authors thank Dr. Xuedong Liu and Dr. Bing Chen (WuXi AppTec) for small-molecule synthesis.

## Author contributions

D.O., B.M., and B.F.C. conceived the study. D.O. generated proteomic data. D.O., B.M., and B.F.C. performed analysis of proteomic data and wrote the manuscript. D.O. confirmed protein-stereoprobe interactions by gel-based profiling. D.O. performed nanoBRET assay. D.O., and D.B.K. synthesized compounds. D.O., D.B.K., B.M., and B.F.C. supervised compound characterization. Z.Y.T., J.L., and K.L.C. performed phenotypic screening and biological characterizations. Additional resources to the study were contributed by S.J.W., T.C., H.L., K.E.D, and E.N. All authors edited and approved the manuscript. B.F.C., R.J.X, and S.L.S. supervised the study.

---

**Figure S1.**
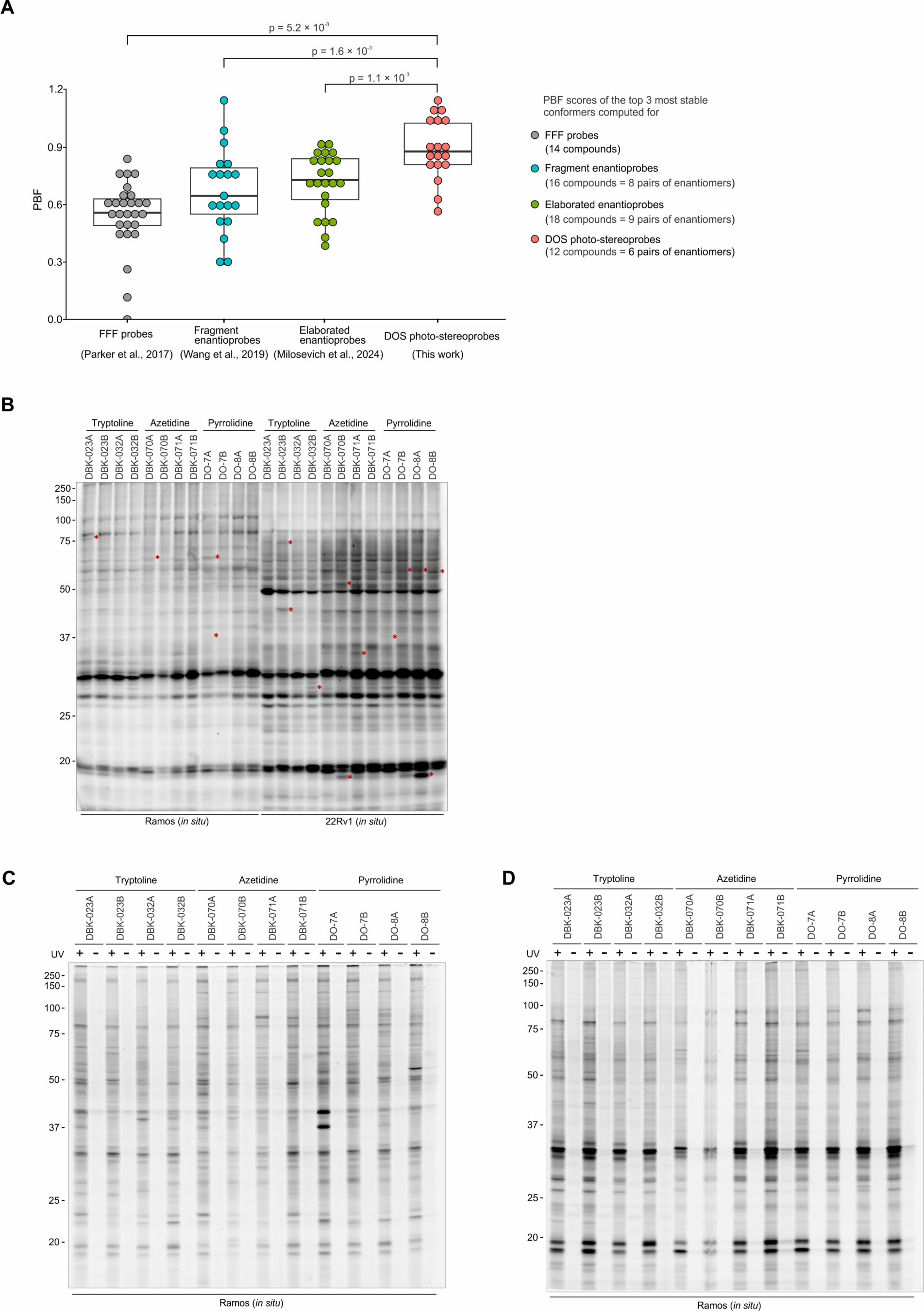
Photo-stereoprobes for mapping reversible small molecule-protein interactions in human cells, related to. Figure 1. (**A**) Plane of Best Fit (PBF) scores were compared for FFF probes (14 compounds), fragment enantioprobes (16 compounds = 8 pairs of enantiomers), elaborated enantioprobes (18 compounds = 9 pairs of enantiomers) and DOS photo-stereoprobes (12 compounds = 6 pairs of enantiomers). Prior to performing PBF score calculations, (1) photo-affinity tags were replaced in chemical structures with propanoyl groups, (2) because enantiomers have identical PBF scores, only one compound was selected from each pair of enantiomers. For each considered compound, 200 conformers were generated, and PBF scores were computed for the top 3 most stable conformers for each probe scaffold. Conformer generation and PBF score calculations were performed using RDKit. Statistical analysis was performed by two-sided Student’s t-test relative to the FFF probes group. (**B**) Gel profiling of stereoprobe-protein interactions in the particulate fraction of human cells. Ramos and 22Rv1 cells were incubated with photo-stereoprobes (20 µM) for 30 min followed by UV cross-linking for 10 min. Stereoprobe-modified proteins in the particulate fraction were conjugated to a tetramethylrhodamine-azide (TAMRA-N_3_) reporter tag by CuAAC and analyzed by SDS-PAGE and in-gel fluorescence scanning. Red asterisks mark proteins showing stereoselective labeling by stereoprobes. (**C**–**D**) UV light-dependence of photostereoprobe-protein interactions in human cells. Ramos cells were incubated with photo-stereoprobes for 30 min *in situ*. Cells were harvested before or after 10 min UV-irradiation. Soluble (**C**) and particulate (**D**) fractions of the harvested cells were subjected to CuAAC reaction with TAMRA-N_3_. The samples were then analyzed by SDS-PAGE and in-gel fluorescence scanning.

**Figure S2.**
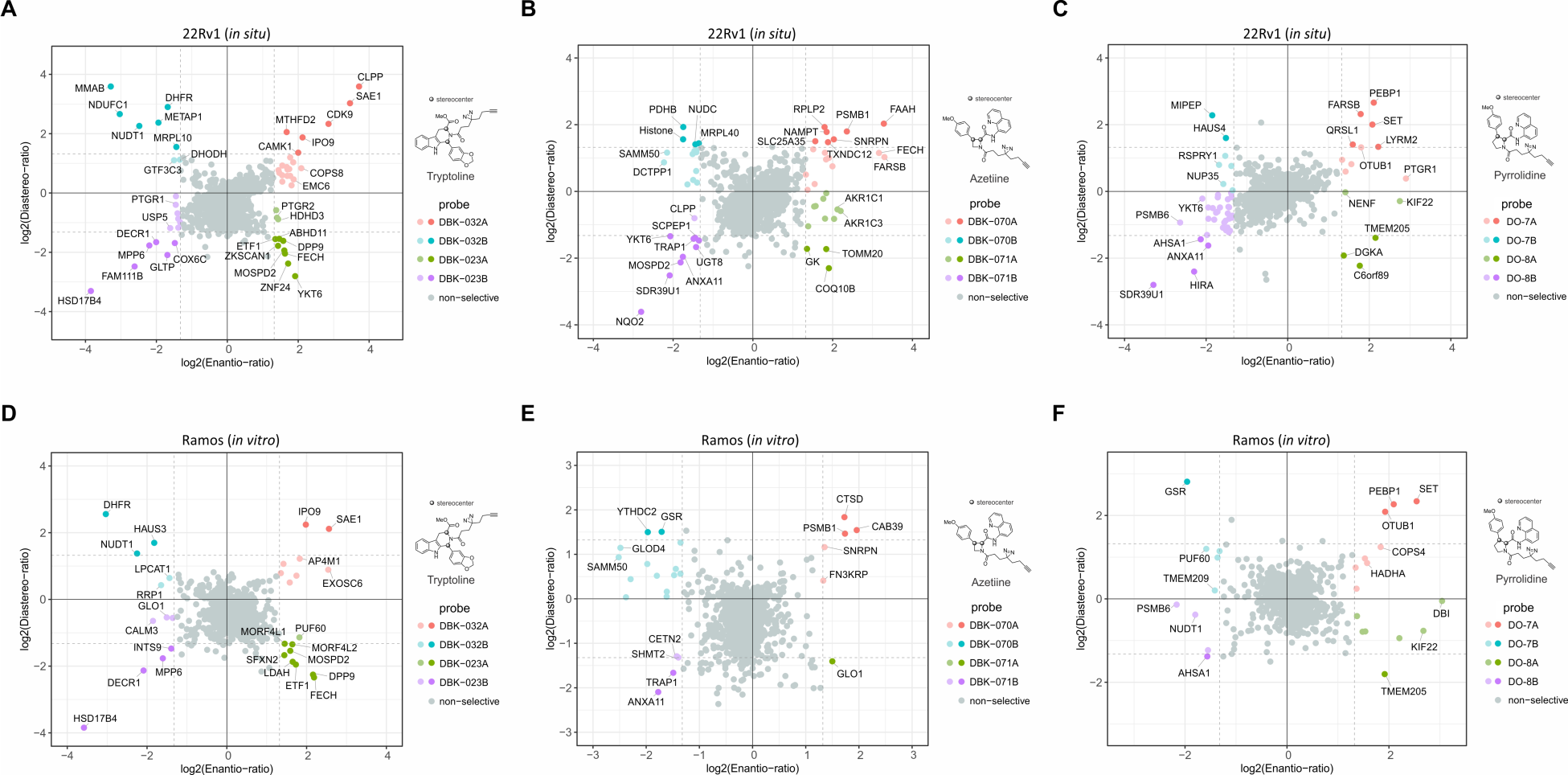
Mass spectrometry (MS)-based proteomic profiling of photo-stereoprobe-protein interactions in human cells, related to. Figure 2. (**A**–**C**) Quadrant plots displaying stereoselectively liganded proteins in 22Rv1 cells for (**A**) tryptoline, (**B**) azetidine, and (**C**) pyrrolidine photo-stereoprobe sets. (**D**–**F**) Quadrant plots displaying stereoselectively liganded proteins in Ramos cell lysates for (**D**) tryptoline, (**E**) azetidine, and (**F**) pyrrolidine photo-stereoprobe sets. Enantio-ratio (x-axis) is the fold-enrichment ratio for the stereoisomer showing maximum engagement of a protein (probe^max^) over its enantiomer. Diastereo-ratio (y-axis) is the fold-enrichment ratio of probe^max^ over the diastereomer with higher relative engagement. Proteins that show an enantio-ratio >= 2.5 are colored as follows. Dark color: proteins that show enantio-ratio >= 2.5 and diastereo-ratio >= 2.5. Light color: proteins that show enantio-ratio >= 2.5 and diastereo-ratio < 2.5. Data represents mean values from at least two independent experiments.

**Figure S3.**
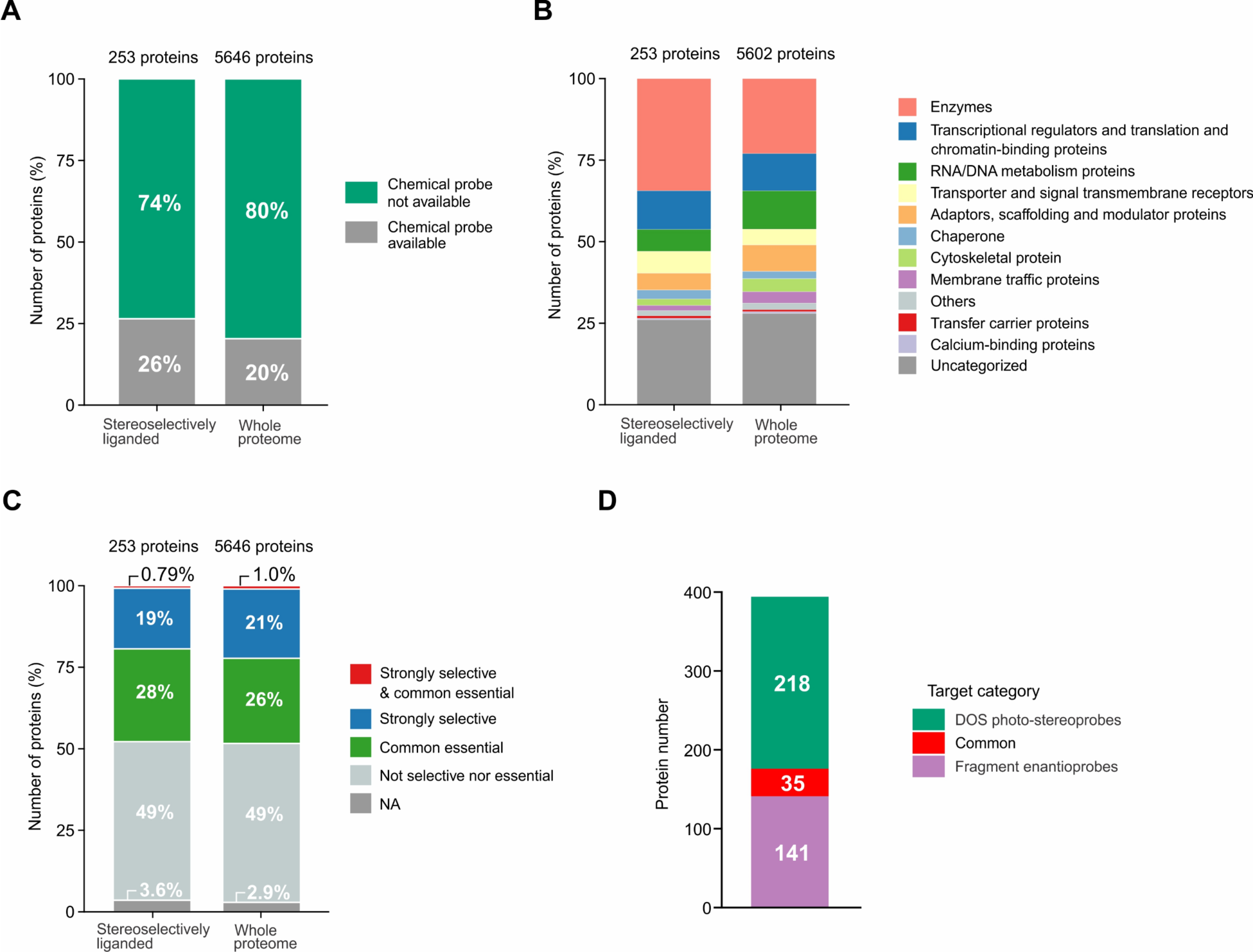
Additional features of stereoselectively liganded proteins by photo-stereoprobes compared to all proteins quantified in MS-based proteomic analysis of Ramos cells, related to. Figure 3. Bar graphs showing the proportion of stereoselectively liganded proteins versus all quantified proteins in Ramos cells classified by (**A**) the availability of chemical probes as assigned by the Probe Miner database, (**B**) protein functional classes as assigned by the Panther database, (**C**) essentiality as assigned in the Cancer Dependency Map. (**D**) Bar graphs showing proportion of proteins stereoselectively liganded by DOS photo-stereoprobes, fragment enantioprobes, or both sets of photo-reactive probes. Data for fragment enantioprobes derived from ref 25.

**Figure S4.**
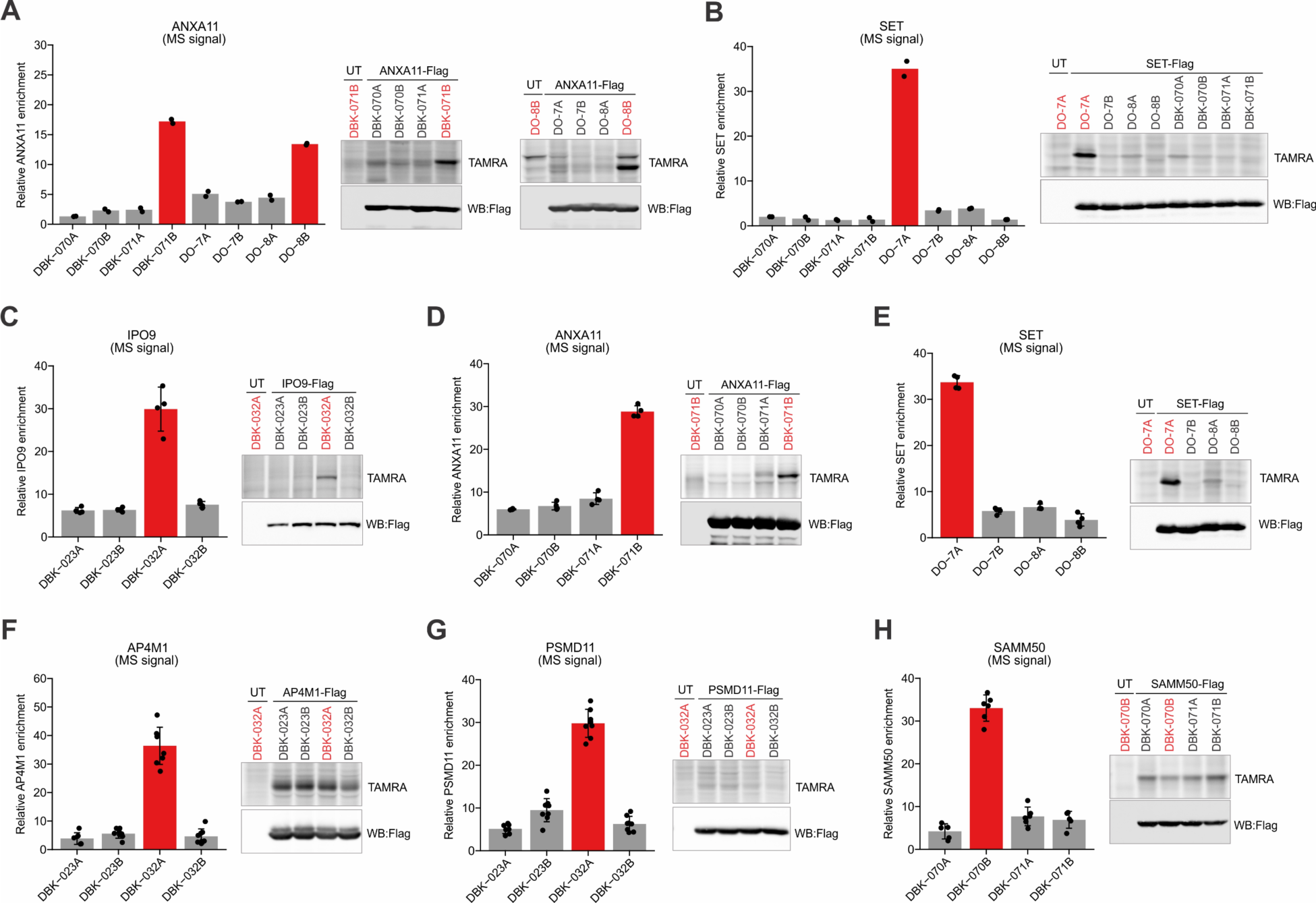
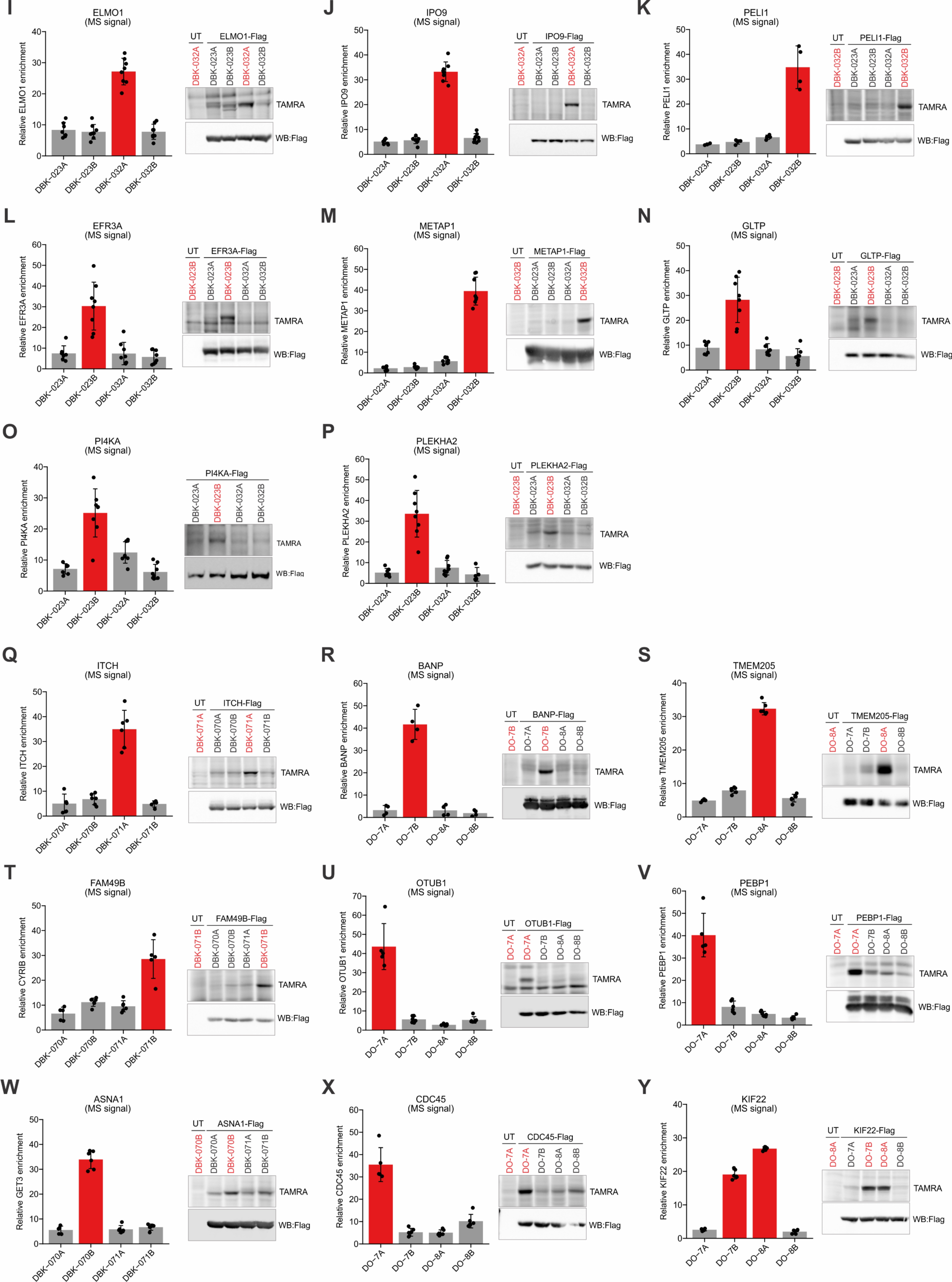
Additional characterization of proteins stereoselectively liganded by photo-stereoprobes, related to. **Figure 4**. (**A**) ANXA11 is a shared stereoselective target of the azetidine and pyrrolidine photo-stereoprobes. (**B**) SET is stereoseletively liganded by pyrrolidine photo-stereoprobe DO-7A, but not the stereochemically matched azetidine photo-stereoprobe DBK-070A. (**C**–**E**) Confirmation of *in vitro* stereoselective engagement of IPO9 (**C**), ANXA11 (**D**), and SET (**E**) by indicated photo-stereoprobes. (**F**–**H**) Example of proteins showing endogenous, but not recombinant stereoselective engagement with photo-stereoprobes. Assessment of photo-stereoprobe-protein interactions are shown for AP4M1 (**F**), PSMD11 (**G**) and SAMM50 (**H**). (**I**–**P**) Additional characterization of proteins stereoselectively liganded by tryptoline photo-stereoprobes. Stereoselective engagement of ELMO1 (**I**), IPO9 (**J**), PELI1 (**K**), EFR3A (**L**), METAP1 (**M**), GLTP (**N**), PI4KA (**O**) and PLEKHA2 (**P**) by the indicated photo-stereoprobes were confirmed. (**Q**–**Y**) Additional characterization of proteins stereoselectively liganded by azetidine and pyrrolidine photo-stereoprobes. Stereoselective engagement of ITCH (**Q**), BANP (**R**), TMEM205 (**S**), FAM49B (**T**), OTUB1 (**U**), PEBP1 (**V**), ASNA1 (**W**), CDC45 (**X**) and (**Y**) KIF22 by indicated photo-stereoprobes were confirmed. For (**A**–**Y**), Left bar graph: relative protein enrichment profiles in Ramos cells (**A**, **B, F**–**Y**) or Ramos cell lysates (**C**–**E**) treated with indicated photo-stereoprobes (20 µM) determined by MS-ABPP. For (**A**, **B**), bar graph data represents mean values of duplicates per group. For (**C**–**Y**), bar graph data represents mean values ± SD for four independent experiments. Right gel image: validation of stereoselective photo-stereoprobe-protein interactions with recombinantly expressed proteins by gel-ABPP. Indicated proteins were recombinantly expressed with Flag epitope tags by transient transfection in HEK293T cells, and transfected cells (**A**, **B, F**–**Y**) and/or the lysates of transfected cells (**C**–**E**) were treated with the indicated photo-stereoprobes (at 20 µM except indicated otherwise) followed by UV cross-linking for 10 min. Probe-modified proteins were conjugated to TAMRA-N_3_ with CuAAC reaction. The samples were then analyzed by SDS-PAGE and in-gel fluorescence scanning.

**Figure S5.**
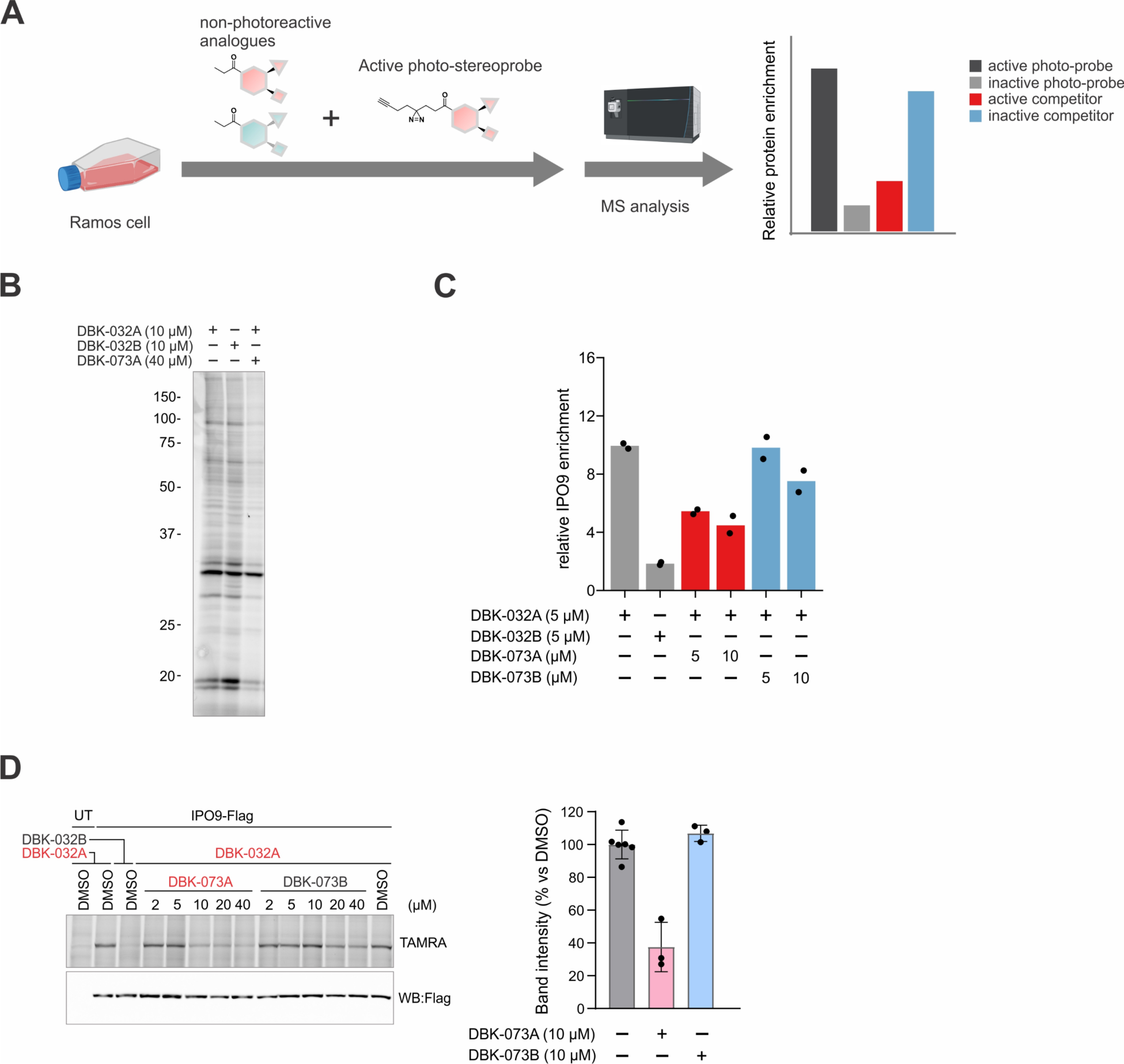
Assessing the stoichiometry of stereoprobe-protein interactions in cells, related to. Figure 5. (**A**) Workflow for the MS-based analysis of the stoichiometry of stereoprobe-protein interactions based on comparative quantification of protein enrichments from Ramos cells co-incubated with DMSO or 1–2X (5–10 µM) non-photoreactive competitor stereoprobes and photo-stereoprobes (5 µM) of the same stereochemistry for 60 min. (**B**) Global suppression of *in situ* photo-stereoprobe-protein interactions by high concentration (40 µM) of non-photoreactive competitor. Ramos cells were treated with DBK-032A (10 µM), DBK-032B (10 µM), or DBK-032A (10 µM) with excess DBK-073A (40 µM) for 20 min followed by UV light-induced photocrosslinking, conjugation of photo-stereoprobe-modified proteins to TAMRA-N_3_ with CuAAC reaction, and analysis by SDS-PAGE and in-gel fluorescence scanning. (**C**) Concentration-dependent stereoselective blockade of DBK-032A enrichment of IPO9 by DBK-073A in Ramos cell as determined by following the workflow shown in panel (**A**). Data represents mean values for two independent experiments per group. (**D**) Concentration-dependent stereoselective blockade of DBK-032A interactions with recombinant Flag-tagged IPO9 in transfected HEK293T cells as determined by gel-based analysis (left) and quantification of data from cells treated with 10 µM of DBK-073A or DKB-073B (right). Data represents mean values ± SD for three independent experiments per group.

**Figure S6.**
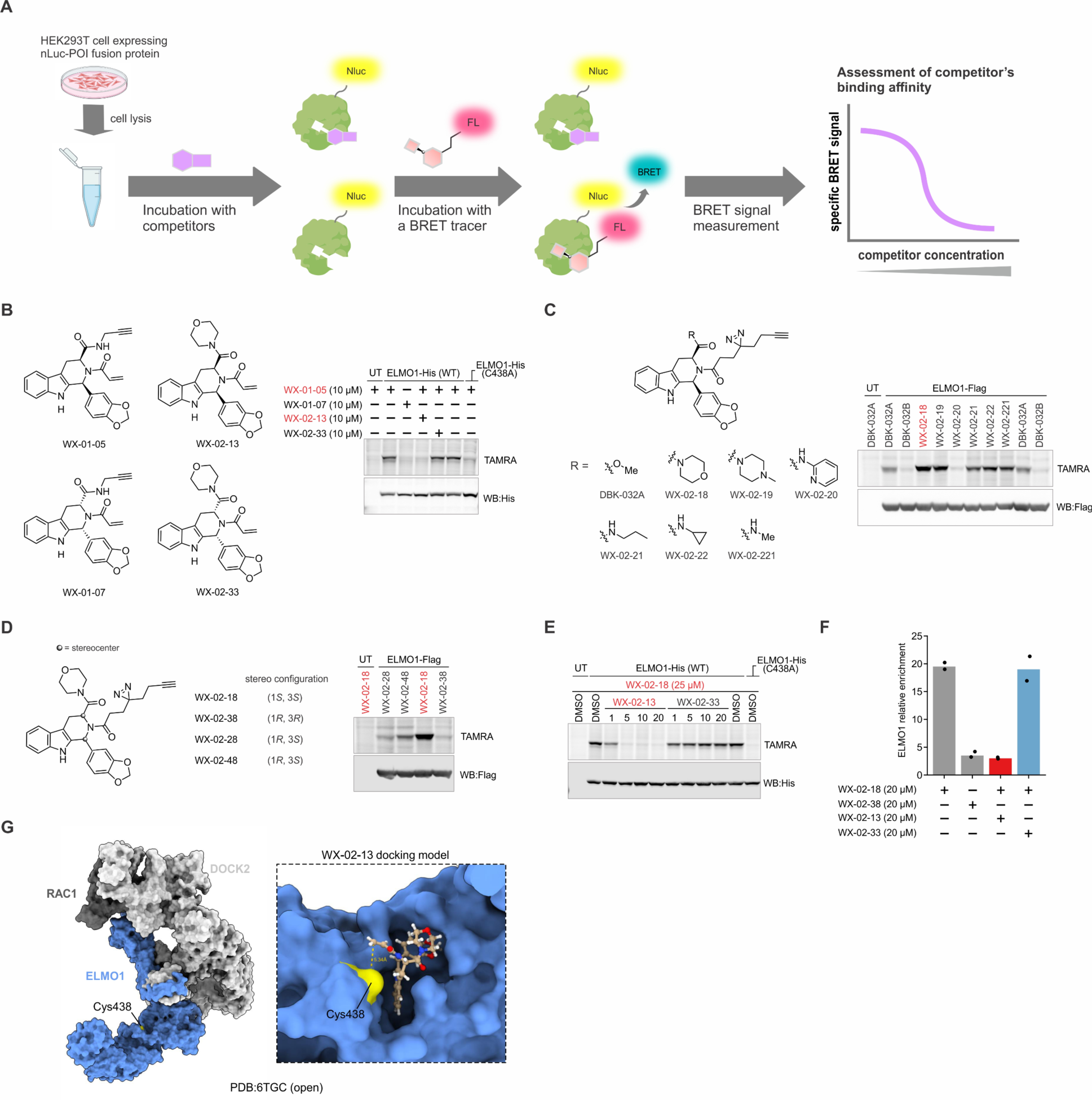
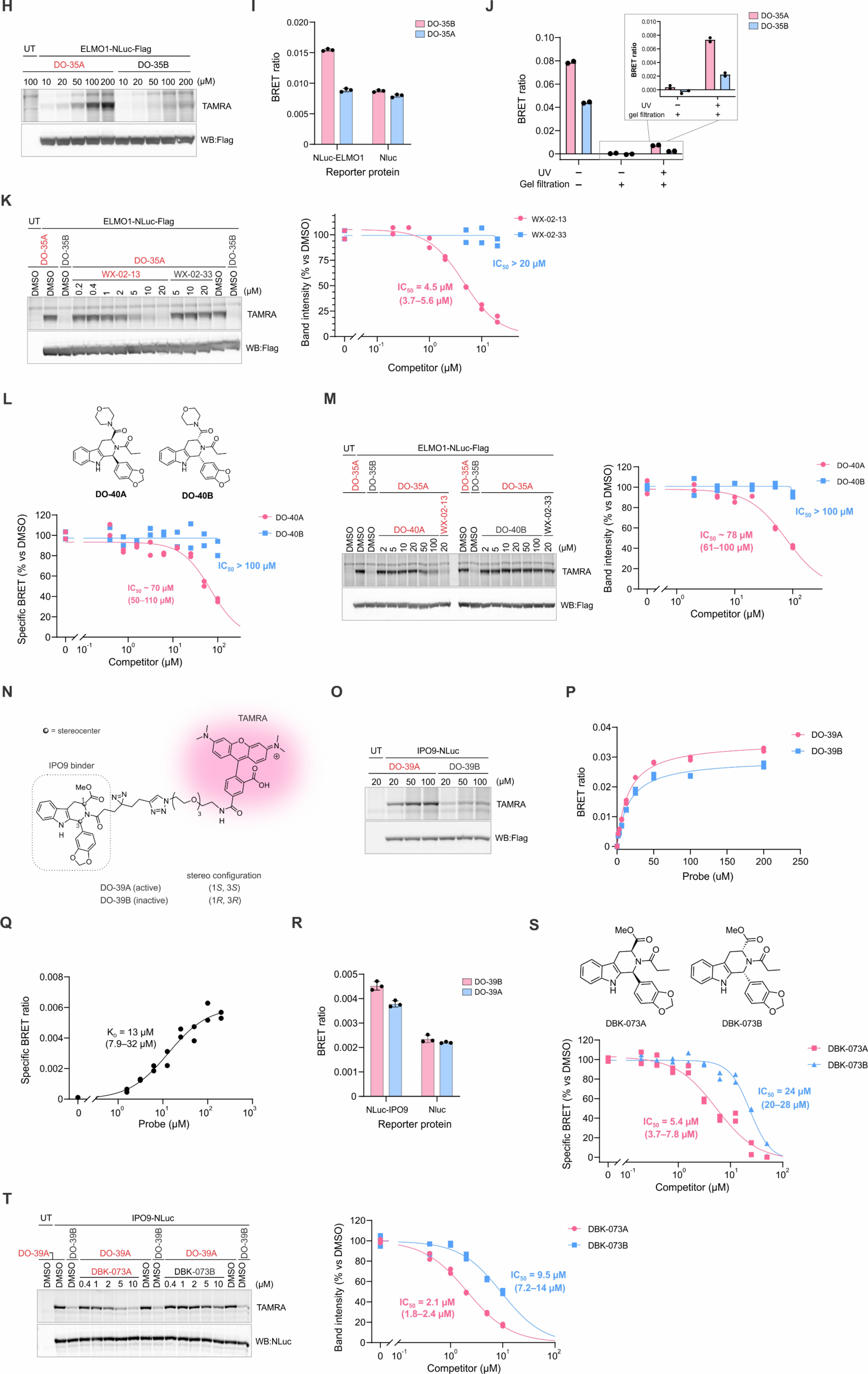
Converting photo-stereoprobe-protein interactions into a high-throughput screening-compatible nanoBRET assay, related to. Figure 6. (**A**) Schematic workflow for competitive nanoBRET assay to assess the binding affinity of a small molecule competitor with a POI. (**B**–**F**) SAR analysis of stereoprobe interactions with ELMO1. (**B**) Structures of electrophilic tryptoline acrylamide ligands for ELMO1 (left) and gel-ABPP data showing stereoselective engagement by alkyne WX-01-05 and stereoselective blockade of this engagement by WX-02-13 (right). (**C**) Structures of DBK-032A analogues and analysis of their engagement with recombinant ELMO1 as measured by gel profiling. (**D**) Confirmation of stereoselective engagement of recombinant ELMO1 by morpholine amide WX-02-18. (**E, F**) WX-02-13, but not the inactive enantiomer WX-02-33 blocked WX-02-18 engagement of recombinant (**E**) or endogenous (**F**) ELMO1 as determined by gel profiling and MS-based proteomics, respectively. WX-02-18 did not interact with a C438A mutant of ELMO1 (**E**). For (**C**) and (**D**), HEK293T cells transiently expressing Flag-ELMO1 were incubated with photo-stereoprobes (20 µM) for 30 min followed by UV cross-linking for 10 min. For (**B**) and (**E**), HEK293T cells transiently expressing His-ELMO1 or His-ELMO1 C438A were incubated with WX-02-13 or WX-02-33 (20 µM) for 60 min. Cells were then incubated with WX-01-05 (10 µM) for 60 min (**B**) or WX-02-18 (25 µM) for 30 min followed by UV cross-linking for 10 min (**C**). For (**B**–**E**), probe-modified proteins were conjugated to TAMRA-N_3_ by CuAAC and then analyzed by SDS-PAGE and in-gel fluorescence scanning. For (**F**), Ramos cells were treated with WX-02-13 or WX-02-33 (20 µM, 1 h) followed by WX-02-18 or WX-02-38 (20 µM, 40 min) and UV cross-linking for 10 min. Relative protein enrichment was quantified following the workflow described in Figure 2. Data represents mean values of duplicates per group. (**G**) Computational docking study revealed that WX-02-13 but not WX-02-33 adopted a binding pose where the acrylamide terminal carbon was positioned in proximity to the ELMO1_C438 sulfhydryl (inset), consistent with the reactivity observed by cysteine-directed ABPP. Non-covalent docking of WX-02-13 and WX-02-33 was performed on previously published cryo-EM structures of ELMO1:DOCK2:RAC1 complexes in open (PDB: 6TGC) conformations. (**H**) Gel assay confirming stereoselective engagement of ELMO1 by DO-35A. Lysate of HEK293T cells recombinantly expressing ELMO1-NLuc-Flag was incubated with varied concentrations of DO-35A or DO-35B for 20 min at room temperature followed by UV cross-linking (10 min). The samples were then analyzed by SDS-PAGE and in-gel fluorescence scanning. (**I**) Comparison of BRET signals for NLuc-ELMO1 and NLuc only control. HEK293T lysate containing recombinant NLuc-ELMO1-Flag or NLuc were incubated with DO-35A or DO-35B (100 µM), and BRET signals measured after 20 min incubation. Data represents mean values ± SD for three independent experiments per group. (**J**) Comparison of BRET signals with or without gel-filtration and UV crosslinking for DO-35A/B. HEK293T lysate containing recombinant NLuc-ELMO1-Flag was incubated with DO-35A or DO-35B (100 µM). The mixture was then gel-filtered with or without prior exposure to UV light (10 min). Data represents mean values of two independent experiments per group. (**K**) WX-02-13, but not WX-02-33, produced concentration-dependent blockade of ELMO1 engagement by DO-35A as determined by gel assay. Left, representative gel; right, quantification of data from two independent experiments. (**L**) Structures of non-electrophilic competitor ligands DO-40A and DO-40B (left) and concentration-dependent blockade of BRET signal for DO-35A-ELMO1 interactions by DO-40A, but not DO-40B (right). NLuc-ELMO1-Flag expressed HEK293T cell lysates was incubated with DO-35A (100 µM) varied concentrations of DO-40A or B, and BRET signal was measured after 20 min incubation. (**M**) DO-40A, but not DO-40B, produced concentration-dependent blockade of ELMO1 engagement by DO-35A as determined by gel analysis. For (**K**–**M**), the experiments were performed in duplicates. (**N**–**T**) Development of nanoBRET assay for IPO9. (**N**) Structures of BRET tracer DO-39A and inactive enantiomer control DO-39B for IPO9. (**O**) Gel assay confirming stereoselective engagement of IPO9 by DO-39A. Lysate of HEK293T cells recombinantly expressing IPO9-NLuc was incubated with varied concentrations of DO-39A or DO-39B for 20 min at room temperature followed by UV cross-linking (10 min). The samples were then analyzed by SDS-PAGE and in-gel fluorescence scanning. (**P**) Concentration-dependent and stereoselective BRET signal for DO-39A-IPO9-NLuc interactions. Data are from two independent experiments. (**Q**) Specific BRET signal as a subtracted component of BRET signals for DO-39A and DO-39B generated in (**P**). (**R**) Comparison of BRET signal for NLuc-IPO9 and NLuc only control. HEK293T lysate containing recombinant NLuc-IPO9-Flag or NLuc were incubated with DO39-A or B (25 µM), and BRET signal was measured after 20 min incubation. Data represents mean values ± SD for three independent experiments per group. (**S**) Structures of competitor compounds DBK-073A and DBK-073B (left) and concentration-dependent blockade of BRET signal for DO-39A-NLuc-IPO9-Flag interactions by these compounds (right). NLuc-IPO9-Flag expressed HEK293T cell lysates was incubated with DO-39A (25 µM) and varied concentrations of DBK-073A or DBK-073B, after which BRET signal was measured as described in (**P**). Data are from two independent experiments. (**T**) DBK-073A, but not DBK-073B, produced concentration-dependent blockade of IPO9 engagement by DO-39A as determined by gel analysis. Data are from two independent experiments.

**Figure S7.**
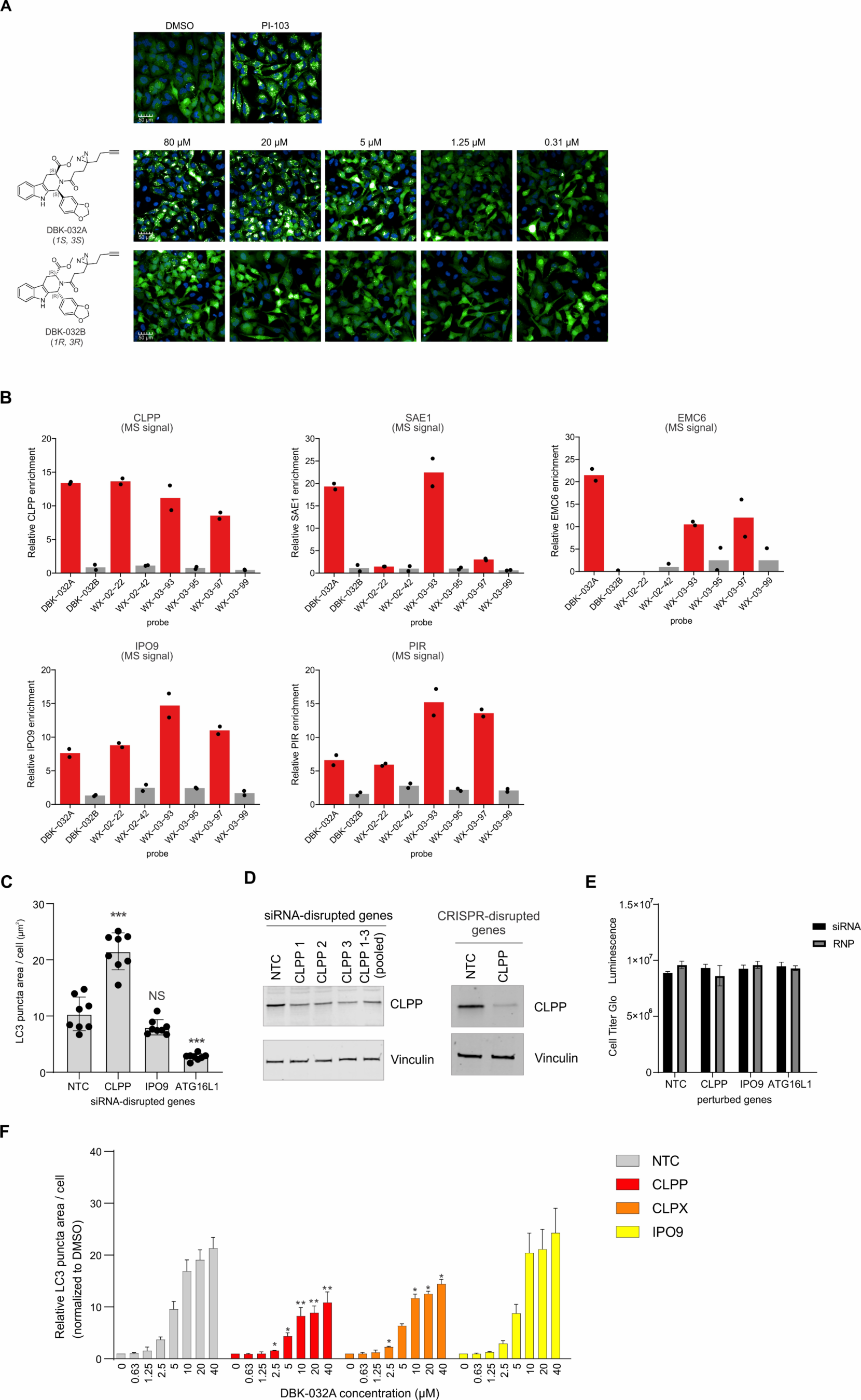
Integrated phenotypic screening and chemical proteomics identifies photo-stereoprobes that modulate autophagy by engaging CLPP, Related to. Figure 7. (**A**) Stereoselective induction of LC3 puncta formation by DBK-032A in HeLa cells. HeLa cells stably expressing LC3-GFP were treated with the indicated concentrations of DBK-032A or DBK-032B for 4 h, after which LC3 puncta formation was measured by fluorescence microscopy. The mTOR inhibitor PI-103 (5 µM) was included as a positive control. Data are from a single experiment representative of at least 3 independent experiments. (**B**) Enrichment profiles for candidate targets relevant to the LC3 puncta induction phenotype in HeLa cells treated with indicated photo-stereoprobes. HeLa cells stably expressing LC3-GFP were treated with the indicated active (LC3 puncta-inducing; red bars) and inactive enantiomeric (gray bars) photo-stereoprobes (20 µM, 1 h), followed by quantitative MS-based proteomic analysis using the workflow described in Figure 2. Data represents mean values of two independent experiments per group. (**C**–**E**) Genetic perturbation of candidate photo-stereoprobe targets responsible for LC3 puncta formation induced by DBK-032A. (**C**) siRNA-mediated knockdown of CLPP, but not IPO9, promoted basal LC3 puncta formation in HeLa cells. (**D**) Immunoblot confirming genetic knockdown/knockout of CLPP. (**E**) siRNA-mediated knockdown and CRISPR-mediated knockout of CLPP or IPO9 did not affect cell viability. (**F**) siRNA-mediated knockdown of CLPP or CLPX, but not IPO9, abrogated LC3 puncta formation induced by DBK-032A. For (**C**), data represent average values ± SD (n = 7 per group). For (**E**), data represent average values ± SD (n = 3–4 per group). For (**F**), data represent average values ± SD (n = 3 per group). NTC: non-targeting control. ATG16L1 (essential autophagy gene) was included as a negative control. *P < 0.05; **P < 0.01; ***P < 0.001 (two-sided Student’s t-test). Two-sided Student’s t-test was performed relative to NTC control group (**C**) or NTC control group with corresponding DBK-032A dose (**F**).

