## Supplementary Chemistry Information for "Chemical tools to expand the ligandable proteome: diversity-oriented synthesis-based photoreactive stereoprobes"

#### Synthetic Chemistry Procedures and Analytical Data

##### General Information

All chemical reagents were purchased from commercial suppliers and were used without further purification. Flash chromatography was performed with 20-40 μm silica gel (60-Å mesh) on a Teledyne Isco Combiflash Rf or a Biotage Isolera Prime, alternatively in a glass column using SiliaFlash® F60 silica gel (40–63 μm, 60 Å). Merck silica gel TLC plates (0.25 mm, 60 F254) were used to monitor reactions. Preparative high-pressure liquid chromatography (prep-HPLC) was performed on a Gilson GX-281 instrument equipped with a Phenomenex Gemini C18 column (75 mm × 30 mm × 3 μM) or Waters Xbridge (150 mm × 25 mm × 5 μM) unless indicated otherwise, eluting with a mixture of acetonitrile and a buffered aqueous phase. Preparative thin layer chromatography (prep-TLC) was performed on Analtech Preparative Uniplates (20x20cm, UV/Silica G, 15 μm particle size, 500-2000 μM thickness). NMR spectra were recorded at 300K on Bruker DRX-600 spectrometer at 600 (1H) MHz or Bruker Avance III 400, Avance III HD 400, Avance Neo 400 spectrometers 400 (1H) MHz. Chemical shifts are recorded in ppm relative to tetramethylsilane (TMS) with peaks being reported as follows: chemical shift, multiplicity (s = singlet, brs = broad singlet, d = doublet, t = triplet, q = quartet, m = multiplet), coupling constant (Hz). High-resolution mass spectra (HRMS) was performed on a 6230 TOF LC/MS with a Dual AJS EI source by injecting 5 μM compounds in MeOH with 20% solvent A (0.1% formic acid in water) and 80% solvent B (0.1% formic acid in acetonitrile).

#### Synthesis of tryptoline photo-stereoprobes

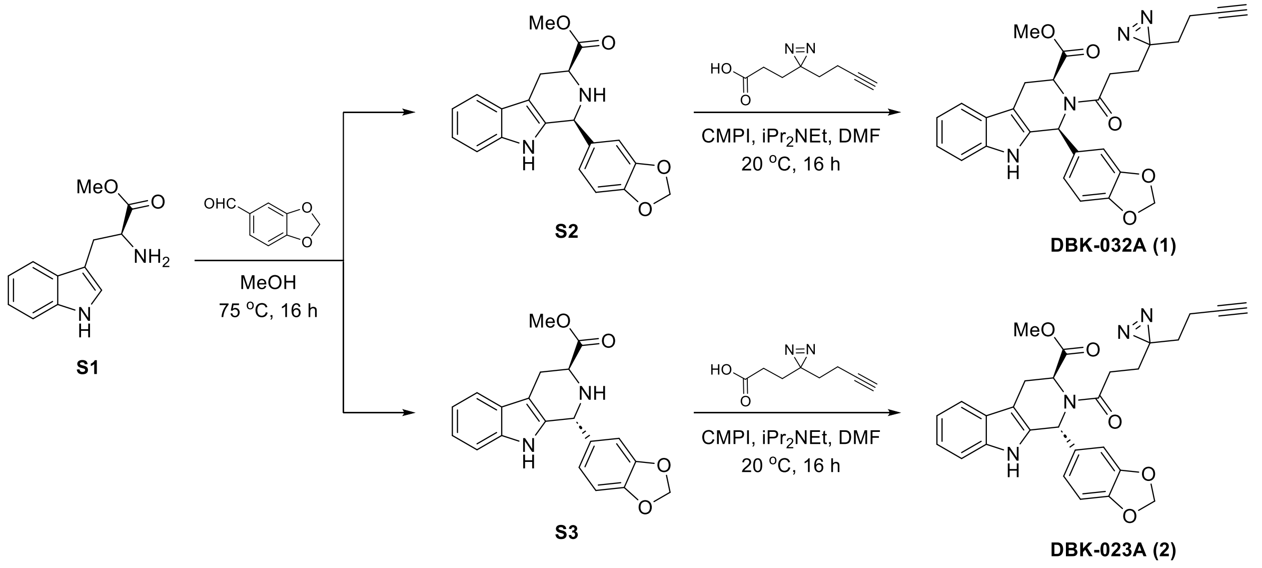


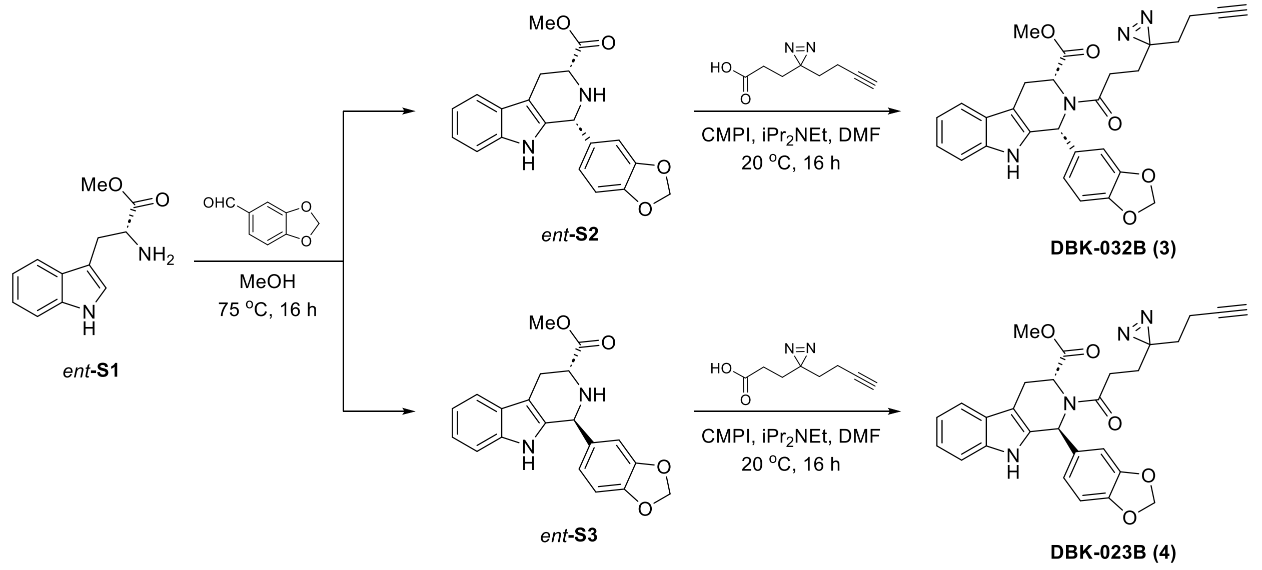

#### Synthesis of DBK-032A and DBK-032B

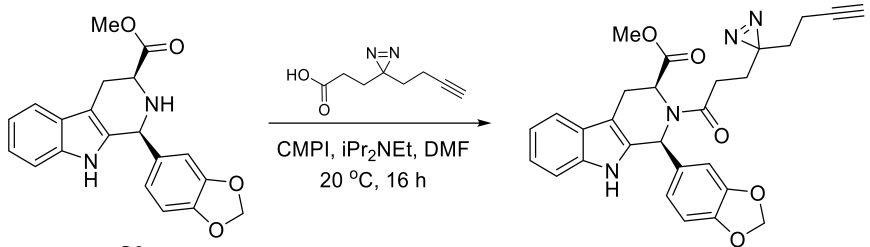


**S2**

**DBK-032A (1)**

##### methyl (1S,3S)-2-(3-(3-(but-3-yn-1-yl)-3H-diazirin-3-yl)propanoyl)-1-(benzo[d][1,3]dioxol-5-yl)-2,3,4,9-tetrahydro-1H-pyrido[3,4-b]indole-3-carboxylate (DBK-032A) (1)

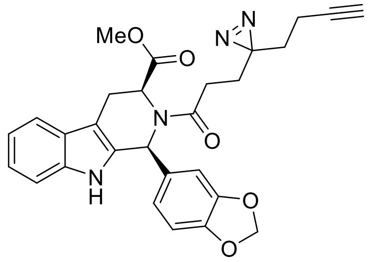


To a solution of **S2**[1] (80 mg, 0.23 mmol) in DMF (2 mL) were added 3-(3-(but-3-yn-1-yl)-3H-diazirin-3-yl)propanoic acid (57 mg, 0.34 mmol), $iPr_2NEt$ (89 mg, 0.69 mmol) and CMPI (120 mg, 0.46 mmol). The mixture was stirred at 30 °C for 12 hours. Upon completion, the resulting mixture was purified by prep-HPLC (column: Waters Xbridge 150 mm × 25 mm × 5 µm; mobile phase: [A: water (10 mM ammonium bicarbonate)–B: MeCN]; B%: 49% – 79%, 10 min) to obtain **DBK-032A** (60 mg, 53% yield) as an off-white solid.

**$^1$H-NMR** (400 MHz, $CD_3OD$): δ ppm 7.54 (d, J = 8.0 Hz, 1H), 7.29 (d, J = 8.0 Hz, 1H), 7.12 (t, J = 7.5 Hz, 1H), 7.06 (t, J = 7.4 Hz, 1H), 6.97 (s, 1H), 6.80 (s, 1H), 6.66 (d, J = 8.0 Hz, 1H), 6.54 (d, J = 8.0 Hz, 1H), 5.89 (s, 2H), 5.16 (d, J = 7.0 Hz, 1H), 3.60 (d, J = 15.9 Hz, 1H), 3.13 (s, 3H), 3.05 (dd, J = 6.6, 16.0 Hz, 1H), 2.48 (t, J = 7.2 Hz, 2H), 2.25 (s, 1H), 2.09 – 1.89 (m, 2H), 1.89 – 1.71 (t, J = 7.2 Hz, 2H), 1.66 (t, J = 7.2 Hz, 2H). 1 exchangeable proton not observed.

**HRMS ESI-TOF** m/z calculated for $C_{28}H_{27}N_4O_5$ $[M+H]^+$ 499.1976. Found 499.1978.

**methyl (1R,3R)-2-(3-(3-(but-3-yn-1-yl)-3H-diazirin-3-yl)propanoyl)-1-(benzo[d][1,3]dioxol-5-yl)-2,3,4,9-tetrahydro-1H-pyrido[3,4-b]indole-3-carboxylate (DBK-032B) (3)**

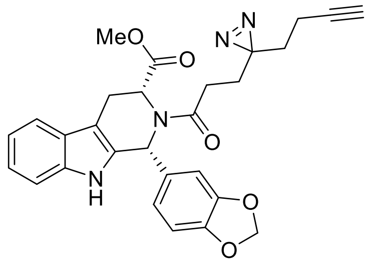


Prepared in an analogous fashion from *ent*-**S2**[1]. 32% yield as an off-white solid.

**$^1$H-NMR** (400 MHz, $CD_3OD$): δ ppm 7.54 (d, J = 7.8 Hz, 1H), 7.29 (d, J = 8.0 Hz, 1H), 7.20 – 7.01 (m, 2H), 6.97 (s, 1H), 6.80 (s, 1H), 6.67 (d, J = 8.1 Hz, 1H), 6.55 (d, J = 8.1 Hz, 1H), 5.90 (s, 2H), 5.16 (d, J = 6.9 Hz, 1H), 3.60 (d, J = 15.9 Hz, 1H), 3.14 (s, 3H), 3.11 – 3.01 (m, 1H), 2.49 (t, J = 7.3 Hz, 2H), 2.26 (t, J = 2.7 Hz, 1H), 2.05 (td, J = 7.5, 2.7 Hz, 2H), 1.87 (t, J = 7.3 Hz, 2H), 1.67 (t, J = 7.1 Hz, 2H). 1 exchangeable proton not observed.

**HRMS ESI-TOF** m/z calculated for $C_{28}H_{27}N_4O_5$ $[M+H]^+$ 499.1976. Found 499.1977.

**Synthesis of DBK-023A and DBK-023B**

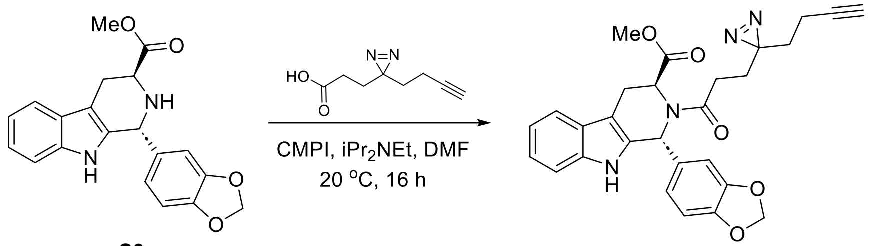


**S3** **DBK-023A (2)**

**methyl (1R,3S)-2-(3-(3-(but-3-yn-1-yl)-3H-diazirin-3-yl)propanoyl)-1-(benzo[d][1,3]dioxol-5-yl)-2,3,4,9-tetrahydro-1H-pyrido[3,4-b]indole-3-carboxylate (DBK-023A) (2)**

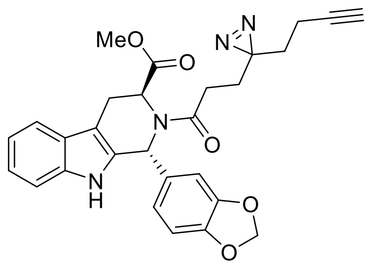


To a solution of **S3**[1] (80 mg, 0.23 mmol) in DMF (2 mL) were added 3-(3-(but-3-yn-1-yl)-3H-diazirin-3-yl)propanoic acid (57 mg, 0.34 mmol), $iPr_2NEt$ (89 mg, 0.69 mmol) and CMPI (120 mg, 0.46 mmol). The mixture was stirred at 30 °C for 12 hours. Upon completion, the resulting mixture was purified by prep-HPLC (column: Waters Xbridge 150 mm × 25 mm × 5 µm; mobile phase: [A: water (10 mM ammonium bicarbonate)–B: MeCN]; B%: 42% – 72%, 10 min) to obtain **DBK-023A** (40 mg, 35% yield) as an off-white solid.

**$^1$H-NMR** (400 MHz, $CD_3OD$): δ ppm 7.44 (d, J = 7.8 Hz, 1H), 7.31 – 7.19 (m, 1H), 7.12 – 6.64 (m, 5H), 6.20 – 5.79 (m, 3H), 5.40 – 5.07 (m, 1H), 3.60 (s, 3H), 3.46 (dd, J = 15.7, 5.1 Hz, 1H),

3.22 (dd, J = 15.9, 5.3 Hz, 1H), 2.37 (dt, J = 15.3, 7.3 Hz, 1H), 2.29 – 2.12 (m, 2H), 1.94 (dt, J = 7.7, 3.9 Hz, 2H), 1.77 – 1.43 (m, 4H). 1 exchangeable proton not observed.

**HRMS ESI-TOF** m/z calculated for $C_{28}H_{27}N_4O_5$ $[M+H]^+$ 499.1976. Found 499.1981.

**methyl (1S,3R)-2-(3-(3-(but-3-yn-1-yl)-3H-diazirin-3-yl)propanoyl)-1-(benzo[d][1,3]dioxol-5-yl)-2,3,4,9-tetrahydro-1H-pyrido[3,4-b]indole-3-carboxylate (DBK-023B) (4)**

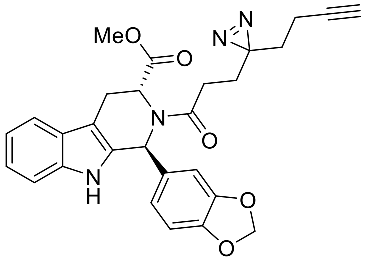


Prepared in an analogous fashion from *ent*-**S3**[1]. 31% yield as an off-white solid.

**$^1$H-NMR** (400 MHz, $CD_3OD$): δ ppm 7.43 (d, J = 7.6 Hz, 1H), 7.33 – 7.18 (m, 1H), 7.13 – 6.64 (m, 5H), 6.20 – 5.78 (m, 3H), 5.44 – 5.06 (m, 1H), 3.61 (s, 3H), 3.52 – 3.37 (m, 1H), 3.28 – 3.17 (m, 1H), 2.50 – 2.30 (m, 1H), 2.28 – 2.10 (m, 2H), 2.01 – 1.86 (m, 2H), 1.81 – 1.44 (m, 4H). 1 exchangeable proton not observed.

**HRMS ESI-TOF** m/z calculated for $C_{28}H_{27}N_4O_5$ $[M+H]^+$ 499.1976. Found 499.1980.

### Synthesis of azetidine photo-stereoprobes

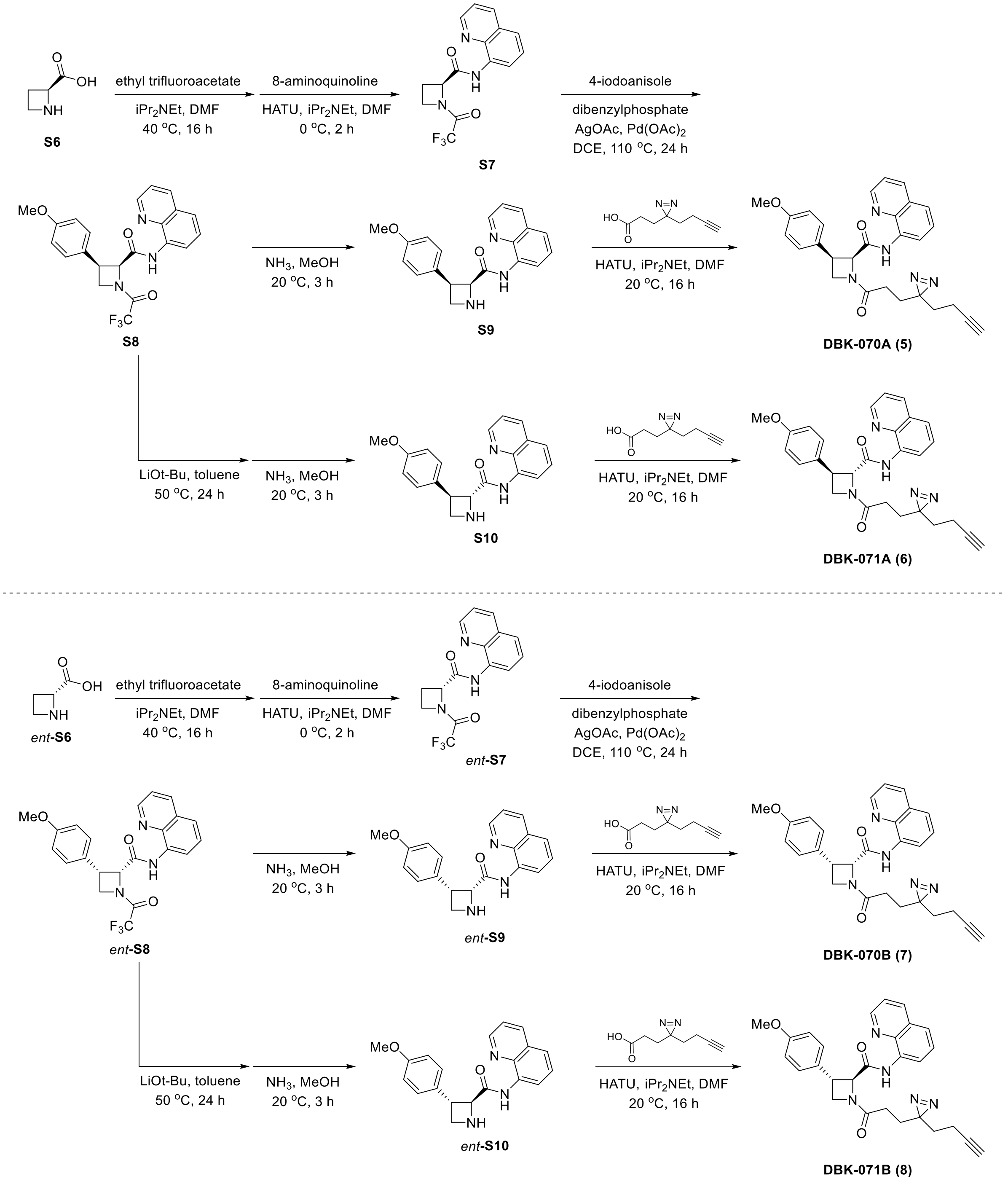

#### Synthesis of DBK-070A and DBK-070B

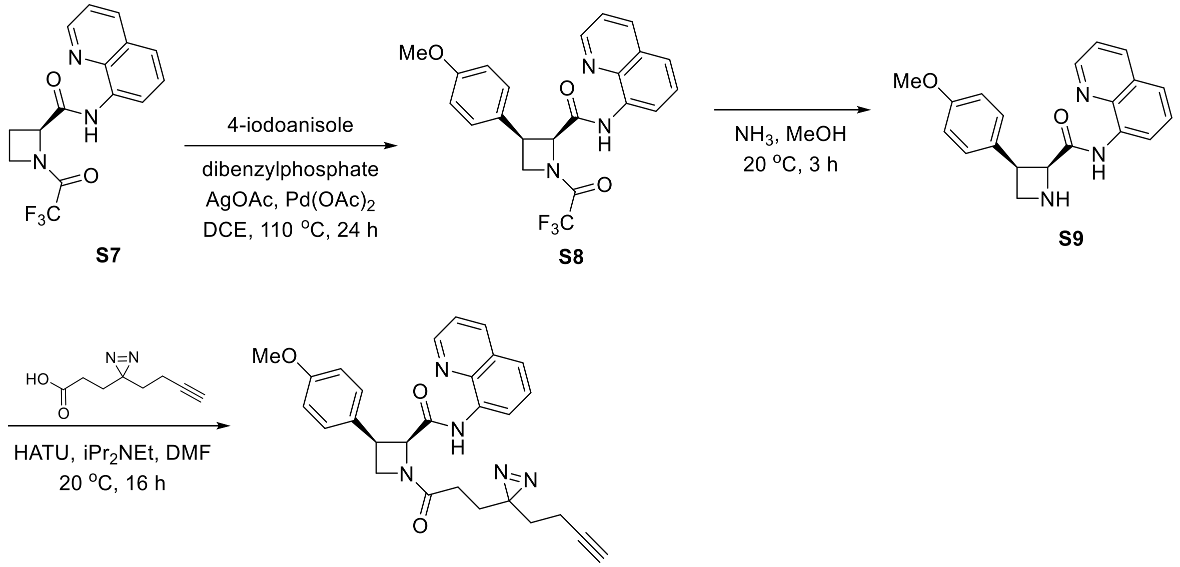


**DBK-070A (5)**

#### (2S,3R)-3-(4-methoxyphenyl)-N-(quinolin-8-yl)azetidine-2-carboxamide (S9)

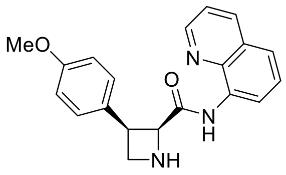

The reaction vessel was charged with **S7**[2] (180 mg, 0.56 mmol), dibenzylphosphate (62 mg, 0.22 mmol), AgOAc (190 mg, 1.1 mmol), 4-iodoanisole (390 mg, 1.7 mmol), and $Pd(OAc)_2$ (25 mg, 0.11 mmol). The reaction vessel was evacuated and refilled with $N_2$ (3×) prior to addition of DCE (1.0 mL). The reaction mixture was stirred at 110 °C for 24 hours. The reaction mixture was allowed to cool to ambient temperature, diluted with DCM, filtered through a pad of Celite, and eluted with DCM before the addition of 7 *N* $NH_3$ in MeOH (6.1 mL). The reaction mixture was stirred at 20 °C for 3 hours before being concentrated under reduced pressure. The residue was purified via prep-TLC (EtOAc only) to obtain **S9** (54 mg, 29% yield) as an off white solid.

**$^1$H-NMR** (400 MHz, $CD_3OD$): δ ppm 8.93 (dd, J = 4.2, 1.7 Hz, 1H), 8.33 (dd, J = 7.7, 1.3 Hz, 1H), 8.28 (dd, J = 8.4, 1.7 Hz, 1H), 7.61 – 7.52 (m, 2H), 7.43 (t, J = 8.0 Hz, 1H), 7.40 – 7.33 (m, 2H), 6.65 – 6.57 (m, 2H), 4.30 – 4.07 (m, 2H), 3.66 (dd, J = 7.5, 4.2 Hz, 1H), 3.55 (s, 3H).

**LC-MS** m/z calculated for $C_{20}H_{20}N_3O_2$ $[M+H]^+$ 334.2. Found 334.1.

**(2S,3R)-1-(3-(3-(but-3-yn-1-yl)-3H-diazirin-3-yl)propanoyl)-3-(4-methoxyphenyl)-N-(quinolin-8-yl)azetidine-2-carboxamide (DBK-070A) (5)**

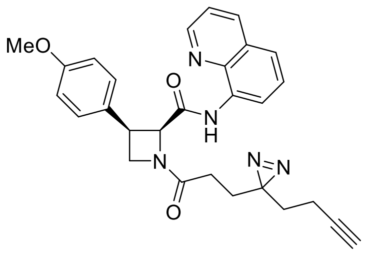

To a solution of **S9** (33 mg, 0.10 mmol) in DMF (0.1 mL) were added 3-(3-(but-3-yn-1-yl)-3H-diazirin-3-yl)propanoic acid (18 mg, 0.11 mmol), $iPr_2NEt$ (26 mg, 35 µL, 0.20 mmol) and HATU (46 mg, 0.12 mmol). The mixture was stirred at 20 °C for 12 hours. Upon completion, the resulting mixture was purified via flash column chromatography (Hexane/EtOAc = 2/1 to 5/4) to obtain **DBK-070A** (9.2 mg, 19% yield) as an off-white amorphous.

**$^1$H-NMR** (600 MHz, $CDCl_3$: mixture of rotamers): δ ppm 10.26 (s, 1H), 8.84 – 8.75 (m, 1H), 8.46 – 8.33 (m, 1H), 8.17 – 8.10 (m, 1H), 7.53 – 7.37 (m, 3H), 7.30 – 7.22 (m, 2H), 6.70 – 6.60 (m, 2H), 5.30 – 5.22 (m, 1H), 4.57 – 4.18 (m, 3H), 3.67 – 3.55 (m, 3H), 2.26 – 1.51 (m, 9H).

**HRMS ESI-TOF** m/z calculated for $C_{28}H_{28}N_5O_3$ $[M+H]^+$ 482.2187. Found 482.2191.

**(2R,3S)-3-(4-methoxyphenyl)-N-(quinolin-8-yl)azetidine-2-carboxamide (*ent*-S9)**

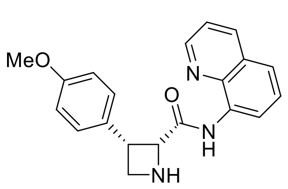

Prepared in an analogous fashion from *ent*-**S7**[2]. 29% yield as an off-white solid.

**$^1$H-NMR** (400 MHz, $CD_3OD$): δ ppm 8.93 (dd, J = 4.2, 1.7 Hz, 1H), 8.33 (dd, J = 7.7, 1.3 Hz, 1H), 8.28 (dd, J = 8.3, 1.7 Hz, 1H), 7.60 – 7.54 (m, 2H), 7.43 (t, J = 8.0 Hz, 1H), 7.41 – 7.33 (m, 2H), 6.65 – 6.56 (m, 2H), 4.30 – 4.07 (m, 2H), 3.66 (dd, J = 7.5, 4.2 Hz, 1H), 3.56 (s, 3H).

**LC-MS** m/z calculated for $C_{20}H_{20}N_3O_2$ $[M+H]^+$ 334.2. Found 334.1.

**(2R,3S)-1-(3-(3-(but-3-yn-1-yl)-3H-diazirin-3-yl)propanoyl)-3-(4-methoxyphenyl)-N-(quinolin-8-yl)azetidine-2-carboxamide (DBK-070B) (7)**

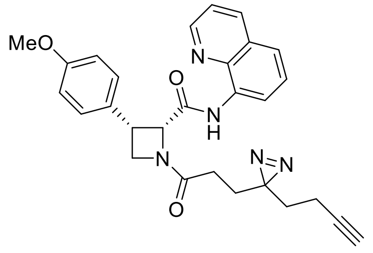


Prepared in an analogous fashion from *ent*-**S9**. 18% yield as an off-white solid.

**$^1$H-NMR** (600 MHz, $CDCl_3$: mixture of rotamers): δ ppm 10.26 (s, 1H), 8.84 – 8.75 (m, 1H), 8.46 – 8.32 (m, 1H), 8.18 – 8.09 (m, 1H), 7.53 – 7.36 (m, 3H), 7.30 – 7.22 (m, 2H), 6.69 – 6.59 (m, 2H), 5.30 – 5.21 (m, 1H), 4.58 – 4.18 (m, 3H), 3.67 – 3.55 (m, 3H), 2.28 – 1.51 (m, 9H).

**HRMS ESI-TOF** m/z calculated for $C_{28}H_{28}N_5O_3$ $[M+H]^+$ 482.2187. Found 482.2191.

**Synthesis of DBK-071A and DBK-071B**

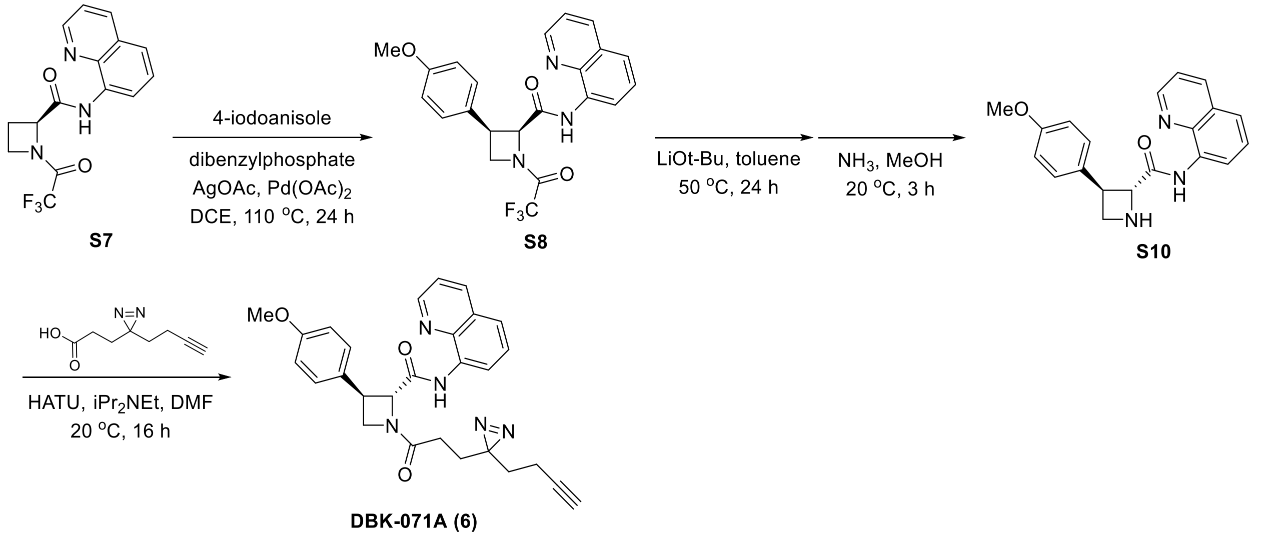


**(2S,3S)-3-(4-methoxyphenyl)-N-(quinolin-8-yl)azetidine-2-carboxamide (S10)**

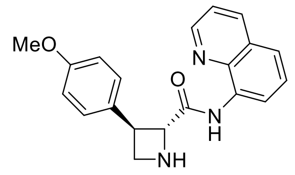


The reaction vessel was charged with **S7**[2] (200 mg, 0.62 mmol), dibenzylphosphate (69 mg, 0.25 mmol), AgOAc (210 mg, 1.2 mmol), 4-iodoanisole (430 mg, 1.9 mmol), and $Pd(OAc)_2$ (28 mg, 0.12 mmol). The reaction vessel was evacuated and refilled with $N_2$ (3×) prior to addition of DCE (0.62 mL). The reaction mixture was stirred at 110 °C for 24 hours. The reaction mixture was allowed to cool to ambient temperature, diluted with DCM, filtered through a pad of Celite. The solvent was removed under reduced pressure and Li*Ot*-Bu (120 mg, 1.6 mmol), was added to the reaction mixture. The reaction vessel was evacuated and refilled with $N_2$ (3×) prior to addition of toluene (3.1 mL), then warmed up to 50 °C and stirred for 24 hours. The reaction mixture was allowed to cool to ambient temperature, filtered through a pad of Celite, and eluted with DCM before the addition of 7 *N* $NH_3$ in MeOH (3.1 mL). The reaction mixture was stirred at 20 °C for 3 hours before being concentrated under reduced pressure. The residue was purified via flash column chromatography (Hexane/EtOAc = 4/1 to 7/3) to obtain **S10** (63 mg, 30% yield) as an off white solid.

**$^1$H-NMR** (400 MHz, $CD_3OD$): δ ppm 8.90 (dd, J = 4.2, 1.7 Hz, 1H), 8.73 (dd, J = 7.6, 1.4 Hz, 1H), 8.30 (dd, J = 8.3, 1.8 Hz, 1H), 7.64 (dd, J = 8.2, 1.4 Hz, 1H), 7.60 – 7.51 (m, 2H), 7.42 – 7.33 (m, 2H), 6.97 – 6.89 (m, 2H), 4.59 – 4.48 (m, 1H), 3.96 – 3.84 (m, 2H), 3.81 (s, 3H), 3.79 – 3.70 (m, 1H).

**LC-MS** m/z calculated for $C_{20}H_{20}N_3O_2$ $[M+H]^+$ 334.2. Found 334.1.

**(2S,3S)-1-(3-(3-(but-3-yn-1-yl)-3H-diazirin-3-yl)propanoyl)-3-(4-methoxyphenyl)-N-(quinolin-8-yl)azetidine-2-carboxamide (DBK-071A) (6)**

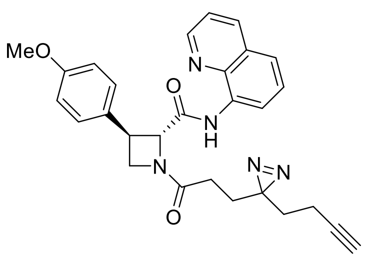


To a solution of **S10** (33 mg, 0.10 mmol) in DMF (0.1 mL) were added 3-(3-(but-3-yn-1-yl)-3H-diazirin-3-yl)propanoic acid (18 mg, 0.11 mmol), $iPr_2NEt$ (26 mg, 35 μL, 0.20 mmol) and HATU (46 mg, 0.12 mmol). The mixture was stirred at 20 °C for 12 hours. Upon completion, the resulting mixture was purified via flash column chromatography (Hexane/EtOAc = 9/1 to 9/2) to obtain **DBK-071A** (12 mg, 26% yield) as an off-white amorphous.

**$^1$H-NMR** (600 MHz, $CDCl_3$: mixture of rotamers): δ ppm 1H NMR (600 MHz, CDCl3) δ 10.79 (s, 0.7H), 10.54 (s, 0.3H), 8.85 – 8.77 (m, 2H), 8.23 – 8.08 (m, 1H), 7.64 – 7.27 (m, 5H), 6.93 (d, J = 8.2 Hz, 2H), 5.10 – 4.81 (m, 1H), 4.61 – 4.42 (m, 1H), 4.36 – 4.06 (m, 1.8H), 3.96 – 3.74 (m, 3.2H), 2.24 – 1.50 (m, 9H).

**HRMS ESI-TOF** m/z calculated for $C_{28}H_{28}N_5O_3$ $[M+H]^+$ 482.2187. Found 482.2192.

**(2R,3R)-3-(4-methoxyphenyl)-N-(quinolin-8-yl)azetidine-2-carboxamide (*ent*-S10)**

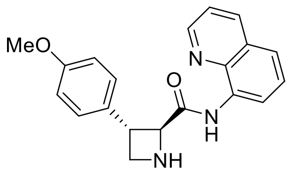

Prepared in an analogous fashion from *ent*-**S7**[2]. 38% yield as an off-white solid.

**$^1$H-NMR** (400 MHz, $CD_3OD$): δ ppm 8.92 (dd, J = 4.2, 1.7 Hz, 1H), 8.75 (dd, J = 7.6, 1.4 Hz, 1H), 8.33 (dd, J = 8.3, 1.7 Hz, 1H), 7.66 (dd, J = 8.3, 1.3 Hz, 1H), 7.62 – 7.54 (m, 2H), 7.44 – 7.31 (m, 2H), 7.01 – 6.89 (m, 2H), 4.62 – 4.51 (m, 1H), 3.96 – 3.87 (m, 2H), 3.83 (s, 3H), 3.81 – 3.76 (m, 1H).

**LC-MS** m/z calculated for $C_{20}H_{20}N_3O_2$ $[M+H]^+$ 334.2. Found 334.1.

**(2R,3R)-1-(3-(3-(but-3-yn-1-yl)-3H-diazirin-3-yl)propanoyl)-3-(4-methoxyphenyl)-N-(quinolin-8-yl)azetidine-2-carboxamide (DBK-071B) (8)**

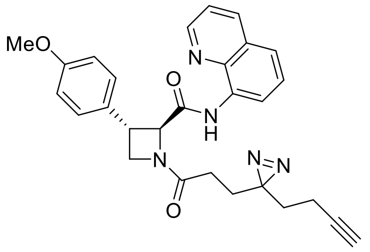

Prepared in an analogous fashion from *ent*-**S10**. 23% yield as an off-white amorphous.

**$^1$H-NMR** (600 MHz, $CDCl_3$: mixture of rotamers): δ ppm 1H NMR (600 MHz, CDCl3) δ 10.80 (s, 0.7H), 10.55 (s, 0.3H), 8.86 – 8.76 (m, 2H), 8.23 – 8.08 (m, 1H), 7.63 – 7.29 (m, 5H), 6.93 (d, J = 8.1 Hz, 2H), 5.06 – 4.81 (m, 1H), 4.61 – 4.41 (m, 1H), 4.34 – 4.06 (m, 1.8H), 3.96 – 3.75 (m, 3.2H), 2.20 – 1.46 (m, 9H).

**HRMS ESI-TOF** m/z calculated for $C_{28}H_{28}N_5O_3$ $[M+H]^+$ 482.2187. Found 482.2190.

#### Synthesis of pyrrolidine photo-stereoprobes

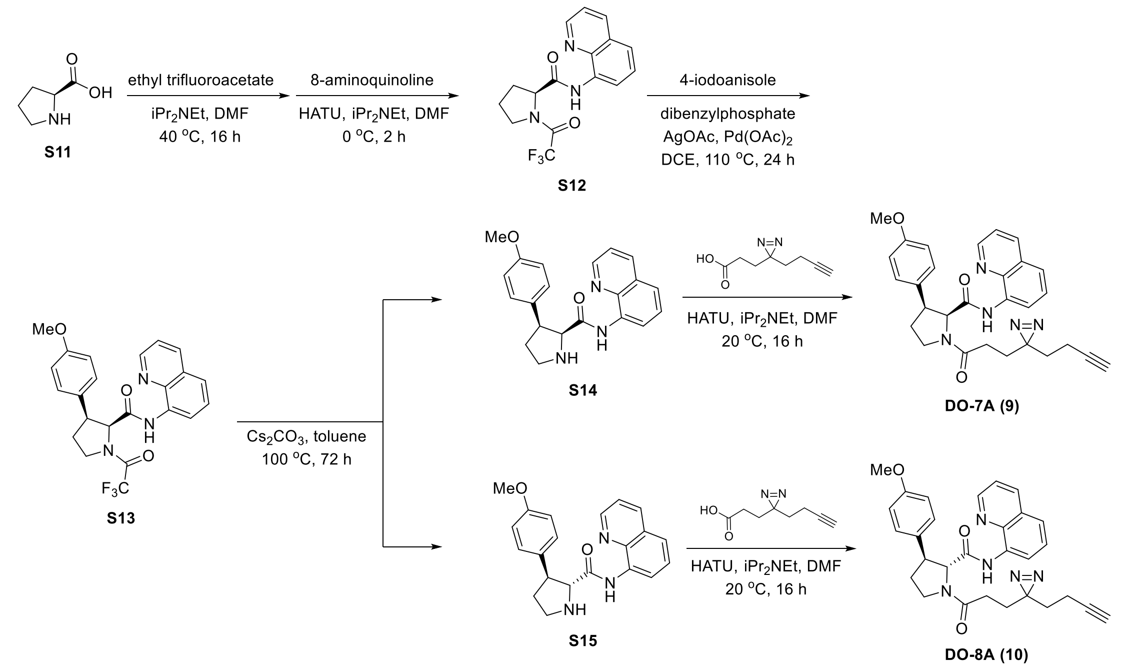


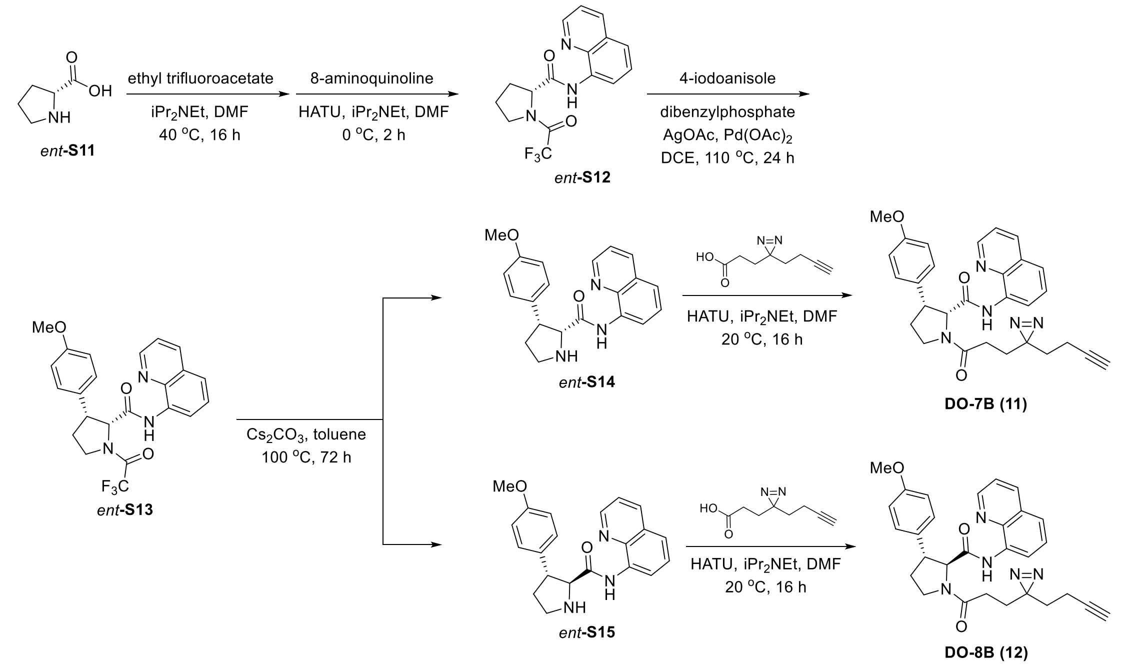

**Synthesis of S13 and *ent*-S13**

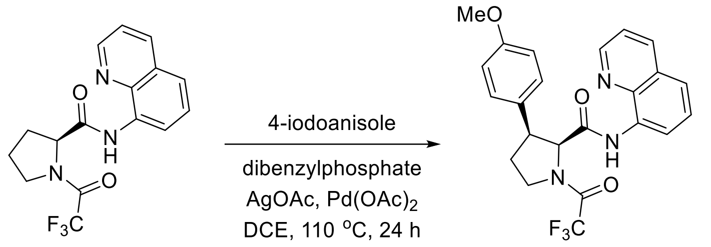


**(2S,3S)-1-(2,2,2-trifluoroacetyl)-3-(4-methoxyphenyl)-N-(quinolin-8-yl)pyrrolidine-2-carboxamide (S13)**

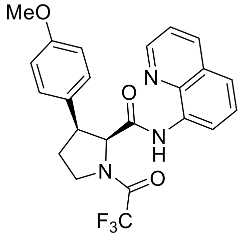


The reaction vessel was charged with **S12**[2] (380 mg, 1.1 mmol), dibenzylphosphate (130 mg, 0.45 mmol), AgOAc (380 mg, 2.3 mmol), 4-iodoanisole (790 mg, 3.4 mmol), and $Pd(OAc)_2$ (51 mg, 0.23 mmol). The reaction vessel was evacuated and refilled with $N_2$ (3×) prior to addition of DCE (1.0 mL). The reaction mixture was stirred at 110 °C for 24 hours. The reaction mixture was allowed to cool to ambient temperature, diluted with DCM, filtered through a pad of $SiO_2$, eluted with Hexane/EtOAc (1/1) and concentrated under reduced pressure. The residue was purified via Teledyne Isco Combiflash Rf (40g, [A: Hexane–B: EtOAc]; B%: 0 – 35%, 16 min) to obtain **S13** (400 mg, 79% yield) as a white solid.

**$^1$H-NMR** (600 MHz, $CDCl_3$: mixture of rotamers): δ ppm 9.39 (s, 0.8H), 9.12 (s, 0.2H), 8.62 (dd, J = 4.2, 1.6 Hz, 0.8H), 8.58 (dd, J = 4.2, 1.7 Hz, 0.2H), 8.52 – 8.49 (m, 0.2H), 8.49 – 8.45 (m, 0.8H), 8.08 (dd, J = 8.2, 1.7 Hz, 0.2H), 8.06 (dd, J = 8.3, 1.7 Hz, 0.8H), 7.47 – 7.29 (m, 3H), 7.21 – 7.15 (m, 2H), 6.58 – 6.51 (m, 2H), 4.93 (d, J = 8.3 Hz, 0.8H), 4.90 (d, J = 7.9 Hz, 0.2H), 4.32 – 4.26 (m, 0.8H), 4.22 – 4.17 (m, 0.2H), 3.91 – 3.79 (m, 1.2H), 3.75 – 3.69 (m, 0.8H), 3.42 (s, 2.4H), 3.38 (s, 0.6H), 2.94 – 2.82 (m, 0.8H), 2.72 – 2.63 (m, 0.2H), 2.30 (dt, J = 12.5, 6.4 Hz, 0.8H), 2.24 (dt, J = 11.8, 6.7 Hz, 0.2H).

**LC-MS** m/z calculated for $C_{23}H_{21}F_3N_3O_3$ $[M+H]^+$ 444.2. Found 444.2.

**(2R,3R)-1-(2,2,2-trifluoroacetyl)-3-(4-methoxyphenyl)-N-(quinolin-8-yl)pyrrolidine-2-carboxamide (*ent*-S13)**

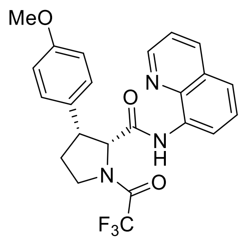

Prepared in an analogous fashion from *ent*-**S12**[2]. 66% yield as a white solid.

**$^1$H-NMR** (600 MHz, $CDCl_3$: mixture of rotamers): δ ppm 9.40 (s, 0.8H), 9.13 (s, 0.2H), 8.62 (dd, J = 4.2, 1.7 Hz, 0.8H), 8.59 (dd, J = 4.2, 1.7 Hz, 0.2H), 8.52 – 8.49 (m, 0.2H), 8.49 – 8.45 (m, 0.8H), 8.09 (dd, J = 8.2, 1.7 Hz, 0.2H), 8.06 (dd, J = 8.2, 1.7 Hz, 0.8H), 7.47 – 7.29 (m, 3H), 7.21 – 7.15 (m, 2H), 6.58 – 6.51 (m, 2H), 4.93 (d, J = 8.3 Hz, 0.8H), 4.90 (d, J = 8.0 Hz, 0.2H), 4.32 – 4.26 (m, 0.8H), 4.22 – 4.17 (m, 0.2H), 3.91 – 3.80 (m, 1.2H), 3.75 – 3.69 (m, 0.8H), 3.42 (s, 2.4H), 3.39 (s, 0.6H), 2.94 – 2.84 (m, 0.8H), 2.72 – 2.62 (m, 0.2H), 2.30 (dt, J = 12.5, 6.4 Hz, 0.8H), 2.25 (dt, J = 13.1, 6.5 Hz, 0.2H).

**LC-MS** m/z calculated for $C_{23}H_{21}F_3N_3O_3$ $[M+H]^+$ 444.2. Found 444.2.

**Synthesis of DO-7A and DO-8A**

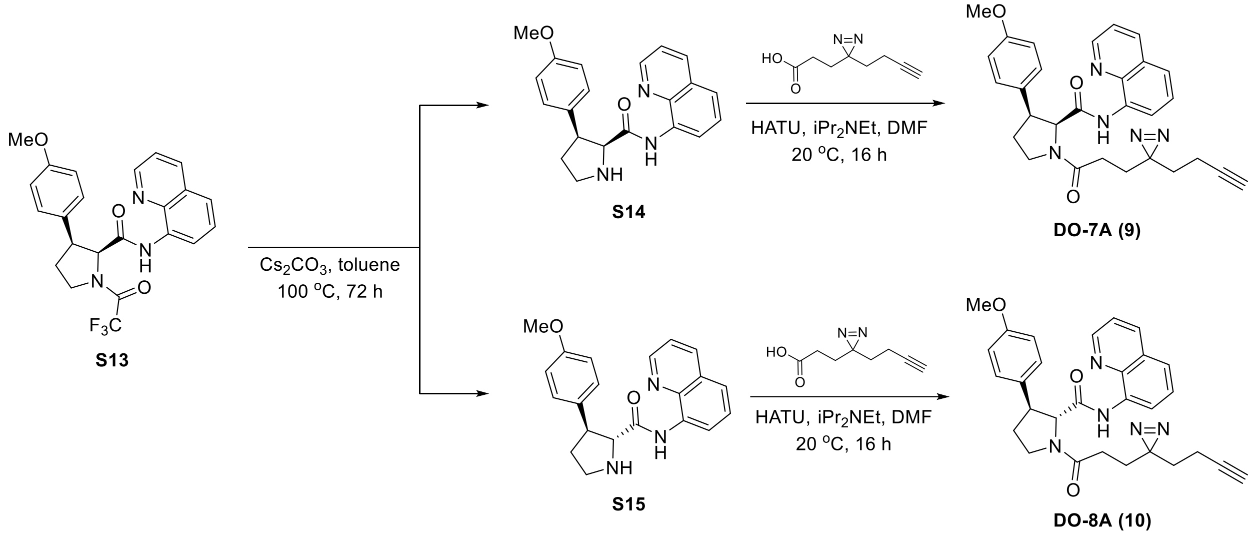


**(2S,3S)-3-(4-methoxyphenyl)-N-(quinolin-8-yl)pyrrolidine-2-carboxamide (S14)**

**(2R,3S)-3-(4-methoxyphenyl)-N-(quinolin-8-yl)pyrrolidine-2-carboxamide (S15)**

To a solution of **S13** (100 mg, 0.23 mmol) in dry toluene (3.5 mL) were added $Cs_2CO_3$ (400 mg, 1.2 mmol). The mixture was stirred at 100 °C for 72 hours. The resulting mixture was then diluted with EtOAc and washed with sat.$NaHCO_3$ (x1). The organic layer was dried over $Na_2SO_4$, filtered, and concentrated in vacuo. The residue was purified via prep-TLC (EtOAc 100%) to obtain **S14** (27 mg, 34% yield) and **S15** (23 mg, 30% yield) as off-white solids.

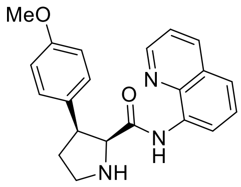

$^1$**H-NMR** (600 MHz, $CDCl_3$): δ ppm 10.87 (s, 1H), 8.83 (dd, *J* = 4.2, 1.7 Hz, 1H), 8.43 (dd, *J* = 7.2, 1.8 Hz, 1H), 8.10 (dd, *J* = 8.2, 1.7 Hz, 1H), 7.47 – 7.37 (m, 3H), 7.21 – 7.14 (m, 2H), 6.62 – 6.56 (m, 2H), 4.20 (d, *J* = 8.9 Hz, 1H), 3.72 (q, *J* = 8.0 Hz, 1H), 3.59 – 3.53 (m, 4H), 3.26 – 3.20 (m, 1H), 2.60 (brs, 1H), 2.31 – 2.22 (m, 1H), 2.19 – 2.09 (m, 1H).

**LC-MS** m/z calculated for $C_{21}H_{22}N_3O_2$ $[M+H]^+$ 348.2. Found 348.2.

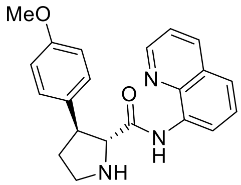

$^1$**H-NMR** (600 MHz, $CDCl_3$): δ ppm 11.28 (s, 1H), 8.85 – 8.78 (m, 2H), 8.13 (dd, J = 8.2, 1.7 Hz, 1H), 7.56 – 7.47 (m, 2H), 7.43 (dd, J = 8.2, 4.2 Hz, 1H), 7.37 – 7.29 (m, 2H), 6.92 – 6.87 (m, 2H), 4.00 (d, J = 6.2 Hz, 1H), 3.81 (s, 3H), 3.58 (td, J = 7.6, 6.1 Hz, 1H), 3.42 – 3.33 (m, 1H), 3.23 (ddd, J = 10.9, 8.0, 6.7 Hz, 1H), 2.91 (brs, 1H), 2.34 – 2.25 (m, 1H), 2.07 – 1.97 (m, 1H).

**LC-MS** m/z calculated for $C_{21}H_{22}N_3O_2$ $[M+H]^+$ 348.2. Found 348.2.

**(2S,3S)-1-(3-(3-(but-3-yn-1-yl)-3H-diazirin-3-yl)propanoyl)-3-(4-methoxyphenyl)-N-(quinolin-8-yl)pyrrolidine-2-carboxamide (DO-7A) (9)**

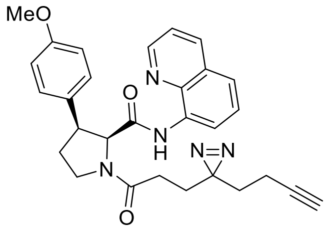

To a solution of **S14** (36 mg, 0.10 mmol) in dry DMF (0.4 mL) were added 3-(3-(but-3-yn-1-yl)-3H-diazirin-3-yl)propanoic acid (18 mg, 0.11 mmol), $iPr_2NEt$ (36 μL, 0.21 mmol) and HATU (47 mg, 0.12 mmol). The mixture was stirred at 25 °C for 16 hours. Upon completion, the resulting mixture was purified by prep-TLC (Hexane/EtOAc = 1/1) to obtain **DO-7A** (25 mg, 49% yield) as a colorless amorphous.

$^1$**H-NMR** (600 MHz, $CDCl_3$: mixture of rotamers): δ ppm 9.52 (s, 0.2H), 9.45 (s, 0.8H), 8.69 (dd, J = 4.2, 1.7 Hz, 0.2H), 8.64 (dd, J = 4.2, 1.7 Hz, 0.8H), 8.50 – 8.45 (m, 1H), 8.12 (dd, J = 8.3, 1.7 Hz, 0.2H), 8.05 (dd, J = 8.2, 1.7 Hz, 0.8H), 7.54 – 7.30 (m, 3H), 7.22 – 7.11 (m, 2H), 6.69 – 6.47 (m, 2H), 4.89 (d, J = 8.2 Hz, 0.8H), 4.61 (d, J = 8.5 Hz, 0.2H), 4.19 – 4.14 (m, 0.2H), 4.02 – 3.91 (m, 0.8H), 3.85 – 3.78 (m, 0.2H), 3.73 – 3.60 (m, 1.8H), 3.49 (s, 0.6H), 3.44 (s, 2.4H), 2.90 – 2.80 (m, 0.8H), 2.67 – 2.55 (m, 0.2H), 2.39 – 1.39 (m, 10H).

**HRMS ESI-TOF** m/z calculated for $C_{29}H_{30}N_5O_3$ $[M+H]^+$ 496.2343. Found 496.2345.

**(2R,3S)-1-(3-(3-(but-3-yn-1-yl)-3H-diazirin-3-yl)propanoyl)-3-(4-methoxyphenyl)-N-(quinolin-8-yl)pyrrolidine-2-carboxamide (DO-8A) (10)**

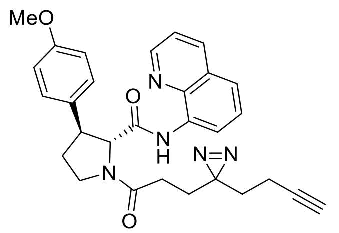

To a solution of **S15** (18 mg, 0.051 mmol) in dry DMF (0.3 mL) were added 3-(3-(but-3-yn-1-yl)-3H-diazirin-3-yl)propanoic acid (8.9 mg, 0.054 mmol), $iPr_2NEt$ (18 µL, 0.10 mmol) and HATU (23 mg, 0.061 mmol). The mixture was stirred at 25 °C for 16 hours. Upon completion, the resulting mixture was purified by prep-TLC (Hexane/EtOAc = 1/1) to obtain **DO-8A** (18 mg, 70% yield) as a colorless amorphous.

**$^1$H-NMR** (600 MHz, $CDCl_3$: mixture of rotamers): δ ppm 10.24 (s, 0.8H), 10.17 (s, 0.2H), 8.76 – 8.70 (m, 2H), 8.17 (dd, J = 8.2, 1.7 Hz, 0.2H), 8.11 (dd, J = 8.3, 1.7 Hz, 0.8H), 7.58 – 7.47 (m, 2H), 7.45 (dd, J = 8.3, 4.2 Hz, 0.2H), 7.41 (dd, J = 8.2, 4.2 Hz, 0.8H), 7.22 – 7.15 (m, 2H), 6.92 – 6.83 (m, 2H), 4.83 (d, J = 4.9 Hz, 0.8H), 4.50 (d, J = 4.5 Hz, 0.2H), 4.15 – 4.09 (m, 0.2H), 3.87 – 3.68 (m, 5.8H), 2.59 – 2.51 (m, 0.8H), 2.43 – 2.35 (m, 0.2H), 2.32 – 1.45 (m, 10H).

**HRMS ESI-TOF** m/z calculated for $C_{29}H_{30}N_5O_3$ $[M+H]^+$ 496.2343. Found 496.2346.

**(2R,3R)-3-(4-methoxyphenyl)-N-(quinolin-8-yl)pyrrolidine-2-carboxamide (*ent*-S14)**

**(2S,3R)-3-(4-methoxyphenyl)-N-(quinolin-8-yl)pyrrolidine-2-carboxamide (*ent*-S15)**

Prepared in an analogous fashion from *ent*-**S13**. 44% yield (*ent*-**S14**) and 27% yield (*ent*-**S15**) as off-white solids.

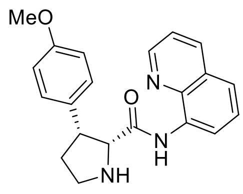

**$^1$H-NMR** (600 MHz, $CDCl_3$): δ ppm 10.89 (s, 1H), 8.83 (dd, *J* = 4.2, 1.7 Hz, 1H), 8.43 (dd, *J* = 7.2, 1.8 Hz, 1H), 8.09 (dd, *J* = 8.3, 1.7 Hz, 1H), 7.46 – 7.37 (m, 3H), 7.20 – 7.14 (m, 2H), 6.62 – 6.56 (m, 2H), 4.20 (d, *J* = 8.9 Hz, 1H), 3.72 (q, *J* = 8.0 Hz, 1H), 3.59 – 3.53 (m, 4H), 3.26 – 3.20 (m, 1H), 2.56 (brs, 1H), 2.30 – 2.22 (m, 1H), 2.19 – 2.10 (m, 1H).

**LC-MS** m/z calculated for $C_{21}H_{22}N_3O_2$ $[M+H]^+$ 348.2. Found 348.2.

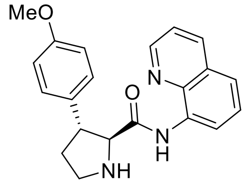

**$^1$H-NMR** (600 MHz, $CDCl_3$): δ ppm 11.30 (s, 1H), 8.85 – 8.78 (m, 2H), 8.13 (dd, J = 8.2, 1.7 Hz, 1H), 7.57 – 7.47 (m, 2H), 7.43 (dd, J = 8.2, 4.2 Hz, 1H), 7.36 – 7.29 (m, 2H), 6.92 – 6.86 (m, 2H), 4.00 (d, J = 6.2 Hz, 1H), 3.81 (s, 3H), 3.59 (td, J = 7.6, 6.1 Hz, 1H), 3.42 – 3.33 (m, 1H), 3.23 (ddd, J = 10.9, 8.0, 6.7 Hz, 1H), 2.80 (brs, 1H), 2.34 – 2.25 (m, 1H), 2.07 – 1.97 (m, 1H).

**LC-MS** m/z calculated for $C_{21}H_{22}N_3O_2$ $[M+H]^+$ 348.2. Found 348.2.

**(2R,3R)-1-(3-(3-(but-3-yn-1-yl)-3H-diazirin-3-yl)propanoyl)-3-(4-methoxyphenyl)-N-(quinolin-8-yl)pyrrolidine-2-carboxamide (DO-7B) (11)**

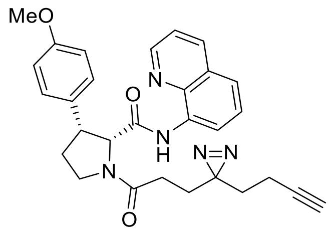

Prepared in analogous fashion from *ent*-**S14**. 42% yield as a colorless amorphous.

**$^1$H-NMR** (600 MHz, $CDCl_3$: mixture of rotamers): δ ppm 9.52 (s, 0.2H), 9.45 (s, 0.8H), 8.69 (dd, J = 4.2, 1.7 Hz, 0.2H), 8.64 (dd, J = 4.2, 1.7 Hz, 0.8H), 8.51 – 8.44 (m, 1H), 8.12 (dd, J = 8.3, 1.7 Hz, 0.2H), 8.05 (dd, J = 8.3, 1.7 Hz, 0.8H), 7.52 – 7.32 (m, 3H), 7.21 – 7.13 (m, 2H), 6.62 – 6.54 (m, 2H), 4.89 (d, J = 8.3 Hz, 0.8H), 4.61 (d, J = 8.5 Hz, 0.2H), 4.20 – 4.13 (m, 0.2H), 4.00 – 3.92 (m, 0.8H), 3.84 – 3.77 (m, 0.2H), 3.73 – 3.62 (m, 1.8H), 3.49 (s, 0.6H), 3.44 (s, 2.4H), 2.90 – 2.79 (m, 0.8H), 2.66 – 2.55 (m, 0.2H), 2.37 – 1.49 (m, 10H).

**HRMS ESI-TOF** m/z calculated for $C_{29}H_{30}N_5O_3$ $[M+H]^+$ 496.2343. Found 496.2344.

**(2S,3R)-1-(3-(3-(but-3-yn-1-yl)-3H-diazirin-3-yl)propanoyl)-3-(4-methoxyphenyl)-N-(quinolin-8-yl)pyrrolidine-2-carboxamide (DO-8B) (12)**

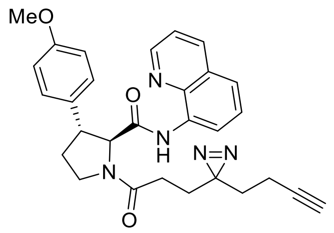

Prepared in analogous fashion from *ent*-**S15**. 73% yield as a colorless amorphous.

**$^1$H-NMR** (600 MHz, $CDCl_3$: mixture of rotamers): δ ppm 10.24 (s, 0.8H), 10.17 (s, 0.2H), 8.76 – 8.71 (m, 2H), 8.17 (dd, J = 8.2, 1.7 Hz, 0.2H), 8.11 (dd, J = 8.3, 1.7 Hz, 0.8H), 7.58 – 7.47 (m, 2H), 7.45 (dd, J = 8.3, 4.2 Hz, 0.2H), 7.41 (dd, J = 8.2, 4.2 Hz, 0.8H), 7.22 – 7.15 (m, 2H), 6.93 – 6.84 (m, 2H), 4.83 (d, J = 4.8 Hz, 0.8H), 4.50 (d, J = 4.5 Hz, 0.2H), 4.15 – 4.08 (m, 0.2H), 3.87 – 3.68 (m, 5.8H), 2.59 – 2.51 (m, 0.8H), 2.43 – 2.35 (m, 0.2H), 2.31 – 1.49 (m, 10H).

**HRMS ESI-TOF** m/z calculated for $C_{29}H_{30}N_5O_3$ $[M+H]^+$ 496.2343. Found 496.2345.

**Synthesis of WX-02-18, WX-02-22, WX-02-20, WX-02-21, WX-02-19, and WX-02-221**

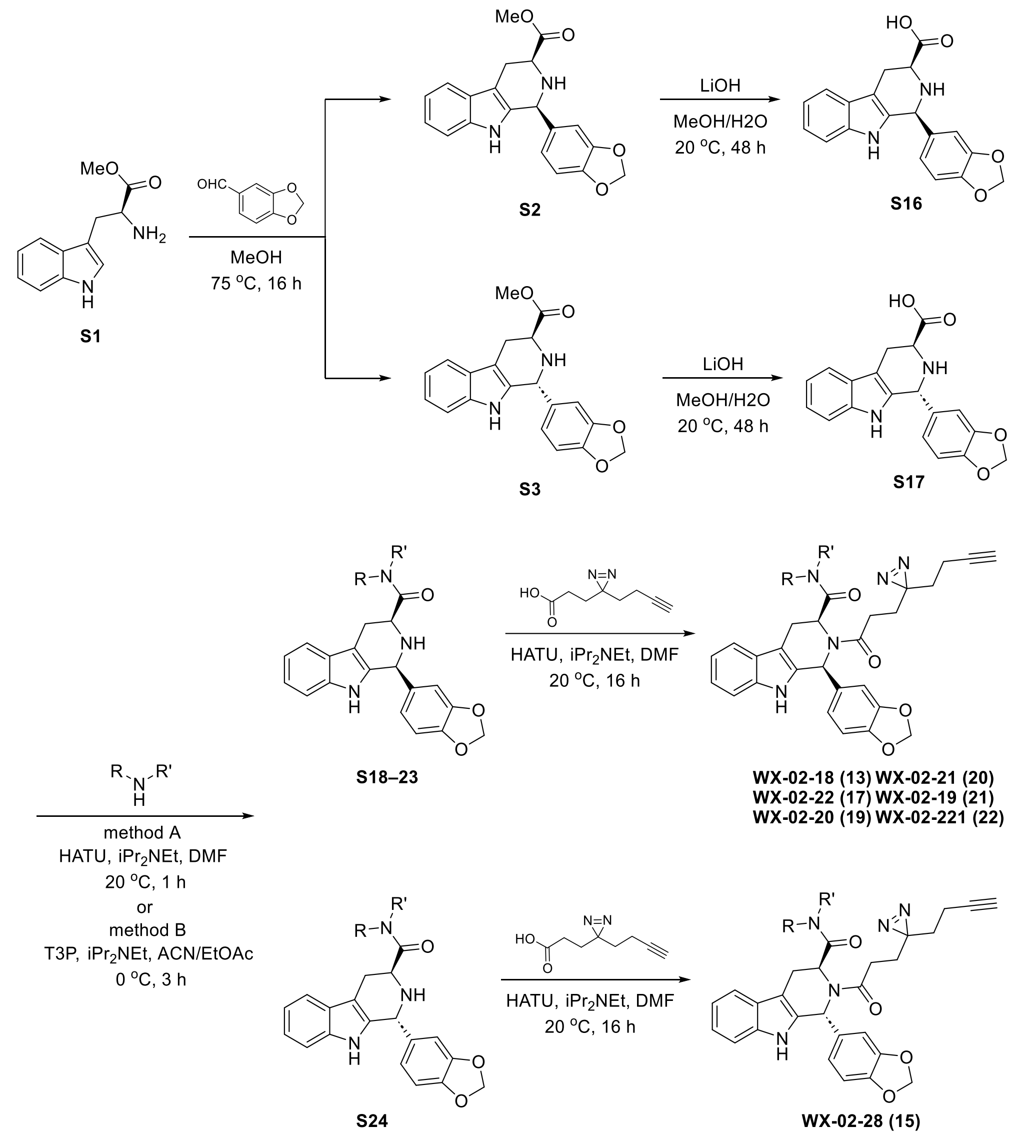

#### Synthesis of WX-02-38, WX-02-42, and WX-02-48

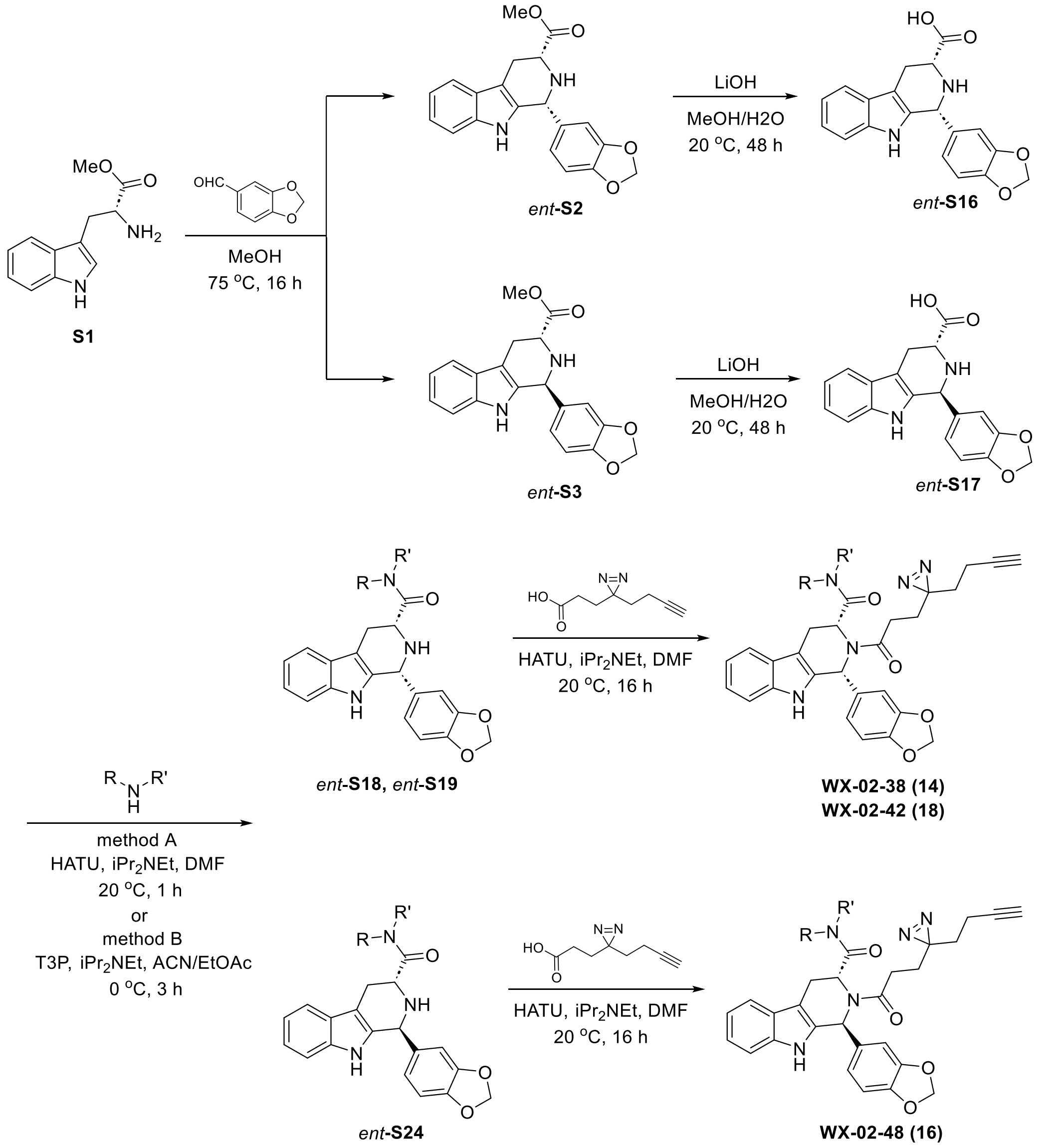

#### Synthesis of WX-02-18 and WX-02-38

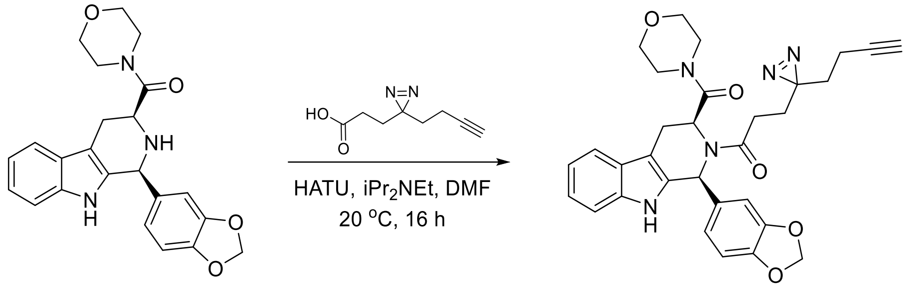


**S18** **WX-02-18 (13)**

#### 1-((1S,3S)-1-(benzo[d][1,3]dioxol-5-yl)-3-(morpholine-4-carbonyl)-1,3,4,9-tetrahydro-2H-pyrido[3,4-b]indol-2-yl)-3-(3-(but-3-yn-1-yl)-3H-diazirin-3-yl)propan-1-one (WX-02-18) (13)

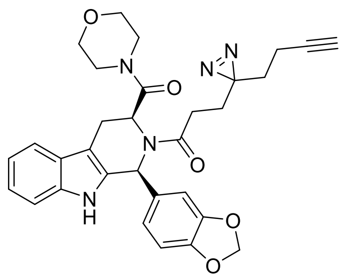

To a solution of **S18**[3] (50 mg, 0.12 mmol) in DMF (1 mL) were added 3-(3-(but-3-yn-1-yl)-3H-diazirin-3-yl)propanoic acid (21 mg, 0.12 mmol), iPr$_2$NEt (24 mg, 0.19 mmol) and HATU (56 mg, 0.15 mmol). The mixture was stirred at 20 °C for 16 hours. Upon completion, the reaction mixture was diluted with water (30 mL) and extracted with EtOAc (40 mL×3). The organic layer was dried over $Na_2SO_4$, filtered, and concentrated in vacuo. The residue was purified by prep-HPLC (column: Waters Xbridge 150 mm × 25 mm × 5 µm; mobile phase: [A: water (10 mM ammonium bicarbonate)–B: MeCN]; B%: 35% – 68%, 9 min) to obtain **WX-02-18** (33 mg, 47% yield) as a white solid.

**$^1$H-NMR** (400 MHz, $CD_3OD$): δ ppm 7.52 (d, J = 7.8 Hz, 1H), 7.29 (d, J = 8.0 Hz, 1H), 7.10 (t, J = 7.6 Hz, 1H), 7.03 (t, J = 7.7 Hz, 1H), 6.94 – 6.69 (m, 3H), 6.31 (brs, 1H), 5.94 (d, J = 3.8 Hz, 2H), 5.83 (brs, 1H), 3.63 – 3.09 (m, 8H), 2.95 (dd, J = 15.6, 6.2 Hz, 1H), 2.65 – 2.33 (m, 3H), 2.24 (t, J = 2.6 Hz, 1H), 2.12 – 1.78 (m, 4H), 1.74 – 1.55 (m, 2H). 1 exchangeable proton not observed.

**HRMS ESI-TOF** m/z calculated for $C_{31}H_{32}N_5O_5$ $[M+H]^+$ 554.2398. Found 554.2396.

**1-((1R,3R)-1-(benzo[d][1,3]dioxol-5-yl)-3-(morpholine-4-carbonyl)-1,3,4,9-tetrahydro-2H-pyrido[3,4-b]indol-2-yl)-3-(3-(but-3-yn-1-yl)-3H-diazirin-3-yl)propan-1-one (WX-02-38) (14)**

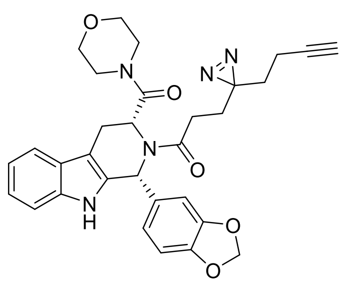

Prepared in an analogous fashion from *ent*-**S18**[3]. 14% yield as a white solid.

**$^1$H-NMR** (400 MHz, $CD_3OD$): δ ppm 7.53 (d, J = 7.7 Hz, 1H), 7.30 (d, J = 8.0 Hz, 1H), 7.12 – 7.01 (m, 2H), 6.87 – 6.74 (m, 3H), 6.31 (brs, 1H), 5.96 – 5.92 (m, 2H), 5.83 (brs, 1H), 3.61 – 3.31 (m, 8H), 2.96 (dd, J = 15.5, 6.3 Hz, 1H), 2.64 – 2.33 (m, 3H), 2.24 (t, J = 2.7 Hz, 1H), 2.09 – 1.78 (m, 4H), 1.72 – 1.55 (m, 2H). 1 exchangeable proton not observed.

**HRMS ESI-TOF** m/z calculated for $C_{31}H_{32}N_5O_5$ $[M+H]^+$ 554.2398. Found 554.2402.

**Synthesis of WX-02-28 and WX-02-48**

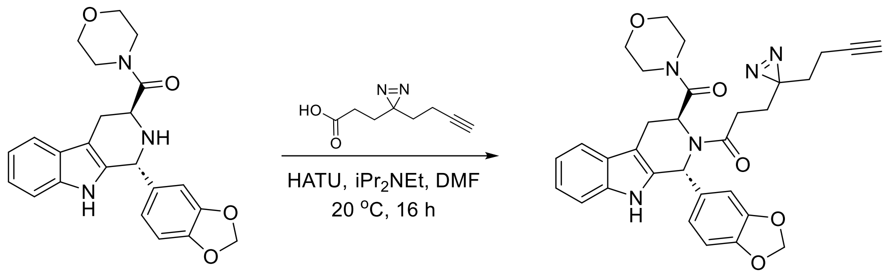


**S24** **WX-02-28 (15)**

**1-((1R,3S)-1-(benzo[d][1,3]dioxol-5-yl)-3-(morpholine-4-carbonyl)-1,3,4,9-tetrahydro-2H-pyrido[3,4-b]indol-2-yl)-3-(3-(but-3-yn-1-yl)-3H-diazirin-3-yl)propan-1-one (WX-02-28) (15)**

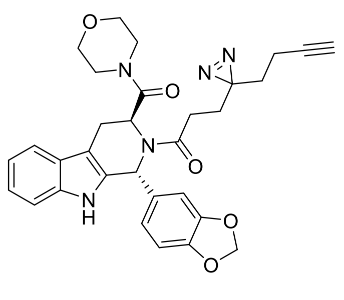

To a solution of **S24**[4] (70 mg, 0.17 mmol) in DMF (1 mL) was added 3-(3-(but-3-yn-1-yl)-3H-diazirin-3-yl)propanoic acid (29 mg, 0.17 mmol), $iPr_2NEt$ (67 mg, 0.52 mmol) and HATU (99 mg, 0.26 mmol). The mixture was stirred at 20 °C for 16 hours. Upon completion, the reaction mixture was diluted with water (20 mL) and extracted with EtOAc (20 mL×3). The organic layer was dried over $Na_2SO_4$, filtered, and concentrated in vacuo. The residue was purified by prep-HPLC (column: Waters Xbridge 150 mm × 25 mm × 5 µm; mobile phase: [A: water (10 mM ammonium bicarbonate)–B: MeCN]; B%: 41% – 71%, 9 min) to obtain **WX-02-28** (15 mg, 16% yield) as an off-white solid.

**$^1$H-NMR** (400 MHz, $CD_3OD$): δ ppm 7.75 (d, J = 7.8 Hz, 1H), 7.57 (d, J = 8.0 Hz, 1H), 7.40 – 7.34 (m, 1H), 7.34 – 7.27 (m, 1H), 7.22 – 7.15 (m, 2H), 7.14 – 7.06 (m, 1H), 6.55 (brs, 1H), 6.22

(d, J = 3.4 Hz, 2H), 5.51 (t, J = 5.5 Hz, 1H), 4.00 – 3.81 (m, 4H), 3.80 – 3.65 (m, 4H), 3.60 – 3.52 (m, 2H), 2.75 – 2.50 (m, 2H), 2.49 (t, J = 2.7 Hz, 1H), 2.25 (td, J = 7.5, 2.7 Hz, 2H), 2.00 (q, J = 7.5 Hz, 2H), 1.83 (t, J = 7.5 Hz, 2H). 1 exchangeable proton not observed.

**HRMS ESI-TOF** m/z calculated for $C_{31}H_{32}N_5O_5$ $[M+H]^+$ 554.2398. Found 554.2400.

**1-((1S,3R)-1-(benzo[d][1,3]dioxol-5-yl)-3-(morpholine-4-carbonyl)-1,3,4,9-tetrahydro-2H-pyrido[3,4-b]indol-2-yl)-3-(3-(but-3-yn-1-yl)-3H-diazirin-3-yl)propan-1-one (WX-02-48) (16)**

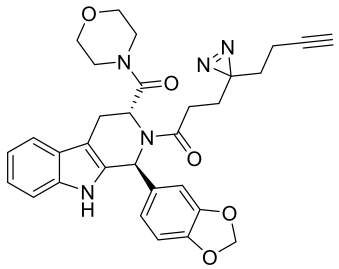

Prepared in an analogous fashion from *ent*-**S24**[4]. 13% yield as a yellow solid.

**$^1$H-NMR** (400 MHz, $CD_3OD$): δ ppm 7.75 (dd, J = 7.7, 1.3 Hz, 1H), 7.57 (d, J = 8.0 Hz, 1H), 7.37 (ddd, J = 8.1, 7.0, 1.3 Hz, 1H), 7.31 (td, J = 7.5, 1.2 Hz, 1H), 7.24 – 7.15 (m, 2H), 7.15 – 7.07 (m, 1H), 6.55 (brs, 1H), 6.23 (d, J = 3.2 Hz, 2H), 5.51 (t, J = 5.5 Hz, 1H), 3.97 – 3.82 (m, 4H), 3.80 – 3.66 (m, 4H), 3.60 – 3.52 (m, 2H), 2.76 – 2.49 (m, 2H), 2.49 (t, J = 2.7 Hz, 1H), 2.25 (td, J = 7.5, 2.5 Hz, 2H), 2.10 – 1.89 (m, 2H), 1.83 (t, J = 7.4 Hz, 2H). 1 exchangeable proton not observed.

**HRMS ESI-TOF** m/z calculated for $C_{31}H_{32}N_5O_5$ $[M+H]^+$ 554.2398. Found 554.2407.

**Synthesis of WX-02-22 and WX-02-42**

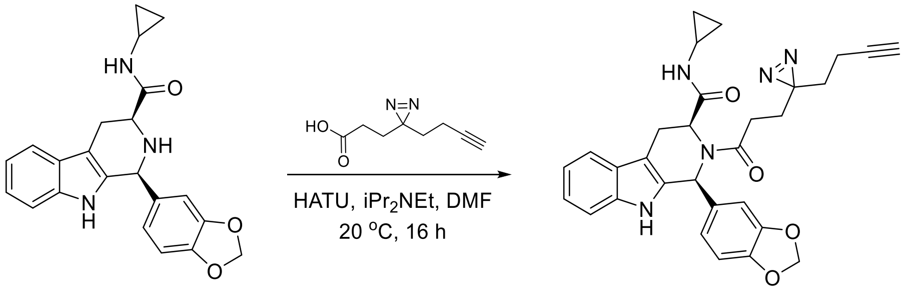


**S19** **WX-02-22 (17)**

**(1S,3S)-2-(3-(3-(but-3-yn-1-yl)-3H-diazirin-3-yl)propanoyl)-1-(benzo[d][1,3]dioxol-5-yl)-N-cyclopropyl-2,3,4,9-tetrahydro-1H-pyrido[3,4-b]indole-3-carboxamide (WX-02-22) (17)**

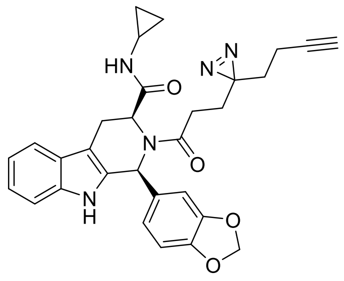

To a solution of **S18**[3] (50 mg, 0.12 mmol) in DMF (1 mL) were added 3-(3-(but-3-yn-1-yl)-3H-diazirin-3-yl)propanoic acid (20 mg, 0.12 mmol), $iPr_2NEt$ (23 mg, 0.18 mmol) and HATU (55 mg, 0.14 mmol). The mixture was stirred at 20 °C for 16 hours. Upon completion, the reaction mixture was diluted with water (30 mL) and extracted with EtOAc (40 mL×3). The organic layer was dried over $Na_2SO_4$, filtered, and concentrated in vacuo. The residue was purified by prep-HPLC (column: Waters Xbridge 150 mm × 25 mm × 5 µm; mobile phase: [A: water (10 mM ammonium bicarbonate)–B: MeCN]; B%: 43% – 76%, 9 min) to obtain **WX-02-22** (27 mg, 38% yield) as a yellow solid.

**$^1$H-NMR** (400 MHz, $CD_3OD$): δ ppm 7.81 (d, J = 7.8 Hz, 1H), 7.58 (d, J = 8.0 Hz, 1H), 7.42 – 7.36 (m, 1H), 7.36 – 7.29 (m, 1H), 7.21 – 7.07 (m, 2H), 7.03 (d, J = 8.0 Hz, 1H), 6.20 (s, 2H), 5.42 – 5.25 (m, 1H), 3.94 – 3.75 (m, 1H), 3.36 – 3.27 (m, 1H), 2.76 – 2.65 (m, 2H), 2.62 – 2.51 (m, 1H), 2.50 (t, J = 2.7 Hz, 1H), 2.36 – 2.28 (m, 2H), 2.26 – 2.08 (m, 2H), 1.96 – 1.88 (m, 2H), 0.91 – 0.69 (m, 2H), 0.59 – 0.45 (m, 2H). 2 exchangeable protons not observed.

**HRMS ESI-TOF** m/z calculated for $C_{30}H_{30}N_5O_4$ $[M+H]^+$ 524.2292. Found 524.2291.

**(1R,3R)-2-(3-(3-(but-3-yn-1-yl)-3H-diazirin-3-yl)propanoyl)-1-(benzo[d][1,3]dioxol-5-yl)-N-cyclopropyl-2,3,4,9-tetrahydro-1H-pyrido[3,4-b]indole-3-carboxamide (WX-02-42) (18)**

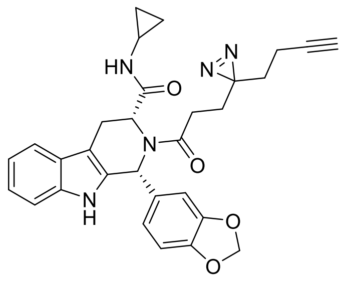

Prepared in an analogous fashion from *ent*-**S18**[3]. 42% yield as a yellow solid.

**$^1$H-NMR** (400 MHz, $CD_3OD$): δ ppm 7.81 (d, J = 7.8 Hz, 1H), 7.58 (d, J = 8.0 Hz, 1H), 7.39 (t, J = 7.6 Hz, 1H), 7.33 (t, J = 7.4 Hz, 1H), 7.23-7.08 (m, 2H), 7.04 (d, J = 8.0 Hz, 1H), 6.21 (s, 2H), 5.42 – 5.26 (m, 1H), 3.94 – 3.76 (m, 1H), 3.37 – 3.27 (m, 1H), 2.92 – 2.64 (m, 2H), 2.64 – 2.46 (m, 2H), 2.32 (td, J = 7.3, 2.6 Hz, 2H), 2.26 – 2.06 (m, 2H), 1.93 (t, J = 7.4 Hz, 2H), 0.89 – 071 (m, 2H), 0.58 – 0.42 (m, 2H). 2 exchangeable protons not observed.

**HRMS ESI-TOF** m/z calculated for $C_{30}H_{30}N_5O_4$ $[M+H]^+$ 524.2292. Found 524.2289.

**Synthesis of WX-02-20**

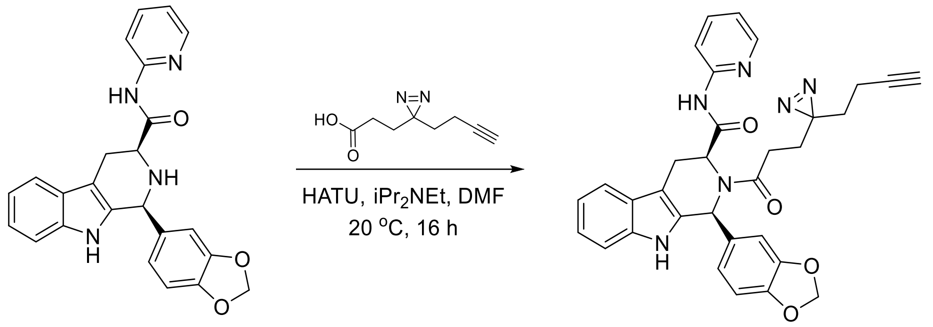



**S20** **WX-02-20 (19)**

**(1S,3S)-2-(3-(3-(but-3-yn-1-yl)-3H-diazirin-3-yl)propanoyl)-1-(benzo[d][1,3]dioxol-5-yl)-N-(pyridine-2-yl)-2,3,4,9-tetrahydro-1H-pyrido[3,4-b]indole-3-carboxamide (WX-02-20) (19)**

To a solution of **S20**[3] (50 mg, 0.12 mmol) in DMF (1 mL) were added 3-(3-(but-3-yn-1-yl)-3H-diazirin-3-yl)propanoic acid (20 mg, 0.12 mmol), $iPr_2NEt$ (23 mg, 0.18 mmol) and HATU (55 mg, 0.14 mmol). The mixture was stirred at 20 °C for 16 hours. Upon completion, the reaction mixture was diluted with water (30 mL) and extracted with EtOAc (40 mL×3). The organic layer was dried over $Na_2SO_4$, filtered, and concentrated in vacuo. The residue was purified by prep-HPLC (column: Waters Xbridge 150 mm × 25 mm × 5 μm; mobile phase: [A: water (10 mM ammonium bicarbonate)–B: MeCN]; B%: 44% – 74%, 10 min) to obtain **WX-02-20** (25 mg, 35% yield) as a yellow solid.

**$^1$H-NMR** (400 MHz, $CD_3OD$): δ ppm 8.39 – 8.34 (m, 1H), 8.10 – 8.03 (m, 1H), 7.94 – 7.85 (m, 1H), 7.83 (d, J = 7.7 Hz, 1H), 7.56 (d, J = 8.0 Hz, 1H), 7.42 – 7.23 (m, 4H), 6.97 (s, 2H), 6.65 (s, 1H), 5.97 – 5.80 (m, 2H), 5.58 (brs, 1H), 4.04 – 3.82 (m, 1H), 3.37 – 3.28 (m, 1H), 3.00 – 2.76 (m, 2H), 2.48 (t, J = 2.7 Hz, 1H), 2.30 (qd, J = 7.6, 3.9 Hz, 2H), 2.20 (t, J = 7.4 Hz, 2H), 1.94 (t, J = 7.5 Hz, 2H). 2 exchangeable protons not observed.

**HRMS ESI-TOF** m/z calculated for $C_{32}H_{29}N_6O_4$ $[M+H]^+$ 561.2250. Found 561.2239.

**Synthesis of WX-02-21**

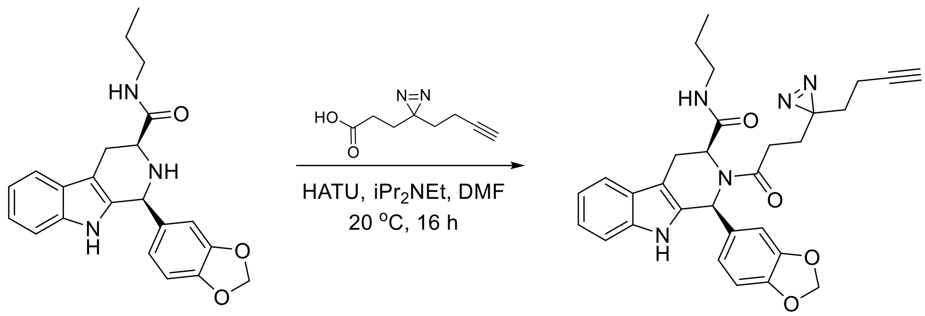

S21

WX-02-21 (20)

**(1S,3S)-2-(3-(3-(but-3-yn-1-yl)-3H-diazirin-3-yl)propanoyl)-1-(benzo[d][1,3]dioxol-5-yl)-N-propyl-2,3,4,9-tetrahydro-1H-pyrido[3,4-b]indole-3-carboxamide (WX-02-21) (20)**

To a solution of **S21**[3] (50 mg, 0.12 mmol) in DMF (1 mL) were added 3-(3-(but-3-yn-1-yl)-3H-diazirin-3-yl)propanoic acid (20 mg, 0.12 mmol), iPr$_2$NEt (23 mg, 0.18 mmol) and HATU (55 mg, 0.14 mmol). The mixture was stirred at 20 °C for 16 hours. Upon completion, the reaction mixture was diluted with water (30 mL) and extracted with EtOAc (40 mL×3). The organic layer was dried over $Na_2SO_4$, filtered, and concentrated in vacuo. The residue was purified by prep-HPLC (column: Waters Xbridge 150 mm × 25 mm × 5 µm; mobile phase: [A: water (10 mM ammonium bicarbonate)–B: MeCN]; B%: 43% – 76%, 10 min) to obtain **WX-02-21** (21 mg, 29% yield) as a yellow solid.

**$^1$H-NMR** (400 MHz, $CD_3OD$): δ ppm 7.52 (d, J = 7.8 Hz, 1H), 7.27 (d, J = 8.0 Hz, 1H), 7.09 (t, J = 7.5 Hz, 1H), 7.06 – 6.91 (m, 2H), 6.94 – 6.64 (m, 3H), 5.90 (s, 2H), 5.08 – 4.93 (m, 1H), 3.69 – 3.52 (m, 1H), 3.01 (dd, J = 15.9, 6.5 Hz, 1H), 2.95 – 2.77 (m, 1H), 2.51 – 2.33 (m, 3H), 2.25 (t, J = 2.7 Hz, 1H), 2.12 – 1.53 (m, 6H), 1.43 – 1.09 (m, 2H), 0.85 – 0.70 (m, 3H). 2 exchangeable protons not observed.

**HRMS ESI-TOF** m/z calculated for $C_{30}H_{32}N_5O_4$ $[M+H]^+$ 526.2454. Found 526.2455.

**Synthesis of WX-02-19**

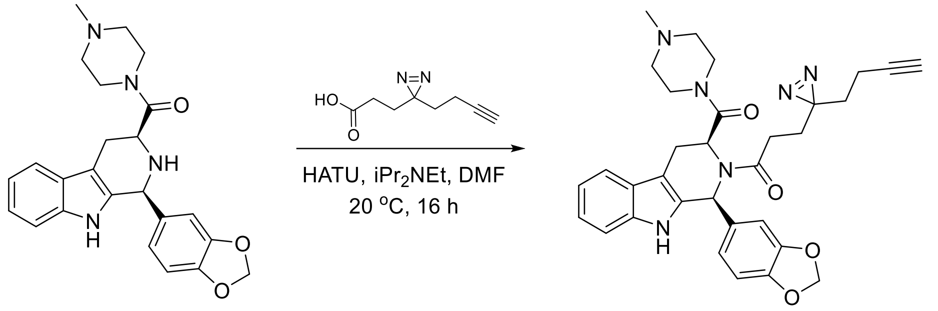


**1-((1S,3S)-1-(benzo[d][1,3]dioxol-5-yl)-3-( 4-methylpiperazine-1-carbonyl)-1,3,4,9-tetrahydro-2H-pyrido[3,4-b]indol-2-yl)-3-(3-(but-3-yn-1-yl)-3H-diazirin-3-yl)propan-1-one (WX-02-19) (21)**

To a solution of **S22**[3] (50 mg, 0.12 mmol) in DMF (1 mL) were added 3-(3-(but-3-yn-1-yl)-3H-diazirin-3-yl)propanoic acid (20 mg, 0.12 mmol), iPr$_2$NEt (23 mg, 0.18 mmol) and HATU (55 mg, 0.14 mmol). The mixture was stirred at 20 °C for 16 hours. Upon completion, the reaction mixture was diluted with water (30 mL) and extracted with EtOAc (40 mL×3). The organic layer was dried over $Na_2SO_4$, filtered, and concentrated in vacuo. The residue was purified by prep-HPLC (column: Phenomenex Synergi C18 150 mm × 25 mm × 10 µm; mobile phase: [A: water (10 mM 0.23% formic acid)–B: MeCN]; B%: 15% – 45%, 10 min) to obtain **WX-02-21** (23 mg, 32% yield) as an off-white solid.

**$^1$H-NMR** (400 MHz, $CD_3OD$): δ ppm 7.51 (d, J = 7.8 Hz, 1H), 7.29 (d, J = 8.0 Hz, 1H), 7.10 (t, J = 7.5 Hz, 1H), 7.03 (t, J = 7.4 Hz, 1H), 6.85 – 6.70 (m, 3H), 6.30 (brs, 1H), 5.95 (d, J = 8.8 Hz, 2H), 5.85 – 5.78 (brs, 1H), 3.64 – 3.43 (m, 2H), 3.37 – 3.27 (m, 2H), 2.95 (dd, J = 15.7, 6.2 Hz, 1H), 2.76 – 2.17 (m, 11H), 2.11 – 1.90 (m, 3H), 1.91 – 1.78 (m, 1H), 1.71 – 1.56 (m, 2H). 1 exchangeable proton not observed.

**HRMS ESI-TOF** m/z calculated for $C_{32}H_{35}N_6O_4$ $[M+H]^+$ 567.2720. Found 567.2723.

**Synthesis of WX-02-221**

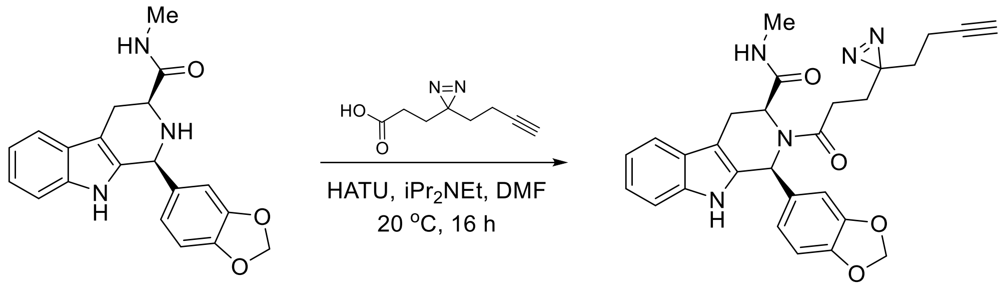

S23

WX-02-221 (22)

**(1S,3S)-2-(3-(3-(but-3-yn-1-yl)-3H-diazirin-3-yl)propanoyl)-1-(benzo[d][1,3]dioxol-5-yl)-N-methyl-2,3,4,9-tetrahydro-1H-pyrido[3,4-b]indole-3-carboxamide (WX-02-221) (22)**

To a solution of **S23**[3] (80 mg, 0.23 mmol) in DMF (1 mL) were added 3-(3-(but-3-yn-1-yl)-3H-diazirin-3-yl)propanoic acid (38 mg, 0.23 mmol), $iPr_2NEt$ (44 mg, 0.34 mmol) and HATU (113 mg, 0.30 mmol). The mixture was stirred at 20 °C for 16 hours. Upon completion, the reaction mixture was diluted with water (30 mL) and extracted with EtOAc (40 mL×3). The organic layer was dried over $Na_2SO_4$, filtered, and concentrated in vacuo. The residue was purified by prep-HPLC (column: Phenomenex Synergi C18 150 mm × 25 mm × 10 µm; mobile phase: [A: water (10 mM 0.23% formic acid)–B: MeCN]; B%: 41% – 71%, 10 min) to obtain **WX-02-221** (19 mg, 16% yield) as a yellow solid.

**$^1$H-NMR** (400 MHz, $CD_3OD$): δ ppm 7.53 (d, J = 7.8 Hz, 1H), 7.26 (d, J = 8.0 Hz, 1H), 7.12 – 6.96 (m, 3H), 6.85 – 6.60 (m, 3H), 5.89 (s, 2H), 5.07 – 4.96 (m, 1H), 3.72 – 3.60 (m, 1H), 2.98 (dd, J = 15.9, 6.7 Hz, 1H), 2.58 – 2.30 (m, 3H), 2.32 – 2.14 (m, 3H), 2.09 – 1.97 (m, 2H), 1.92 – 1.81 (s, 2H), 1.72 – 1.56 (m, 2H). 2 exchangeable protons not observed.

**HRMS ESI-TOF** m/z calculated for $C_{28}H_{28}N_5O_4$ $[M+H]^+$ 498.2141. Found 498.2134.

#### Synthesis of WX-03-93, WX-03-95, WX-03-97, and WX-03-99

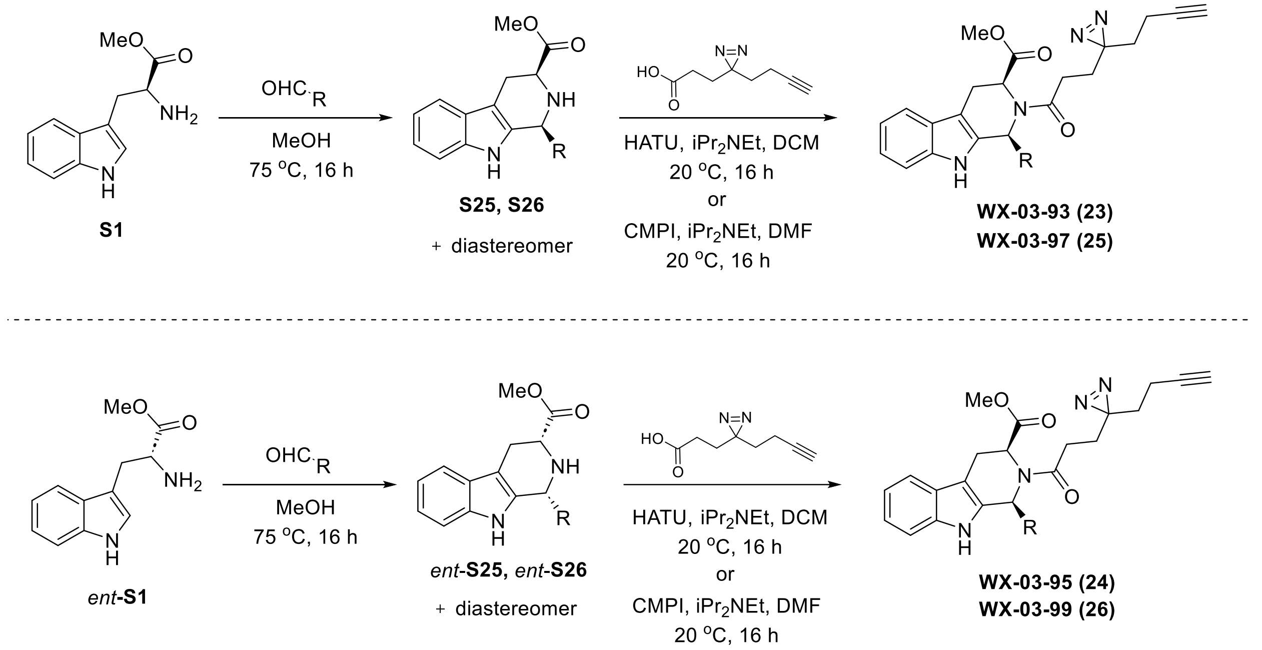


#### Synthesis of WX-03-93 and WX-03-95

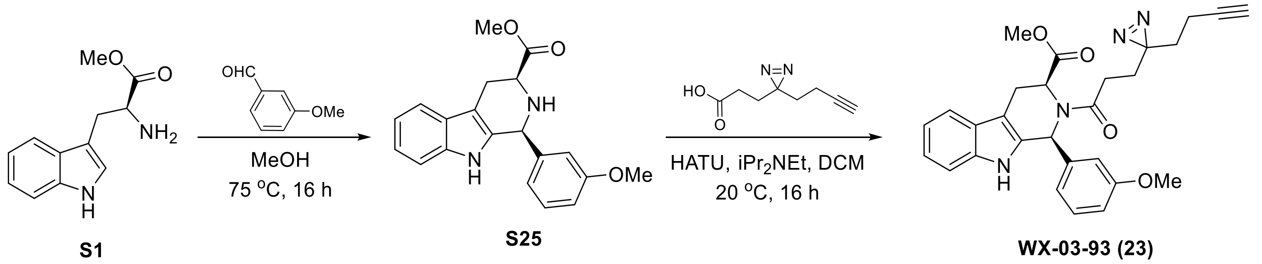


#### methyl (1S,3S)-1-(3-methoxyphenyl)-2,3,4,9-tetrahydro-1H-pyrido[3,4-b]indole-3-carboxylate (S25)

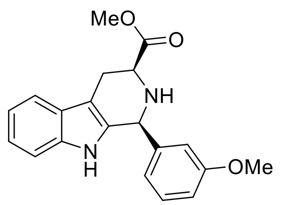

A solution of **S1** (2.0 g, 7.9 mmol) and 3-methoxybenzaldehyde (1.1 g, 7.9 mmol) in MeOH (30 mL) was stirred at 80 °C for 12 hours. Upon completion, the reaction mixture was concentrated in vacuo. The residue was diluted with EtOAc (200 mL) and washed with $NaHCO_3$ aqueous (100 mL). The organic layer was dried over $Na_2SO_4$, filtered, and concentrated in vacuo. The residue was purified by column chromatography ($SiO_2$, Petroleum ether/Ethyl acetate=10/1 to 1/1) to give **S25** (1.2 g, 46% yield) as a yellow amorphous.

**$^1$H-NMR** (400 MHz, $CDCl_3$): δ ppm 7.58 – 7.52 (m, 1H), 7.48 (brs, 1H), 7.30 (t, J = 8.0 Hz, 1H), 7.25 – 7.20 (m, 1H), 7.18 – 7.10 (m, 2H), 6.99 (d, J = 7.6 Hz, 1H), 6.95 (d, J = 2.4 Hz, 1H), 6.91

(dd, J = 2.4, 8.0 Hz, 1H), 5.24 (s, 1H), 3.99 (dd, J = 4.0, 11.2 Hz, 1H), 3.83 (s, 3H), 3.79 (s, 3H), 3.28 – 3.20 (m, 1H), 3.08 - 2.96 (m, 1H).

**LC-MS** m/z calculated for $C_{20}H_{21}N_2O_3$ $[M+H]^+$ 337.2. Found 337.1.

**methyl (1S,3S)-2-(3-(3-(but-3-yn-1-yl)-3H-diazirin-3-yl)propanoyl)-1-(3-methoxyphenyl)-2,3,4,9-tetrahydro-1H-pyrido[3,4-b]indole-3-carboxylate (WX-03-93) (23)**

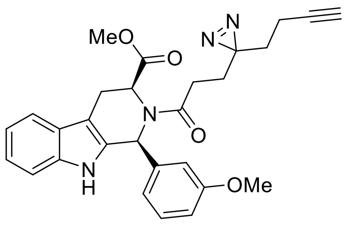


To a solution of **S25** (70 mg, 0.21 mmol) in DCM (1.5 mL) were added 3-(3-(but-3-yn-1-yl)-3H-diazirin-3-yl)propanoic acid (52 mg, 0.31 mmol), $iPr_2NEt$ (81 mg, 0.62 mmol) and HATU (120 mg, 0.31 mmol). The mixture was stirred at 20 °C for 12 hours. Upon completion, the reaction mixture was concentrated under reduced pressure to give a residue. The residue was purified by prep-TLC (Petroleum ether/Ethyl acetate =1/1) and followed by prep-HPLC (column: Phenomenex Gemini-NX C18 75 mm × 30 mm × 3 μm; mobile phase: [A: water (10 mM ammonium bicarbonate)–B: MeCN]; B%: 42% – 72%, 8 min) to obtain **WX-03-93** (60 mg, 60% yield) as an off-white solid.

**$^1$H-NMR** (400 MHz, $CD_3OD$): δ ppm 7.56 (d, J = 7.7 Hz, 1H), 7.31 (d, J = 8.0 Hz, 1H), 7.21 – 7.11 (m, 2H), 7.11 – 7.03 (m, 2H), 6.86 – 6.74 (m, 3H), 5.19 (d, J = 6.9 Hz, 1H), 3.72 (s, 3H), 3.63 (d, J = 15.9 Hz, 1H), 3.09 (dd, J = 15.9, 6.8 Hz, 1H), 3.03 (s, 3H), 2.51 (t, J = 7.3 Hz, 2H), 2.28 (t, J = 2.7 Hz, 1H), 2.07 (td, J = 7.5, 2.6 Hz, 2H), 1.91 (td, J = 7.2, 3.6 Hz, 2H), 1.70 (t, J = 7.5 Hz, 2H). 1 exchangeable proton not observed.

**HRMS ESI-TOF** m/z calculated for $C_{28}H_{29}N_4O_4$ $[M+H]^+$ 485.2184. Found 485.2192.

**methyl (1R,3R)-1-(3-methoxyphenyl)-2,3,4,9-tetrahydro-1H-pyrido[3,4-b]indole-3-carboxylate (*ent*-S25)**

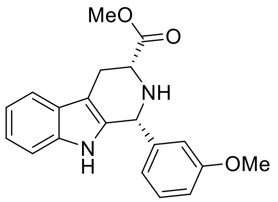


Prepared in an analogous fashion from *ent*-**S1**. 44% yield as an off-white solid.

**$^1$H-NMR** (400 MHz, $CDCl_3$): δ ppm 7.59 – 7.54 (m, 1H), 7.52 – 7.46 (m, 1H), 7.34 – 7.29 (m, 1H), 7.26 – 7.22 (m, 1H), 7.20 – 7.11 (m, 2H), 7.02 – 6.95 (m, 2H), 6.94 – 6.90 (m, 1H), 5.28 – 5.23 (m, 1H), 3.98 (dd, J = 11.2, 4.3 Hz, 1H), 3.85 – 3.84 (m, 3H), 3.82 – 3.79 (m, 3H), 3.29 – 3.22 (m, 1H), 3.09 – 2.99 (m, 1H).

**LC-MS** m/z calculated for $C_{20}H_{21}N_2O_3$ $[M+H]^+$ 337.2. Found 337.1.

**methyl (1R,3R)-2-(3-(3-(but-3-yn-1-yl)-3H-diazirin-3-yl)propanoyl)-1-(3-methoxyphenyl)-2,3,4,9-tetrahydro-1H-pyrido[3,4-b]indole-3-carboxylate (WX-03-95) (24)**

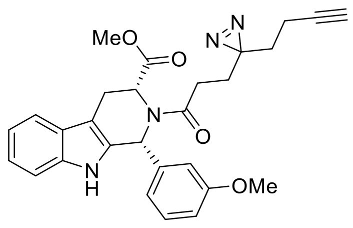

Prepared in an analogous fashion from *ent*-**S25**. 30% yield as a yellow solid.

**$^1$H-NMR** (400 MHz, $CD_3OD$): δ ppm 7.55 (d, J = 7.8 Hz, 1H), 7.30 (d, J = 8.0 Hz, 1H), 7.20 – 7.02 (m, 4H), 6.90 – 6.73 (m, 3H), 5.18 (d, J = 6.9 Hz, 1H), 3.71 (s, 3H), 3.61 (d, J = 16.0 Hz, 1H), 3.08 (dd, J = 15.9, 6.6 Hz, 1H), 3.01 (s, 3H), 2.50 (t, J = 7.3 Hz, 2H), 2.27 (t, J = 2.6 Hz, 1H), 2.06 (td, J = 7.5, 2.6 Hz, 2H), 1.89 (td, J = 7.2, 3.5 Hz, 2H), 1.68 (t, J = 7.5 Hz, 2H). 1 exchangeable proton not observed.

**HRMS ESI-TOF** m/z calculated for $C_{28}H_{29}N_4O_4$ $[M+H]^+$ 485.2184. Found 485.2192.

**Synthesis of WX-03-97 and WX-03-99**

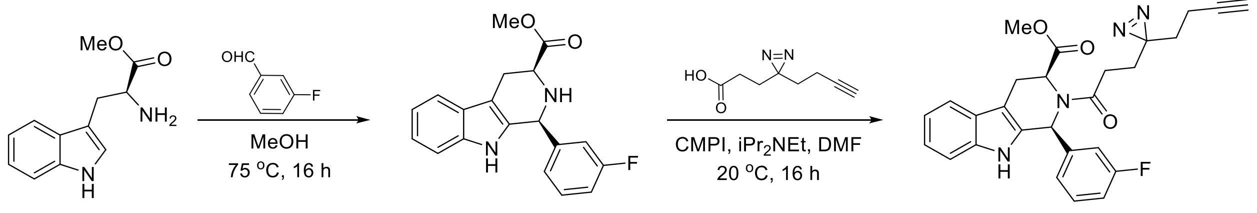


**methyl (1S,3S)-1-(3-fluorophenyl)-2,3,4,9-tetrahydro-1H-pyrido[3,4-b]indole-3-carboxylate (S26)**

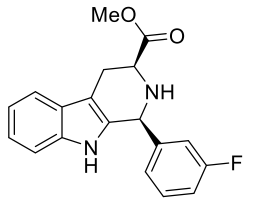

A solution of **S1** (5.0 g, 20 mmol) and 3-fluorobenzaldehyde (3.7 g, 29 mmol) in MeOH (60 mL) was stirred at 80 °C for 12 hours. Upon completion, the reaction mixture was concentrated in vacuo. The residue was diluted with EtOAc (200 mL) and washed with $NaHCO_3$ aqueous (100 mL). The organic layer was dried over $Na_2SO_4$, filtered, and concentrated in vacuo. The residue was purified by column chromatography ($SiO_2$, Petroleum ether/Ethyl acetate=10/1 to 1/1) to give **S26** (2.8 g, 44% yield) as a white solid.

**$^1$H-NMR** (400 MHz, $CDCl_3$): δ ppm 7.55 (d, J = 7.2 Hz, 1H), 7.44 (brs, 1H), 7.36 (dt, J = 6.0, 8.0 Hz, 1H), 7.23 (dd, J = 7.2, 14.8 Hz, 2H), 7.19 – 7.10 (m, 3H), 7.07 (dt, J = 2.0, 8.0 Hz, 1H), 5.28 (s, 1H), 3.98 (dd, J = 11.2, 4.2 Hz, 1H), 3.84 (s, 3H), 3.28 – 3.20 (m, 1H), 3.07 – 2.97 (m, 1H), 2.90 – 2.18 (m, 1H).

**LC-MS** m/z calculated for $C_{19}H_{18}FN_2O_3$ [M+H]$^+$ 325.1. Found 325.1.

**methyl (1S,3S)-2-(3-(3-(but-3-yn-1-yl)-3H-diazirin-3-yl)propanoyl)-1-(3-fluorophenyl)-2,3,4,9-tetrahydro-1H-pyrido[3,4-b]indole-3-carboxylate (WX-03-97) (25)**

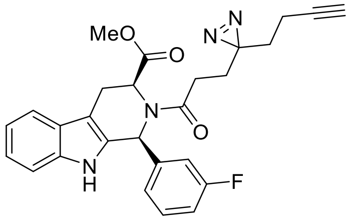

To a solution of **S26** (70 mg, 0.21 mmol) in DMF (1.0 mL) were added 3-(3-(but-3-yn-1-yl)-3H-diazirin-3-yl)propanoic acid (54 mg, 0.32 mmol), iPr$_2$NEt (84 mg, 0.65 mmol) and CMPI (110 mg, 0.43 mmol). The mixture was stirred at 20 °C for 12 hours. Upon completion, the reaction mixture was diluted with EtOAc (30 mL) and washed with water (20 mL x 3). The combined organic layers were dried over anhydrous sodium sulfate, filtered and concentrated under reduced pressure to give a residue. The residue was purified by prep-TLC (Petroleum ether/Ethyl acetate =1/1) and followed by prep-HPLC (column: Waters Xbridge 150 mm × 25 mm × 5 μm; mobile phase: [A: water (10 mM ammonium bicarbonate)–B: MeCN]; B%: 53% – 83%, 10 min) to obtain **WX-03-97** (60 mg, 59% yield) as a white solid.

**$^1$H-NMR** (400 MHz, CD$_3$OD): δ ppm 7.56 (d, J = 7.8 Hz, 1H), 7.34 – 7.24 (m, 2H), 7.17 – 7.11 (m, 1H), 7.11 – 7.03 (m, 3H), 7.03 – 6.94 (m, 2H), 5.22 (d, J = 6.9 Hz, 1H), 3.62 (d, J = 15.9 Hz, 1H), 3.14 – 3.06 (m, 1H), 3.05 (s, 3H), 2.51 (t, J = 7.2 Hz, 2H), 2.27 (t, J = 2.7 Hz, 1H), 2.06 (td, J = 7.4, 2.6 Hz, 2H), 1.90 (td, J = 7.3, 2.6 Hz, 2H), 1.69 (t, J = 7.4 Hz, 2H). 1 exchangeable proton not observed.

**HRMS ESI-TOF** m/z calculated for $C_{27}H_{26}N_4O_3$ [M+H]$^+$ 473.1984. Found 473.1990.

**methyl (1R,3R)-1-(3-fluorophenyl)-2,3,4,9-tetrahydro-1H-pyrido[3,4-b]indole-3-carboxylate (*ent*-S26)**

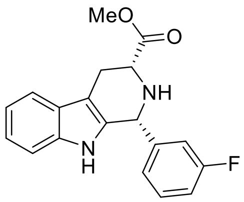

Prepared in an analogous fashion from *ent*-**S1**. 46% yield as an off-white solid.

**$^1$H-NMR** (400 MHz, CDCl$_3$): δ ppm 7.60 – 7.53 (m, 1H), 7.45 (brs, 1H), 7.40 – 7.32 (m, 1H), 7.26 – 7.10 (m, 5H), 7.10 – 7.03 (m, 1H), 5.28 (s, 1H), 3.97 (dd, J = 11.1, 4.2 Hz, 4H), 3.83 (s, 3H), 3.28 – 3.20 (m, 1H), 3.08 – 2.98 (m, 1H), 2.53 (brs, 1H).

**LC-MS** m/z calculated for $C_{19}H_{18}FN_2O_3$ [M+H]$^+$ 325.1. Found 325.1.

**methyl (1R,3R)-2-(3-(3-(but-3-yn-1-yl)-3H-diazirin-3-yl)propanoyl)-1-(3-fluorophenyl)-2,3,4,9-tetrahydro-1H-pyrido[3,4-b]indole-3-carboxylate (WX-03-99) (26)**

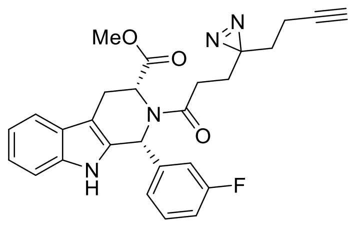

Prepared in an analogous fashion from *ent*-**S26**. 43% yield as a white solid.

**[1]H-NMR** (400 MHz, $CD_3OD$): δ ppm 7.56 (d, J = 7.8 Hz, 1H), 7.36 – 7.23 (m, 2H), 7.17 – 7.11 (m, 1H), 7.11 – 7.03 (m, 3H), 7.02 – 6.95 (m, 2H), 5.20 (d, J = 6.8 Hz, 1H), 3.61 (d, J = 15.9 Hz, 1H), 3.13 – 3.05 (m, 1H), 3.05 (s, 3H), 2.50 (t, J = 7.3 Hz, 2H), 2.26 (d, J = 2.7 Hz, 1H), 2.06 (td, J = 7.5, 2.6 Hz, 2H), 1.89 (td, J = 7.4, 2.8 Hz, 2H), 1.68 (t, J = 7.8 Hz, 2H). 1 exchangeable proton not observed.

**HRMS ESI-TOF** m/z calculated for $C_{27}H_{26}N_4O_3$ $[M+H]^+$ 473.1984. Found 473.1987.

**Synthesis of DBK-073A and DBK-073B**

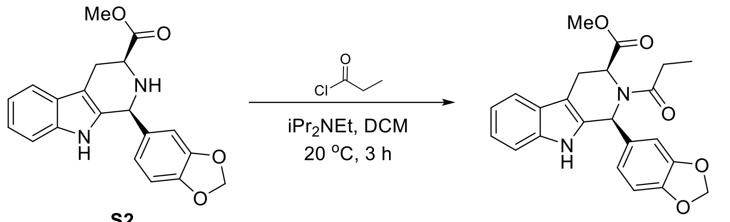


**methyl (1S,3S)-2-propanoyl-1-(benzo[d][1,3]dioxol-5-yl)-2,3,4,9-tetrahydro-1H-pyrido[3,4-b]indole-3-carboxylate (DBK-073A) (27)**

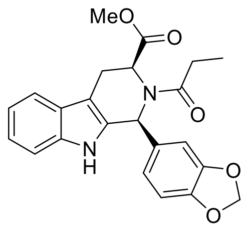

To a solution of **S2**[1] (35 mg, 0.10 mmol) in EtOAc (1.0 mL) were added propionyl chloride (12 mg, 0.13 mmol) and $K_2CO_3$ (42 mg, 0.30 mmol) at 0 °C. The reaction was warmed to 20 °C and stirred for 2 hours. Upon completion, the resulting mixture was diluted with EtOAc and washed with sat.$NaHCO_3$ (x1). The organic layer was dried over $Na_2SO_4$, filtered, and concentrated in vacuo. The residue was purified via prep-TLC (Hexane/EtOAc =3/2) to obtain **DBK-073A** (21 mg, 51% yield) as an off-white amorphous.

**[1]H-NMR** (600 MHz, $CDCl_3$): δ ppm 7.84 (brs, 1H), 7.59 (d, J = 7.7 Hz, 1H), 7.29 (d, J = 8.0 Hz, 1H), 7.20 (t, J = 7.5 Hz, 1H), 7.15 (t, J = 7.4 Hz, 1H), 6.97 (brs, 1H), 6.92 (brs, 1H), 6.70 – 6.58 (m, 2H), 5.90 (s, 2H), 4.93 (brs, 1H), 3.67 (d, J = 15.8 Hz, 1H), 3.16 (s, 3H), 3.05 (dd, J = 15.6, 7.1 Hz, 1H), 2.64 – 2.47 (m, 2H), 1.24 (t, J = 6.8 Hz, 3H).

**HRMS ESI-TOF** m/z calculated for $C_{23}H_{23}N_2O_5$ $[M+H]^+$ 407.1602. Found 407.1602.

**methyl (1R,3R)-2-propanoyl-1-(benzo[d][1,3]dioxol-5-yl)-2,3,4,9-tetrahydro-1H-pyrido[3,4-b]indole-3-carboxylate (DBK-073B) (28)**

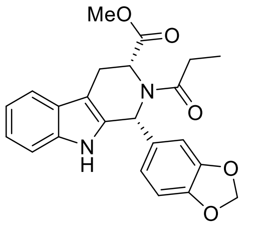

Prepared in an analogous fashion from *ent*-**S2**[1]. 83% yield as an off-white amorphous.

**$^1$H-NMR** (600 MHz, $CDCl_3$): δ ppm 7.87 (brs, 1H), 7.59 (d, J = 7.8 Hz, 1H), 7.29 (d, J = 7.9 Hz, 1H), 7.20 (ddd, J = 8.1, 7.0, 1.3 Hz, 1H), 7.15 (ddd, J = 8.0, 7.1, 1.1 Hz, 1H), 6.97 (brs, 1H), 6.92 (brs, 1H), 6.69 – 6.57 (m, 2H), 5.90 (s, 2H), 4.93 (brs, 1H), 3.67 (d, J = 15.8 Hz, 1H), 3.16 (s, 3H), 3.05 (ddd, J = 15.7, 6.9, 1.8 Hz, 1H), 2.64 – 2.47 (m, 2H), 1.25 (t, J = 7.1 Hz, 3H).

**HRMS ESI-TOF** m/z calculated for $C_{23}H_{23}N_2O_5$ $[M+H]^+$ 407.1602. Found 407.1608.

**Synthesis of DO-40A and DO-40B**

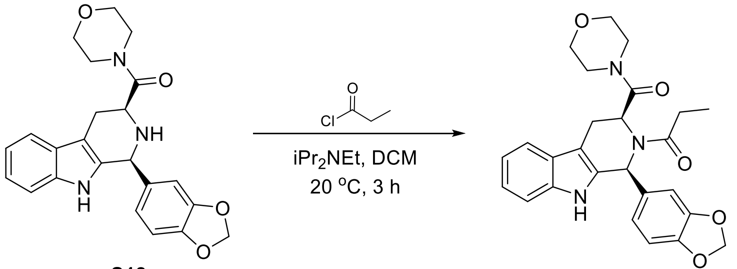


**S18** **DO-40A (29)**

**1-((1S,3S)-1-(benzo[d][1,3]dioxol-5-yl)-3-(morpholine-4-carbonyl)-1,3,4,9-tetrahydro-2H-pyrido[3,4-b]indol-2-yl)-propan-1-one (DO-40A) (29)**

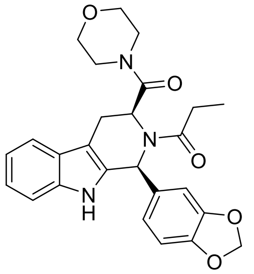

To a solution of **S17**[3] (15 mg, 0.037 mmol) in DCM (0.53 mL) were added propionyl chloride (3.9 μL, 0.044 mmol) and $iPr_2NEt$ (9.7 μL, 0.055 mmol). The mixture was stirred at 20 °C for 3 hours. Upon completion, the resulting mixture was diluted with DCM and washed with sat.$NaHCO_3$ (x1). The organic layer was dried over $Na_2SO_4$, filtered, and concentrated in vacuo. The residue was purified via prep-TLC (Hexane/EtOAc =1/1) to obtain **DO-40A** (11 mg, 65% yield) as an off-white amorphous.

**$^1$H-NMR** (600 MHz, $CDCl_3$): δ ppm 7.74 (brs, 1H), 7.59 (d, J = 7.7 Hz, 1H), 7.27 – 7.24 (m, 1H, overlapped with solvent peak), 7.20 – 7.15 (m, 1H), 7.13 (t, J = 7.4 Hz, 1H), 6.80 (brs, 1H), 6.77

– 6.71 (m, 2H), 6.14 (brs, 1H), 5.98 – 5.91 (m, 3H), 3.57 – 3.30 (m, 7H), 3.04 – 2.87 (m, 2H), 2.70 (brs, 2H), 2.44 (brs, 1H), 1.23 (t, J = 7.3 Hz, 4H).

**HRMS ESI-TOF** m/z calculated for $C_{26}H_{28}N_3O_5$ $[M+H]^+$ 462.2024. Found 462.2037.

**1-((1S,3S)-1-(benzo[d][1,3]dioxol-5-yl)-3-(morpholine-4-carbonyl)-1,3,4,9-tetrahydro-2H-pyrido[3,4-b]indol-2-yl)-propan-1-one (DO-40B) (30)**

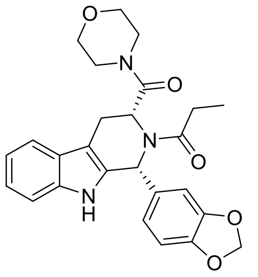

Prepared in an analogous fashion from *ent*-**S17**[3]. 62% yield as an off-white amorphous.

**$^1$H-NMR** (600 MHz, $CDCl_3$): δ ppm 7.69 (brs, 1H), 7.60 (d, J = 7.8 Hz, 1H), 7.29 – 7.25 (m, 1H, overlapped with solvent peak), 7.20 – 7.16 (m, 1H), 7.14 (t, J = 7.3 Hz, 1H), 6.81 (brs, 1H), 6.78 – 6.72 (m, 2H), 6.19 (brs, 1H), 5.97 – 5.93 (m, 3H), 3.58 – 3.31 (m, 7H), 3.03 – 2.89 (m, 2H), 2.70 (brs, 2H), 2.44 (brs, 1H), 1.23 (t, J = 7.2 Hz, 4H).

**HRMS ESI-TOF** m/z calculated for $C_{26}H_{28}N_3O_5$ $[M+H]^+$ 462.2024. Found 462.2024.

#### Synthesis of DO-35A and DO-35B

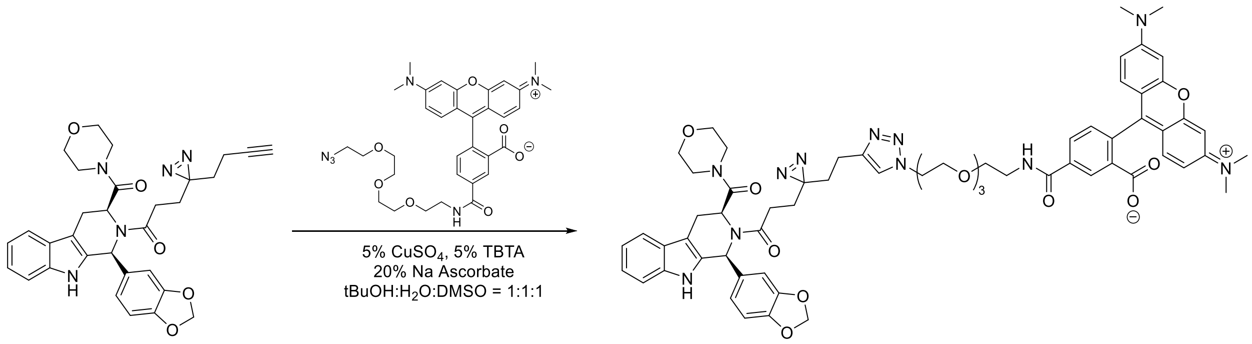


**WX-02-18 (13)** **DO-35A (31)**

**5-[2-[2-[2-(2-(4-(2-(3-(3-((1S,3S)-1-(benzo[d][1,3]dioxol-5-yl)-3-(morpholine-4-carbonyl)-1,3,4,9-tetrahydro-2H-pyrido[3,4-b]indol-2-yl)-3-oxopropyl)-3H-diazirin-3-yl)ethyl)-1H-1,2,3-triazol-1-yl)ethoxy)ethoxy]ethoxy]ethylcarbamoyl]-2-(3,6-Bis(dimethylamino)xanthylium-9-yl)benzoate (DO-35A) (31)**

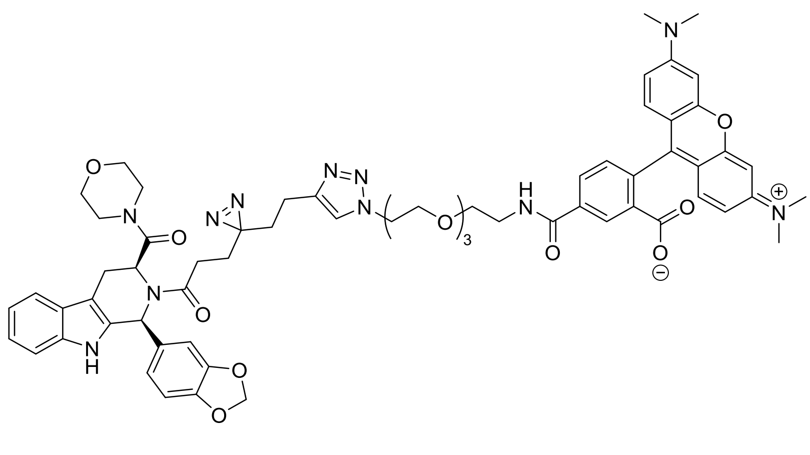

To a solution of **WX-02-18** (4.4 mg, 0.0079 mmol) in H2O/t-BuOH/DMSO (0.2 mL/0.2 mL/0.1 mL) were added TAMRA-(PEG)$_3$-N$_3$ (5.0 mg, 0.0079 mmol), TBTA (0.2 mg, 0.4 μmol), sodium ascorbate (0.3 mg, 1.6 μmol) and $CuSO_4$ (0.2 mg, 0.4 μmol). The mixture was stirred at 20 °C for 12 hours. Upon completion, the resulting mixture was purified by prep-HPLC (column: Waters BEH C18 160 mm × 19 mm × 5 μm; mobile phase: [water (0.1% formic acid)-MeCN]; B%: 35% – 55%, 8 min) to obtain **DO-35A** (5.6 mg, 60% yield) as a red solid.

**$^1$H-NMR** (600 MHz, $CDCl_3$): δ ppm 8.95 (s, 1H), 8.44 (s, 1H), 8.20 (dd, J = 8.0, 1.5 Hz, 1H), 7.57 (brs, 1H), 7.55 – 7.50 (m, 2H), 7.37 – 7.31 (m, 1H), 7.21 (d, J = 8.0 Hz, 1H), 7.19 – 7.14 (m, 1H), 7.12 – 7.03 (m, 2H), 6.78 (d, J = 8.9 Hz, 2H), 6.71 – 6.64 (m, 1H), 6.61 (d, J = 9.0 Hz, 1H), 6.50 (dd, J = 18.4, 2.6 Hz, 2H), 6.44 (d, J = 9.0 Hz, 1H), 6.37 (d, J = 9.1 Hz, 1H), 6.25 (brs, 1H), 5.86 (d, J = 4.6 Hz, 2H), 5.71 (brs, 1H), 5.49 (s, 1H), 4.47 – 4.40 (m, 2H), 3.86 – 3.80 (m, 1H), 3.73 – 3.63 (m, 4H), 3.66 – 3.60 (m, 8H), 3.58 – 3.29 (m, 7H), 3.03 (s, 6H), 3.00 (s, 6H), 2.90 (dd, J = 15.3, 6.1 Hz, 1H), 2.52 – 2.36 (m, 4H), 1.85 – 1.73 (m, 6H).

**HRMS ESI-TOF** m/z calculated for $C_{64}H_{70}N_{11}O_{12}$ $[M+H]^+$ 1184.5200. Found 1184.5204.

**5-[2-[2-[2-(2-(4-(2-(3-(3-((1R,3R)-1-(benzo[d][1,3]dioxol-5-yl)-3-(morpholine-4-carbonyl)-1,3,4,9-tetrahydro-2H-pyrido[3,4-b]indol-2-yl)-3-oxopropyl)-3H-diazirin-3-yl)ethyl)-1H-1,2,3-triazol-1-yl)ethoxy)ethoxy]ethoxy]ethylcarbamoyl]-2-(3,6-Bis(dimethylamino)xanthylium-9-yl)benzoate (DO-35B) (32)**

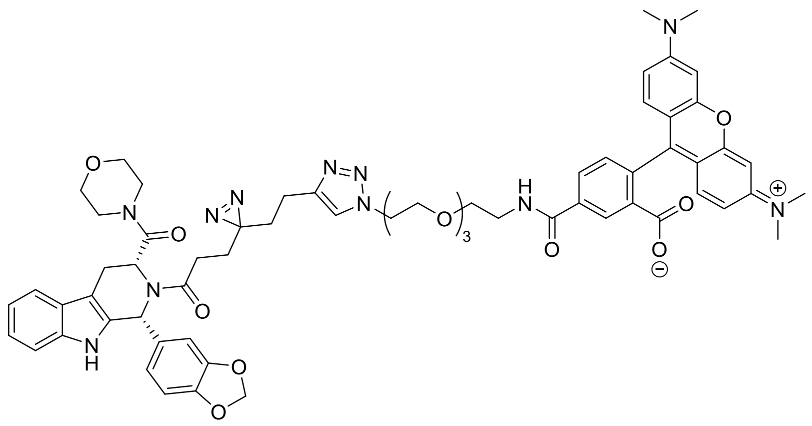

Prepared in an analogous fashion from **WX-02-38**. 61% yield as a red solid.

**$^1$H-NMR** (600 MHz, $CDCl_3$): δ ppm 9.02 (s, 1H), 8.46 (s, 1H), 8.20 (d, J = 8.0 Hz, 1H), 7.64 (brs, 1H), 7.56 (s, 1H), 7.51 (d, J = 7.6 Hz, 1H), 7.38 – 7.31 (m, 1H), 7.20 (d, J = 8.0 Hz, 1H), 7.18 – 7.12 (m, 1H), 7.11 – 7.04 (m, 2H), 6.79 (d, J = 7.1 Hz, 2H), 6.72 (d, J = 6.1 Hz, 1H), 6.67 (d, J = 8.3 Hz, 1H), 6.64 (d, J = 9.0 Hz, 1H), 6.53 (d, J = 2.5 Hz, 1H), 6.49 (d, J = 2.5 Hz, 1H), 6.46 (d, J = 9.2 Hz, 1H), 6.38 (d, J = 9.1 Hz, 1H), 6.27 (brs, 1H), 5.85 (s, 2H), 5.70 (brs, 1H), 5.50 (brs, 1H), 4.47 – 4.41 (m, 2H), 3.86 – 3.80 (m, 1H), 3.74 – 3.67 (m, 4H), 3.67 – 3.59 (m, 8H), 3.58 – 3.27 (m, 7H), 3.04 (s, 6H), 3.01 (s, 6H), 2.90 (dd, J = 15.3, 6.1 Hz, 1H), 2.54 – 2.35 (m, 4H), 1.85 – 1.70 (m, 6H).

**HRMS ESI-TOF** m/z calculated for $C_{64}H_{70}N_{11}O_{12}$ $[M+H]^+$ 1184.5200. Found 1184.5232.

**Synthesis of DO-39A and DO-39B**

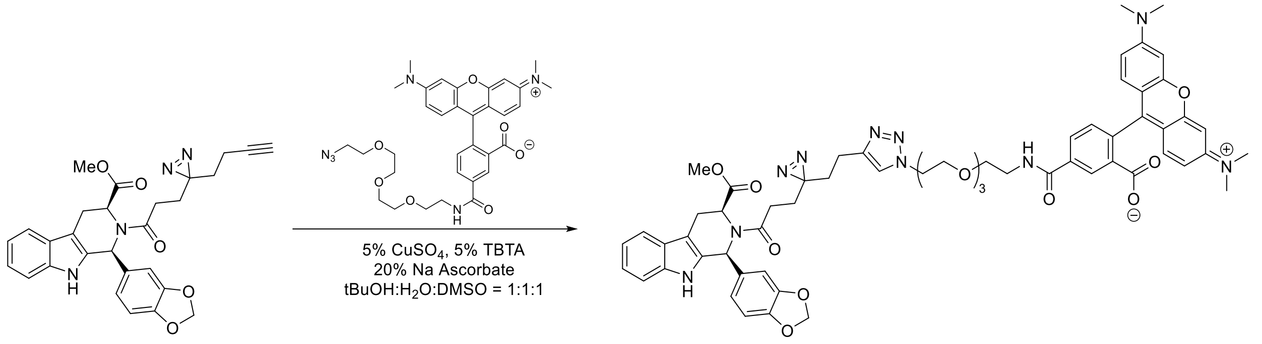


**DBK-032A (1)** **DO-39A (33)**

**5-[2-[2-[2-(2-(4-(2-(3-(3-((1S,3S)-1-(benzo[d][1,3]dioxol-5-yl)- 3-(methoxycarbonyl)-1,3,4,9-tetrahydro-2H-pyrido[3,4-b]indol-2-yl)-3-oxopropyl)-3H-diazirin-3-yl)ethyl)-1H-1,2,3-triazol-1-yl)ethoxy)ethoxy]ethoxy]ethylcarbamoyl]-2-(3,6-Bis(dimethylamino)xanthylium-9-yl)benzoate (DO-39A) (33)**

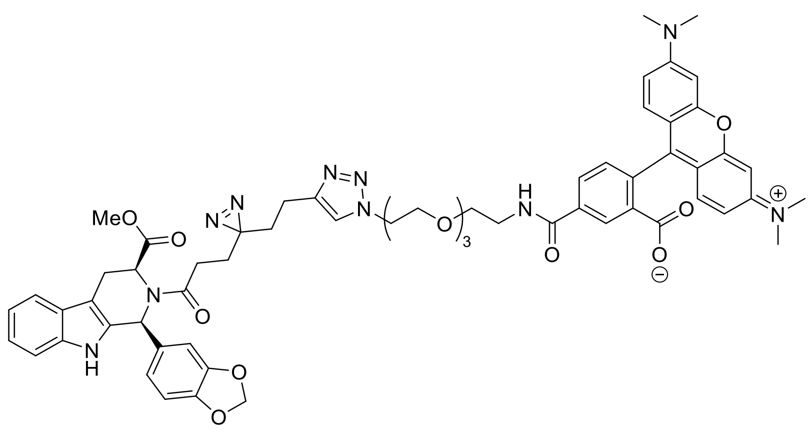

To a solution of **DBK-032A** (7.9 mg, 0.016 mmol) in H2O/t-BuOH/DMSO (0.2 mL/0.2 mL/0.1 mL) were added TAMRA-(PEG)$_3$-N$_3$ (10 mg, 0.016 mmol), TBTA (0.4 mg, 0.8 μmol), sodium ascorbate (0.6 mg, 3.2 μmol) and $CuSO_4$ (0.3 mg, 0.8 μmol). The mixture was stirred at 20 °C for 12 hours. Upon completion, the resulting mixture was purified by prep-TLC (DCM/MeOH = 5/1) to obtain **DO-39A** (14 mg, 80% yield) as a red solid.

**$^1$H-NMR** (600 MHz, $CDCl_3$): δ ppm 8.40 (s, 1H), 8.21 (d, J = 7.8 Hz, 2H), 7.58 – 7.41 (m, 3H), 7.27 (s, 1H), 7.22 (d, J = 8.0 Hz, 1H), 7.16 (t, J = 7.5 Hz, 1H), 7.12 (t, J = 7.4 Hz, 1H), 7.08 – 7.03 (m, 1H), 7.00 – 6.93 (m, 1H), 6.85 (s, 1H), 6.78 – 6.63 (m, 1H), 6.60 – 6.45 (m, 6H), 6.42 – 6.25 (m, 2H), 5.85 (s, 2H), 4.83 (d, J = 7.0 Hz, 1H), 4.48 – 4.37 (m, 2H), 3.87 – 3.79 (m, 2H), 3.74 – 3.68 (m, 4H), 3.67 – 3.59 (m, 9H), 3.48 (s, 1H), 3.13 (s, 3H), 3.03 – 2.97 (m, 12H), 2.50 (t, J = 8.0 Hz, 2H), 2.33 – 2.20 (m, 2H), 1.90 – 1.82 (m, 2H).

**HRMS ESI-TOF** m/z calculated for $C_{61}H_{65}N_{10}O_{12}$ $[M+H]^+$ 1129.4778. Found 1129.4799.

**5-[2-[2-[2-(2-(4-(2-(3-(3-((1R,3R)-1-(benzo[d][1,3]dioxol-5-yl)- 3-(methoxycarbonyl)-1,3,4,9-tetrahydro-2H-pyrido[3,4-b]indol-2-yl)-3-oxopropyl)-3H-diazirin-3-yl)ethyl)-1H-1,2,3-triazol-1-yl)ethoxy)ethoxy]ethoxy]ethylcarbamoyl]-2-(3,6-Bis(dimethylamino)xanthylium-9-yl)benzoate (DO-39B) (34)**

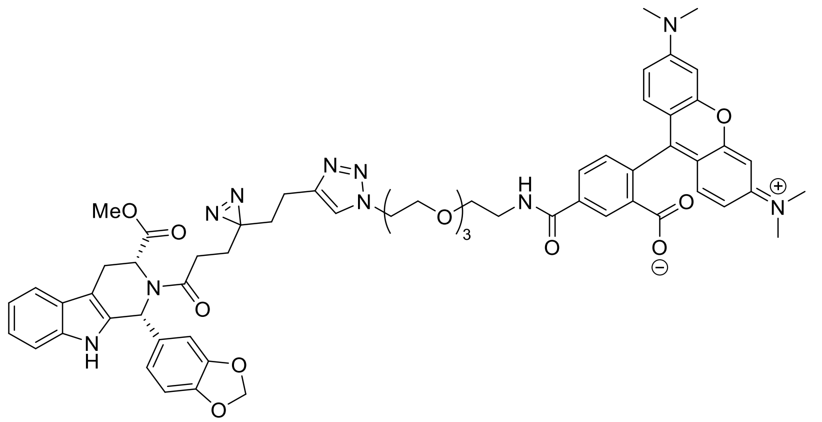

Prepared in an analogous fashion from **DBK-032B**. 81% yield as a red solid.

**$^1$H-NMR** (600 MHz, $CDCl_3$): δ ppm 8.39 (s, 1H), 8.29 – 8.13 (m, 2H), 7.62 – 7.41 (m, 3H), 7.27 (s, 1H, overlapped with solvent peak), 7.22 (d, J = 8.0 Hz, 1H), 7.16 (t, J = 7.5 Hz, 1H), 7.12 (t, J = 7.4 Hz, 1H), 7.09 – 7.03 (m, 1H), 6.95 (s, 1H), 6.85 (s, 1H), 6.74 – 6.61 (m, 1H), 6.60 – 6.45 (m, 6H), 6.42 – 6.22 (m, 2H), 5.85 (s, 2H), 4.83 (d, J = 7.0 Hz, 1H), 4.49 – 4.37 (m, 2H), 3.87 – 3.79 (m, 2H), 3.75 – 3.67 (m, 4H), 3.67 – 3.59 (m, 9H), 3.47 (s, 1H), 3.13 (s, 3H), 3.03 – 2.97 (m, 12H), 2.51 (t, J = 8.0 Hz, 2H), 2.31 – 2.21 (m, 2H), 1.90 – 1.81 (m, 2H).

**HRMS ESI-TOF** m/z calculated for $C_{61}H_{65}N_{10}O_{12}$ $[M+H]^+$ 1129.4778. Found 1129.4805.

### Spectroscopic data

#### $^1$H NMR spectrum of DBK-032A in $CD_3OD$

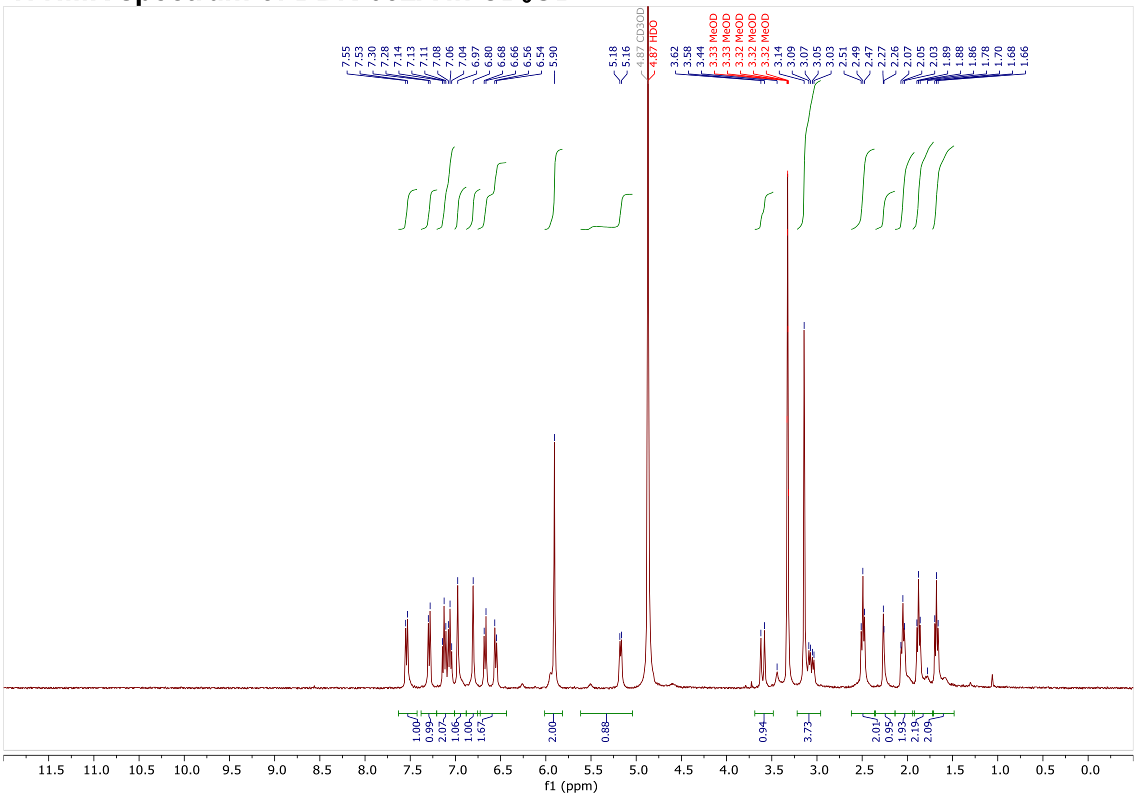

#### $^1$H NMR spectrum of DBK-032B in $CD_3OD$

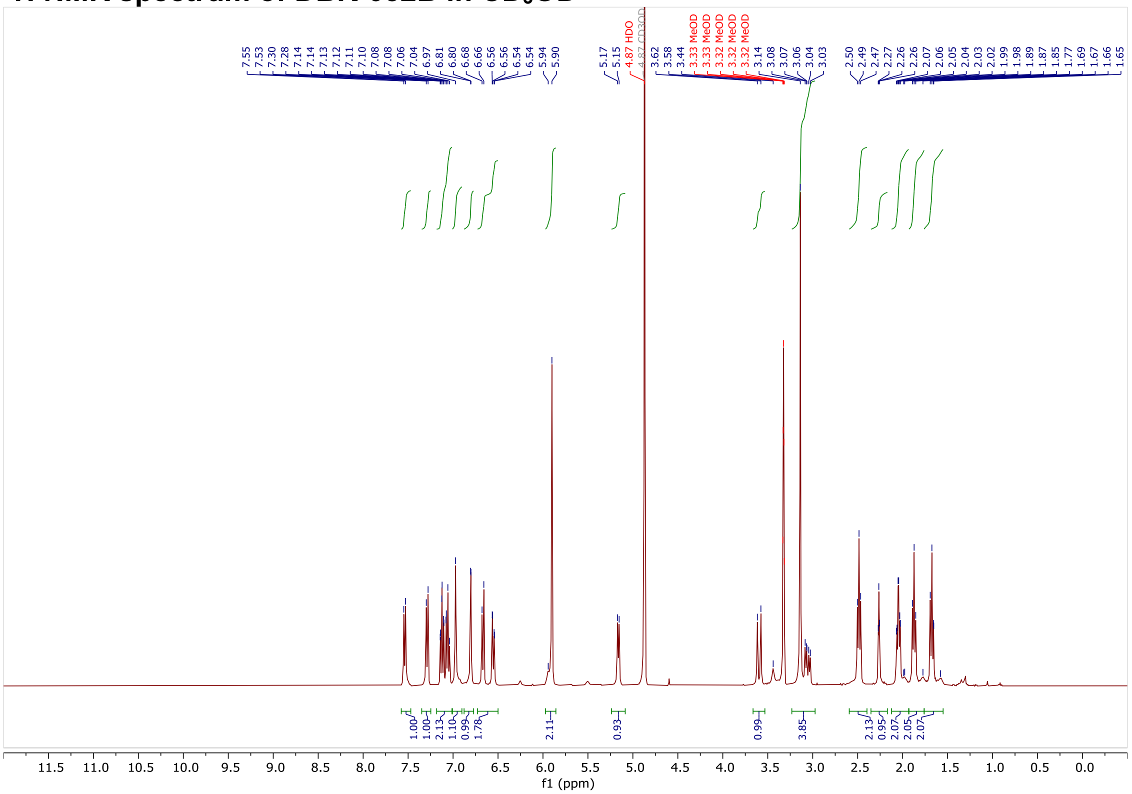

#### $^{1}H$ NMR spectrum of DBK-023A in $CD_3OD$

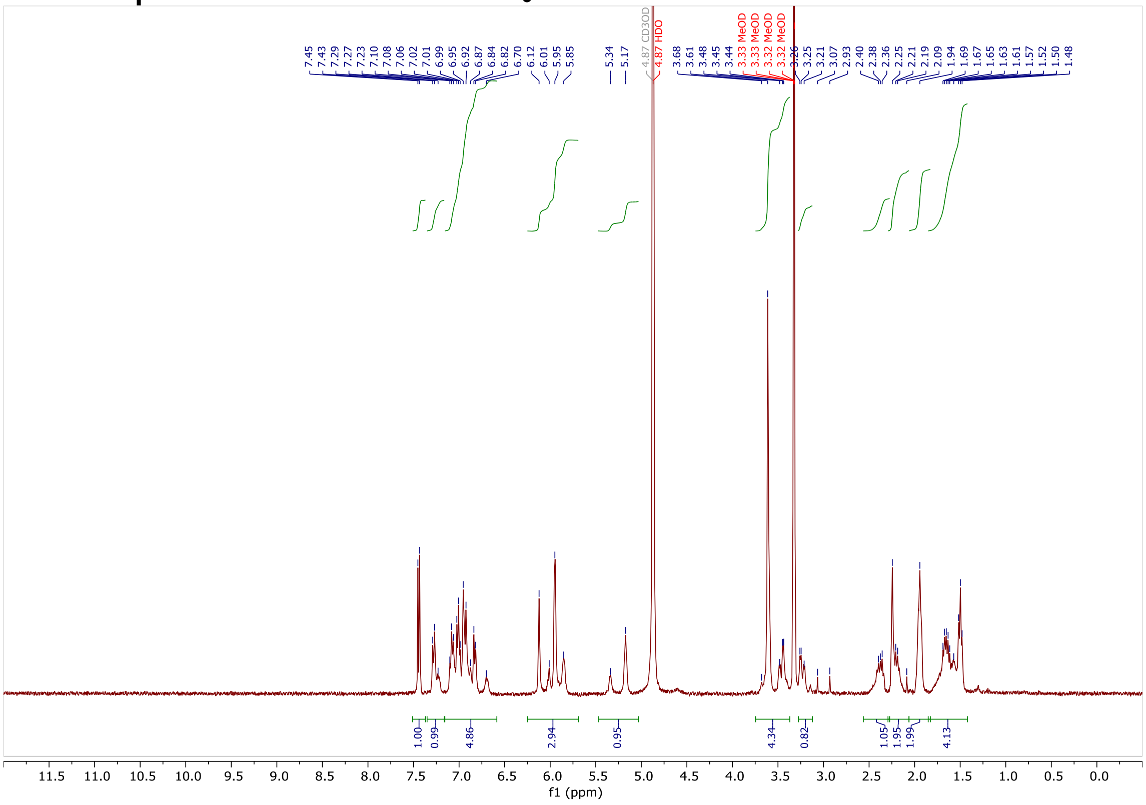

#### $^{1}H$ NMR spectrum of DBK-023B in $CD_3OD$

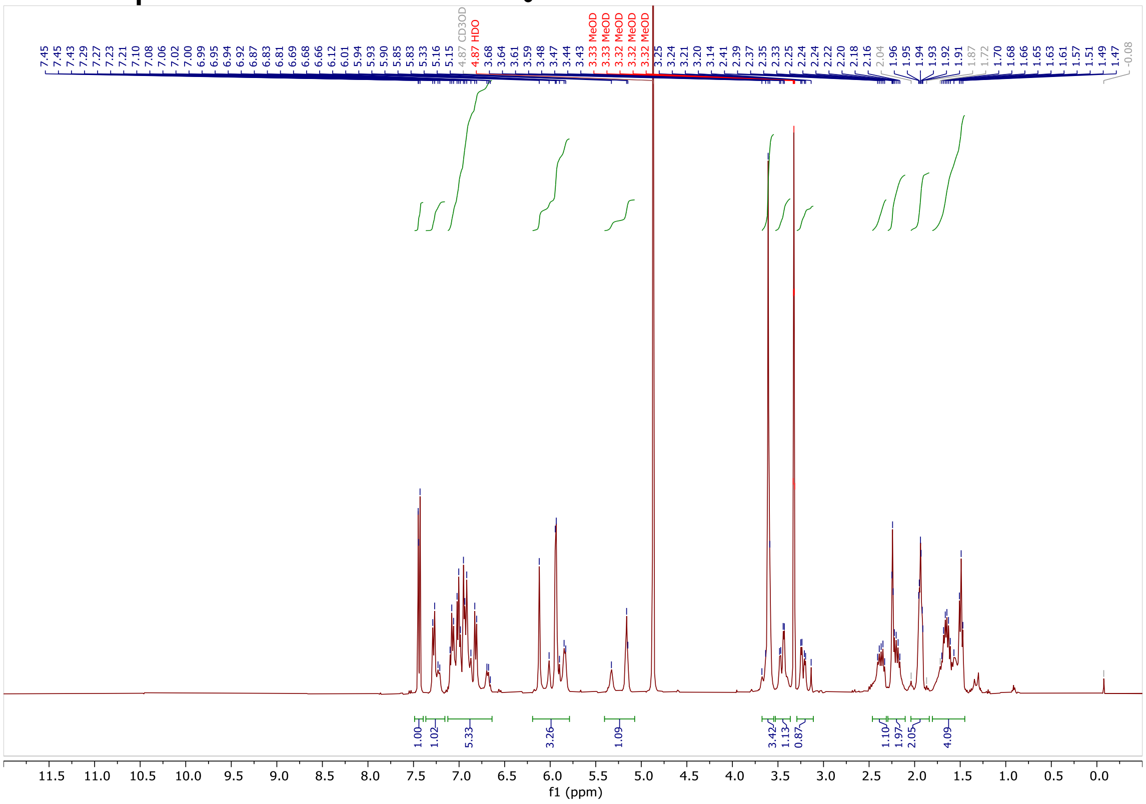

#### $^1$H NMR spectrum of DBK-070A in $CDCl_3$

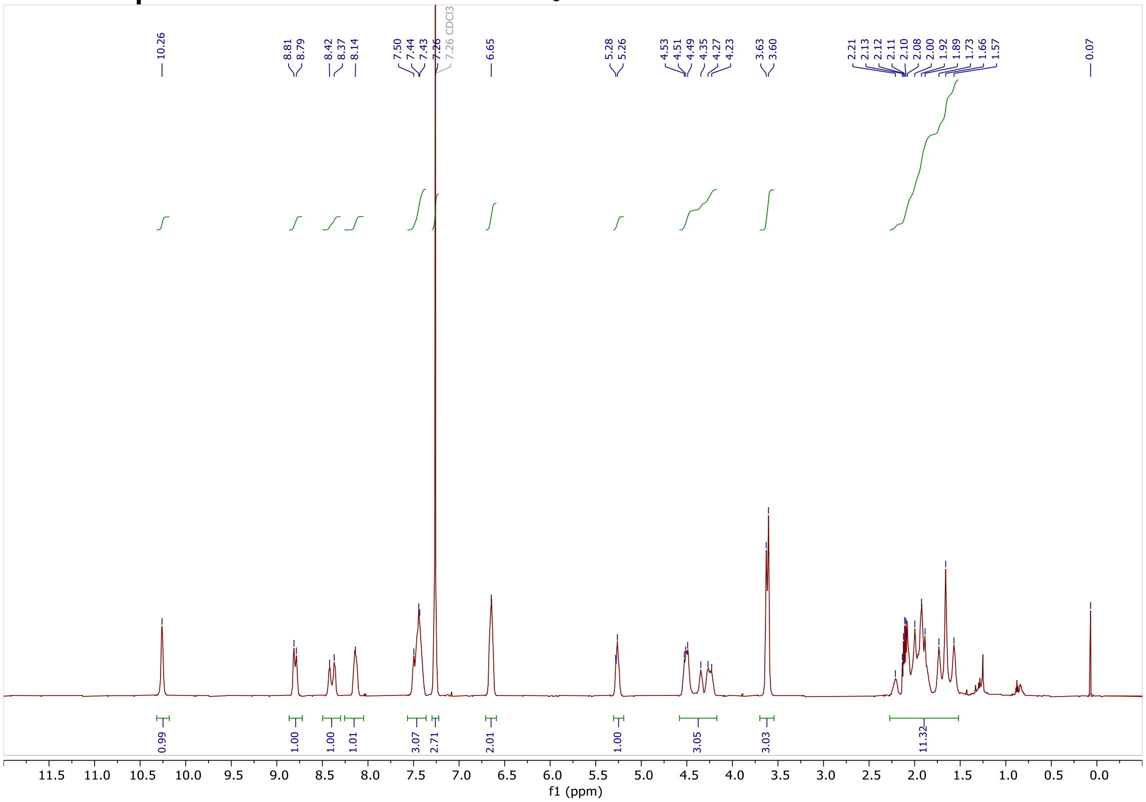

#### $^1$H NMR spectrum of DBK-070B in $CDCl_3$

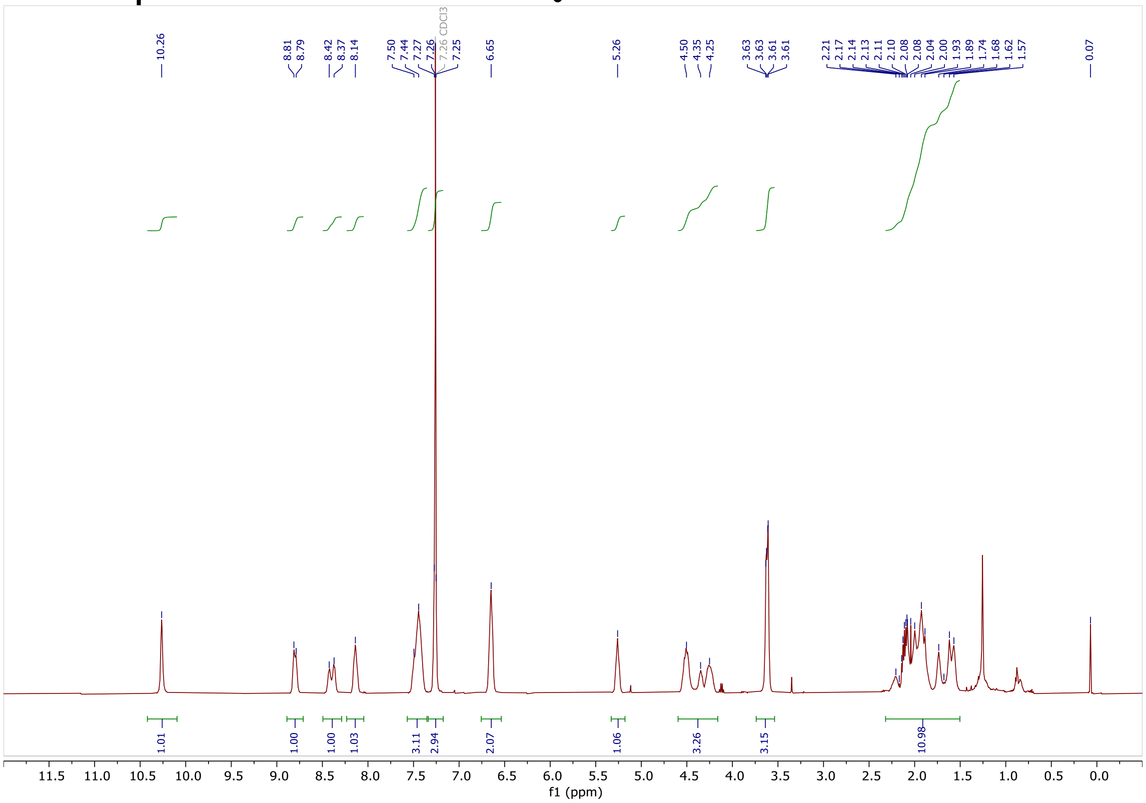

#### $^{1}H$ NMR spectrum of DBK-071A in $CDCl_3$

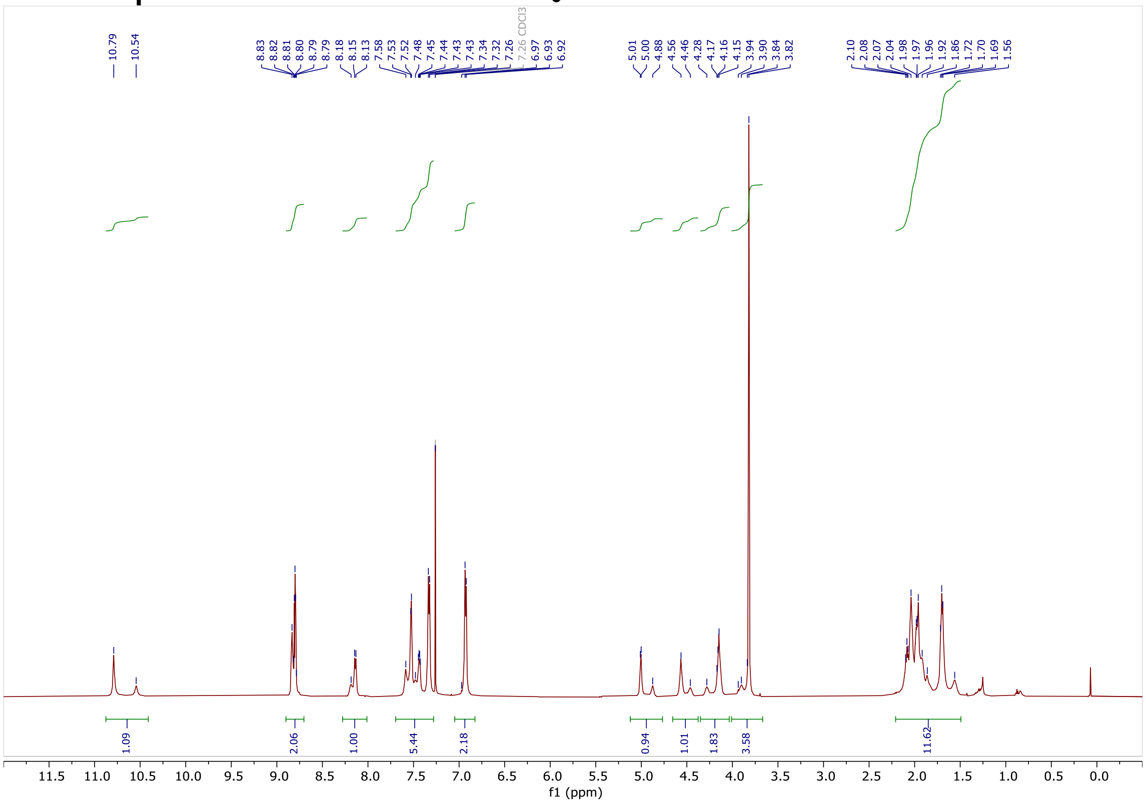

#### $^{1}H$ NMR spectrum of DBK-071B in $CDCl_3$

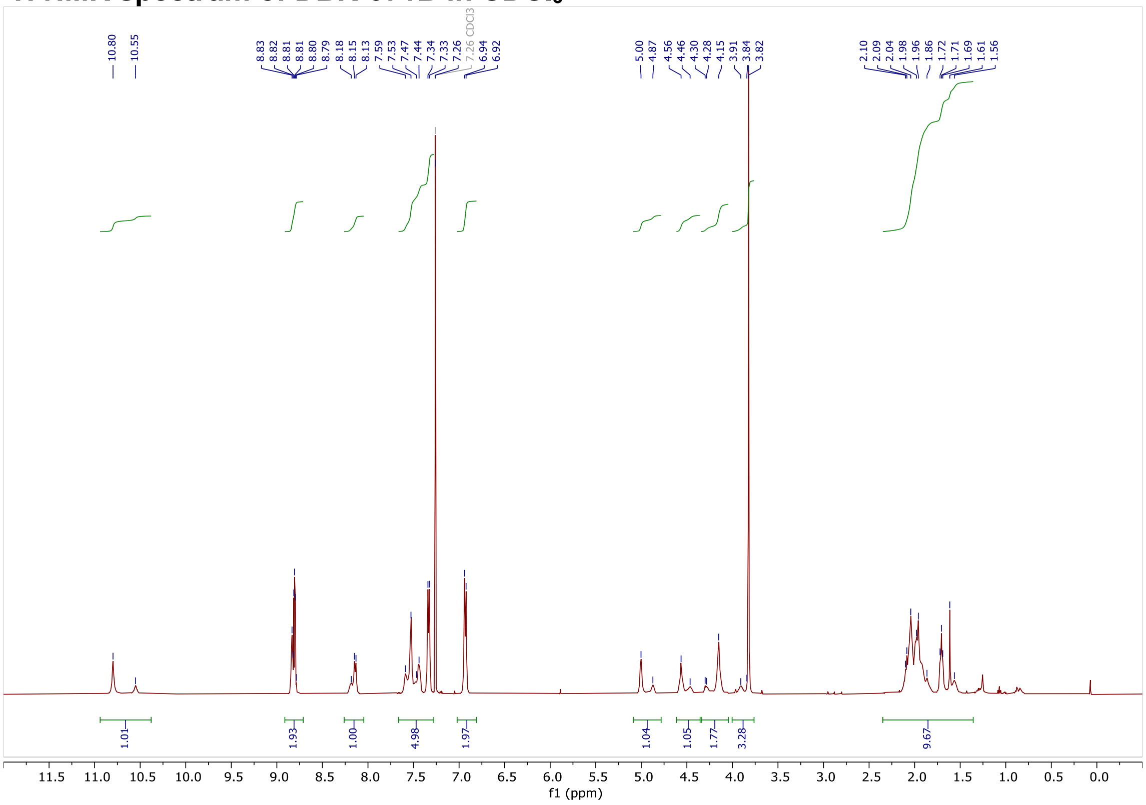

#### $^1$H NMR spectrum of DO-7A in $CDCl_3$

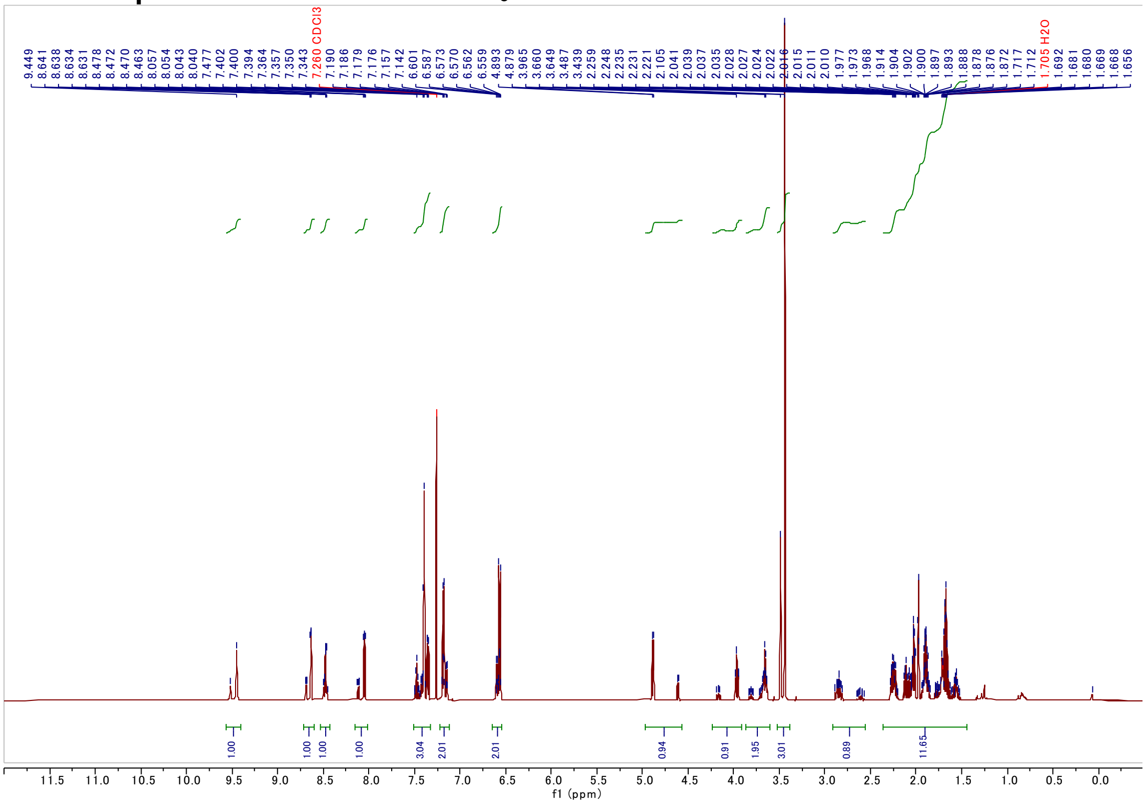

#### $^1$H NMR spectrum of DO-7B in $CDCl_3$

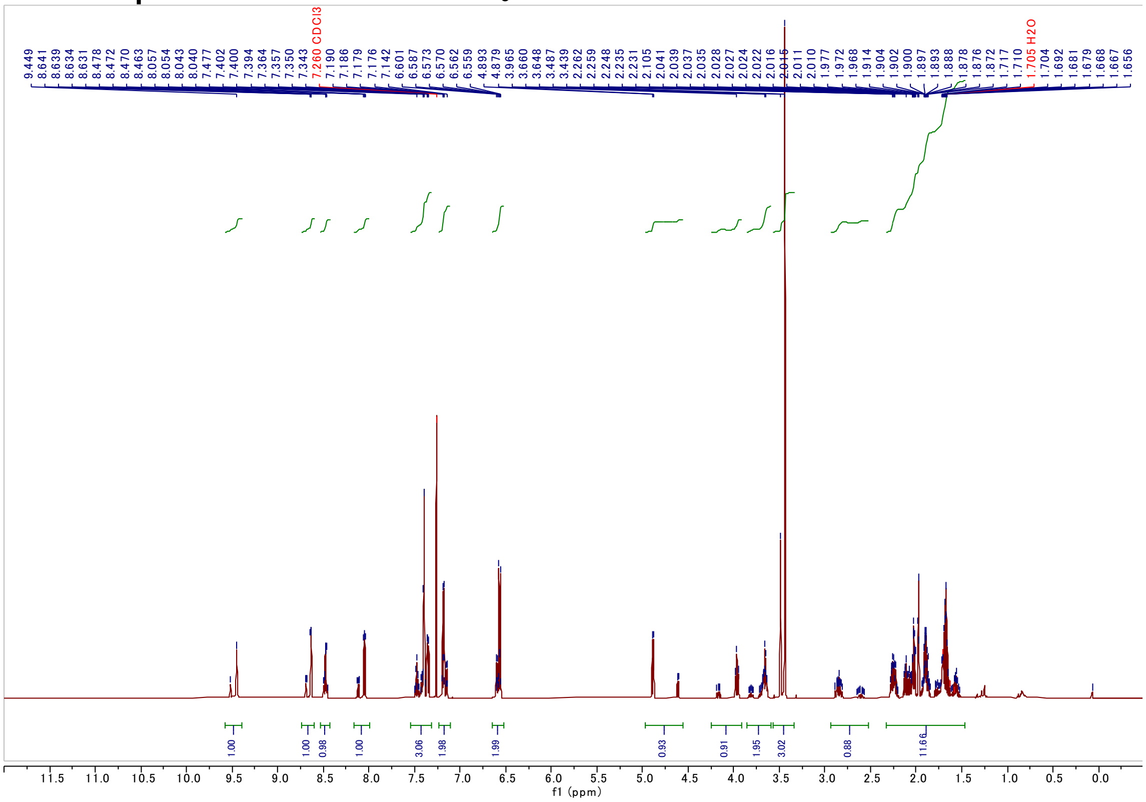

#### $^{1}H$ NMR spectrum of DO-8A in $CDCl_3$

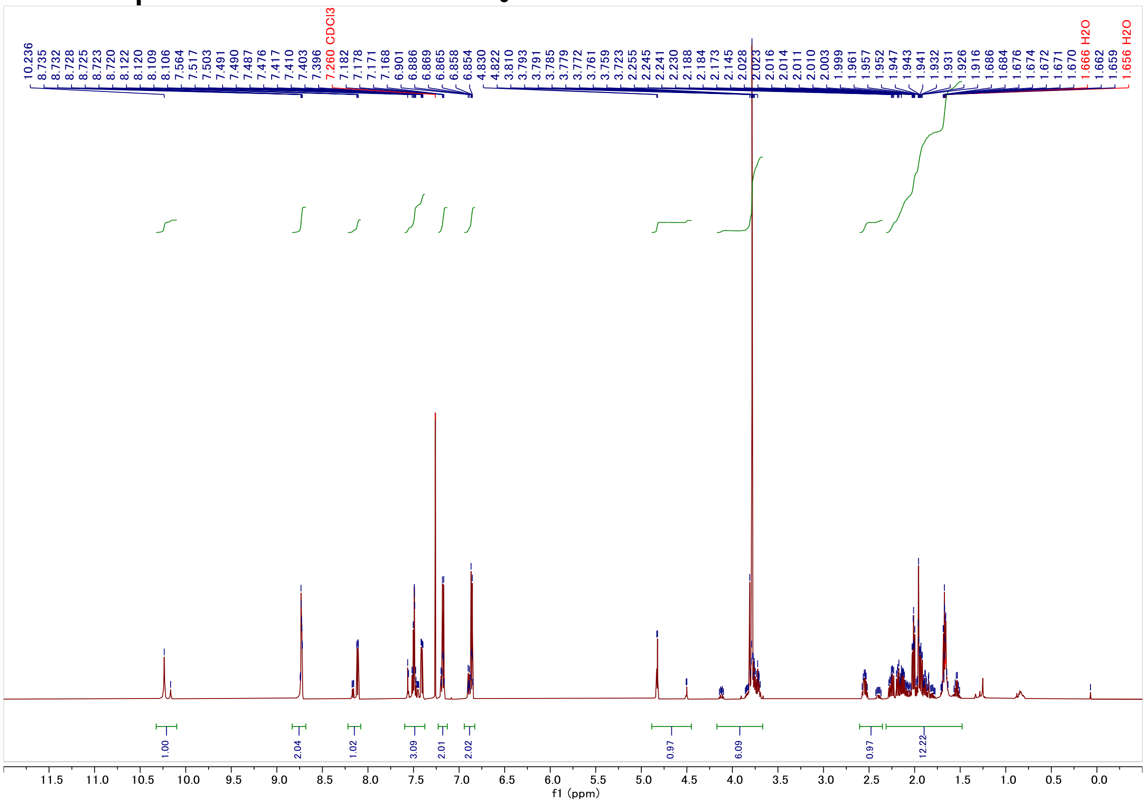

#### $^{1}H$ NMR spectrum of DO-8B in $CDCl_3$

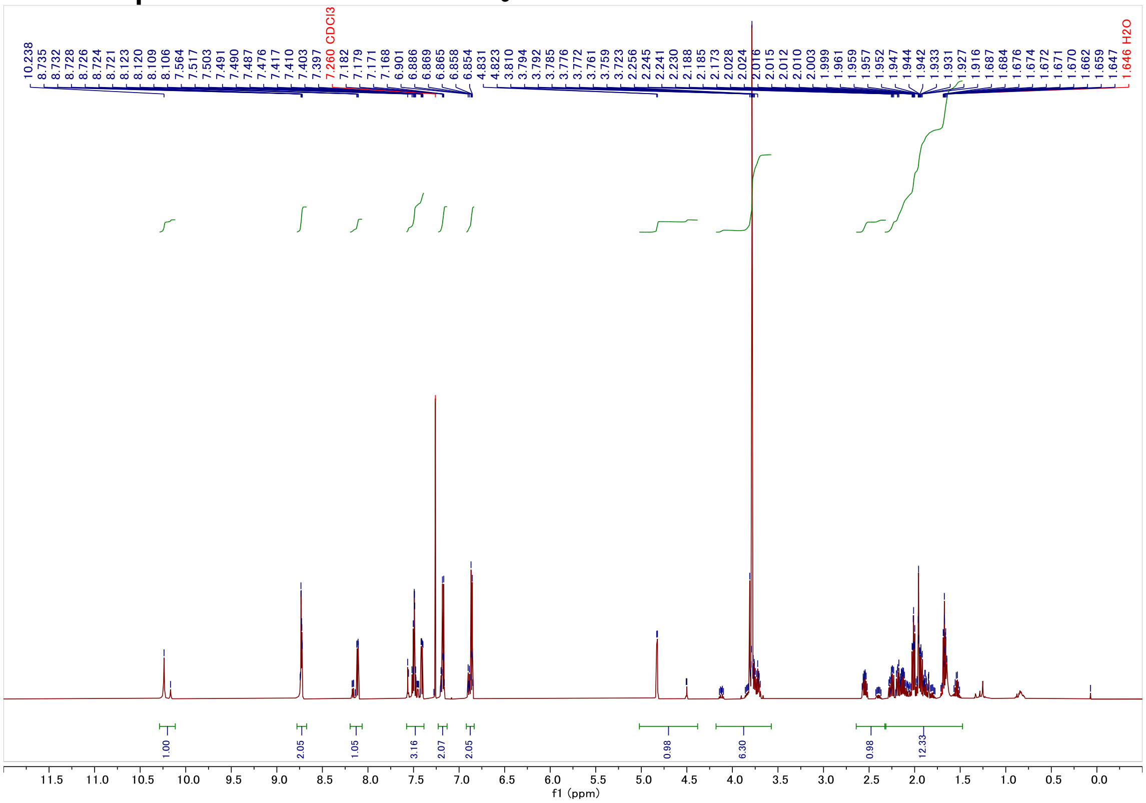

#### $^{1}H$ NMR spectrum of WX-02-18 in $CD_3OD$

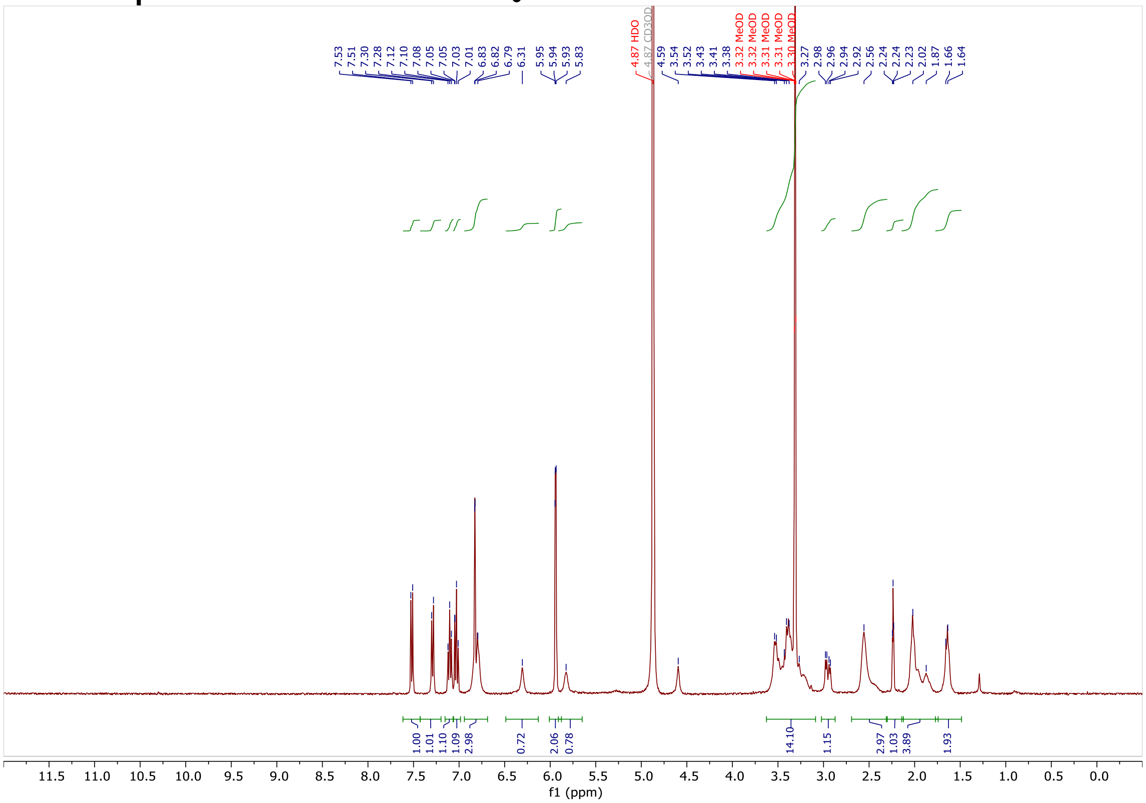

#### $^{1}H$ NMR spectrum of WX-02-38 in $CD_3OD$

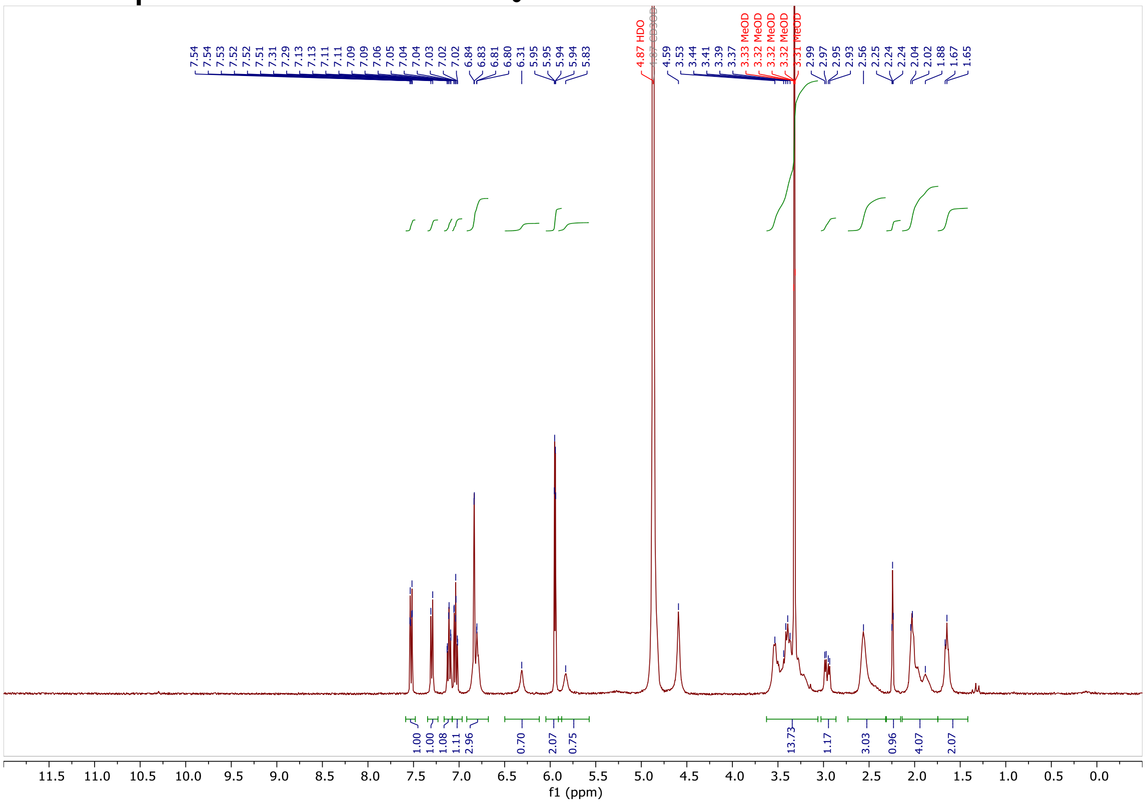

#### $^1$H NMR spectrum of WX-02-28 in $CD_3OD$

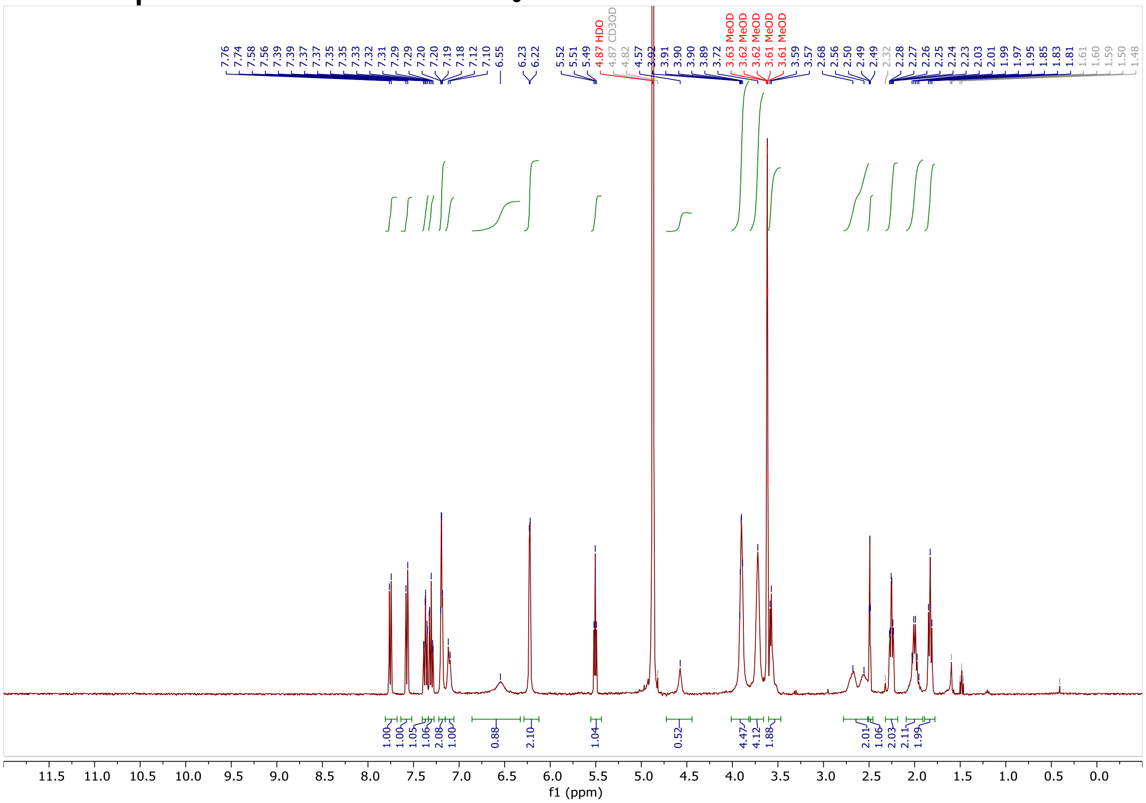

#### $^1$H NMR spectrum of WX-02-48 in $CD_3OD$

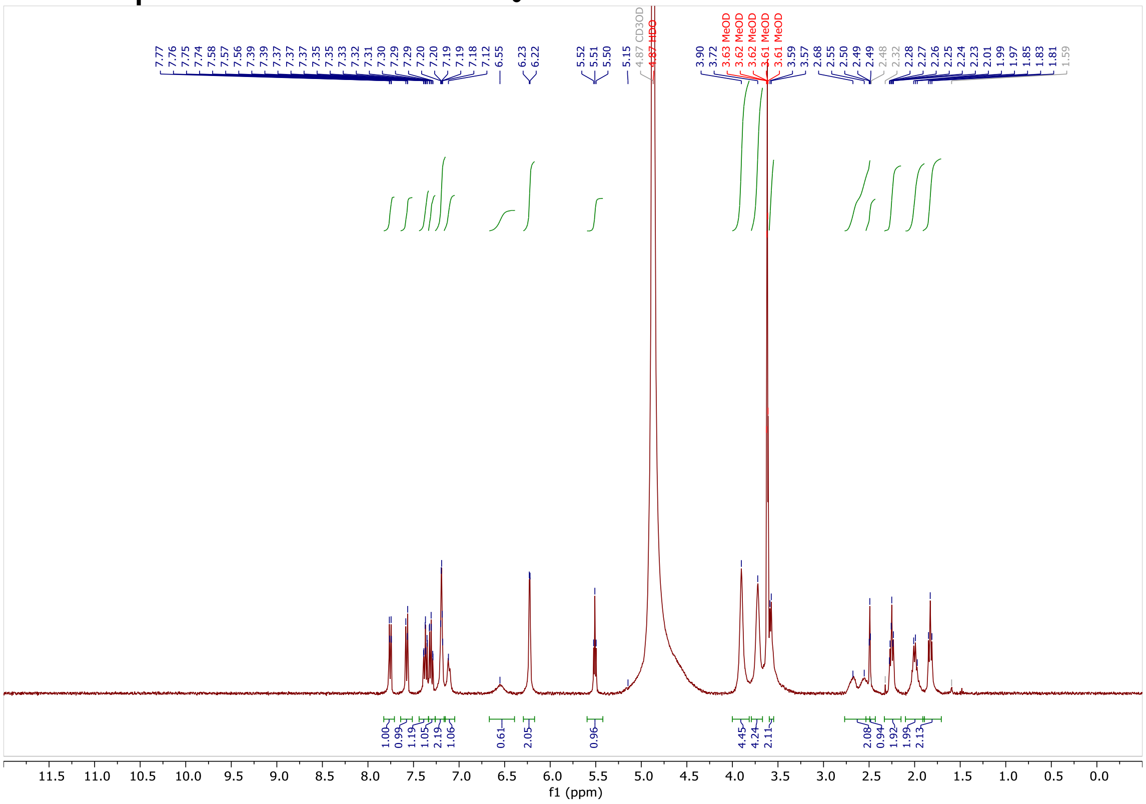

#### $^1$H NMR spectrum of WX-02-22 in $CD_3OD$

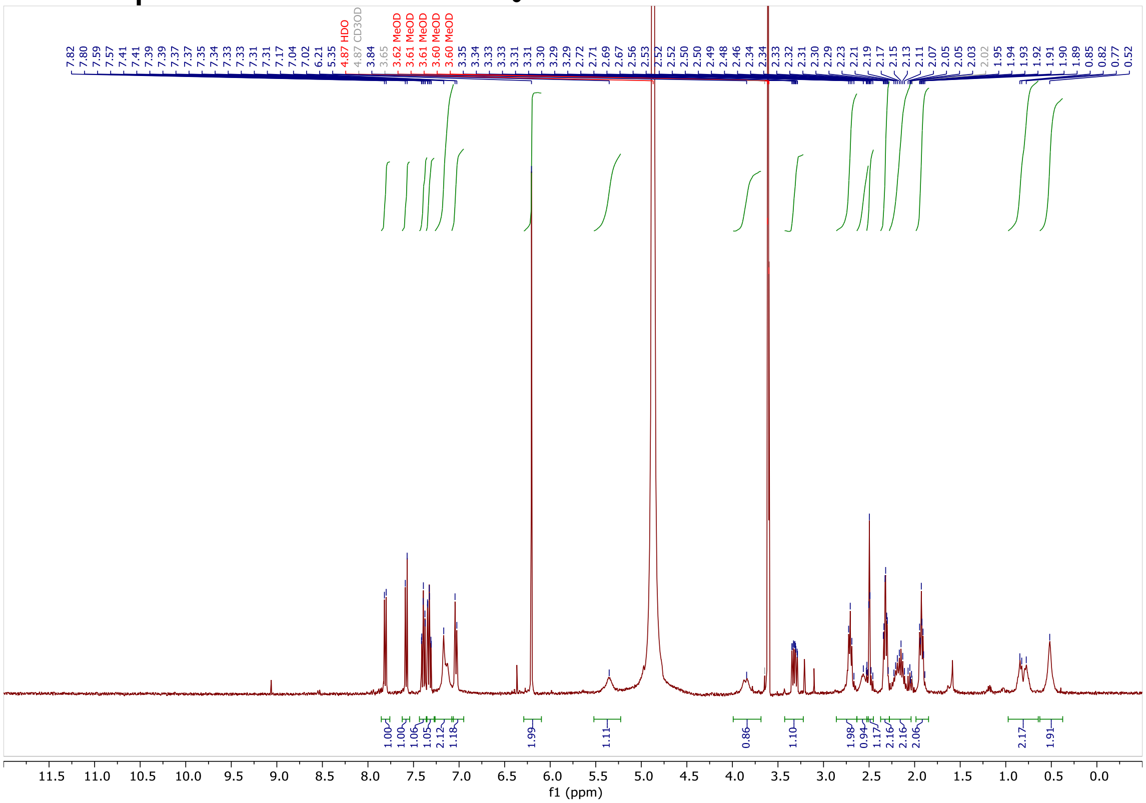

#### $^1$H NMR spectrum of WX-02-42 in $CD_3OD$

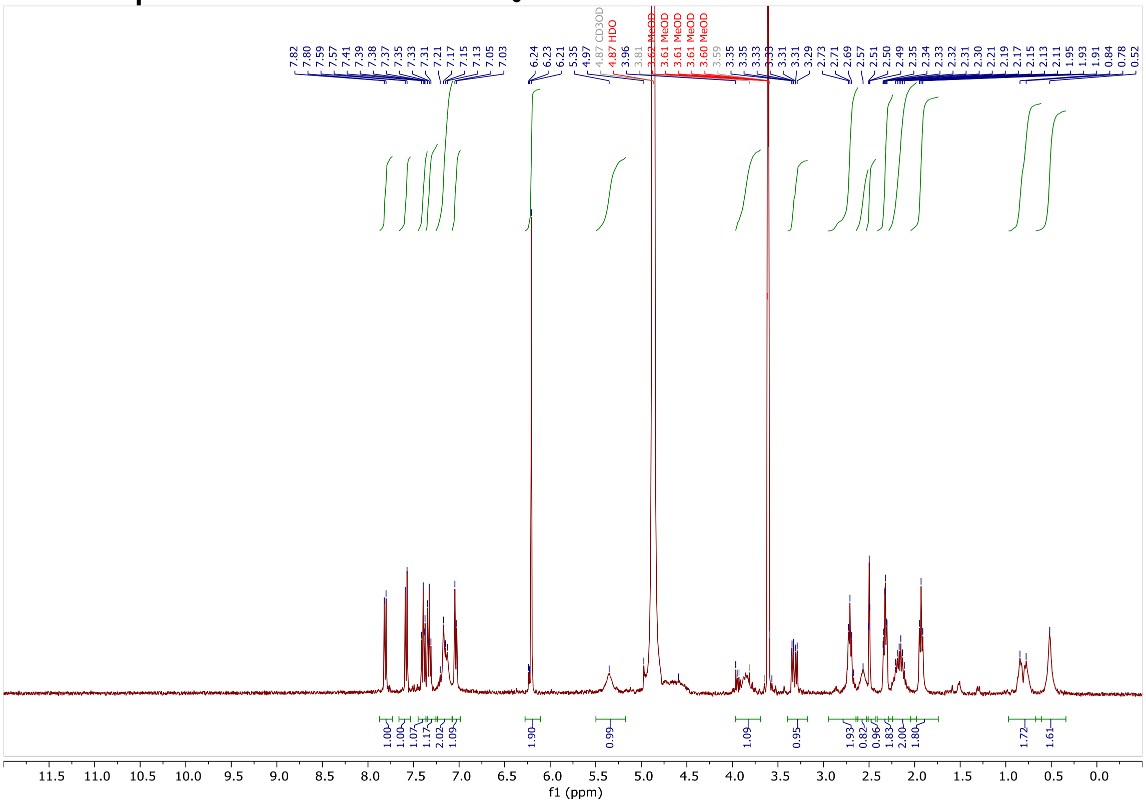

### $^1$H NMR spectrum of WX-02-20 in $CD_3OD$

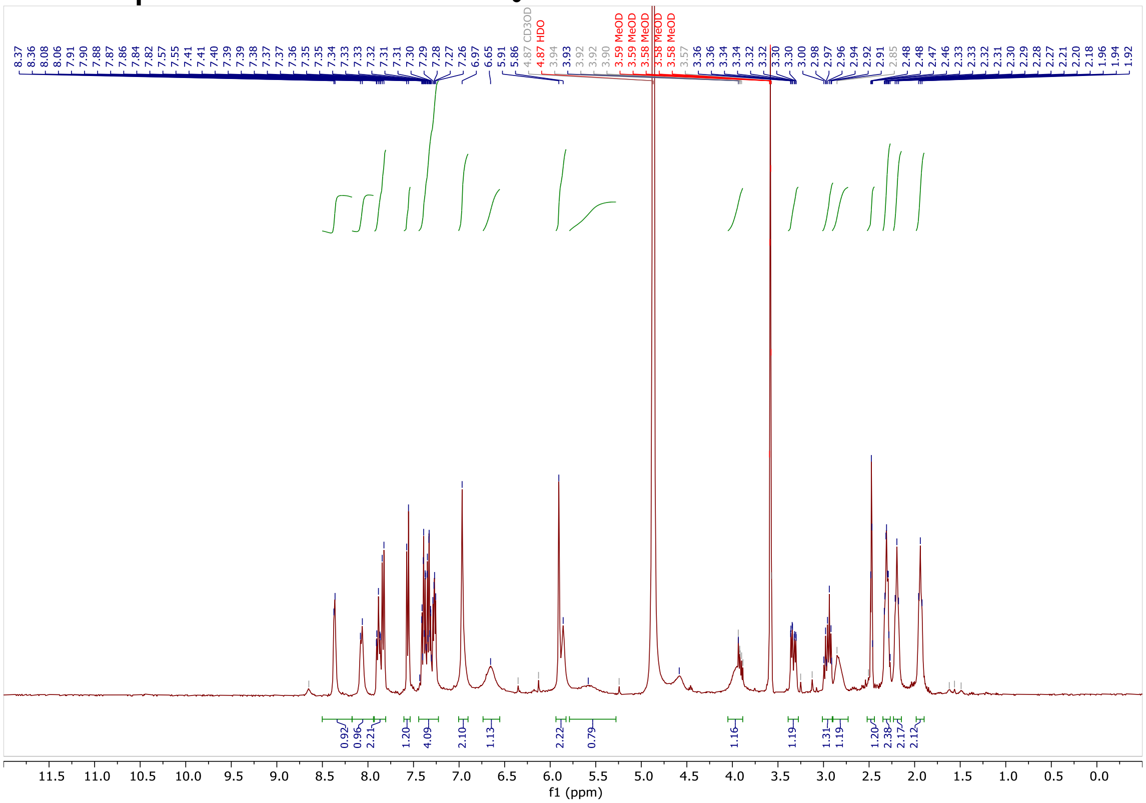

### $^1$H NMR spectrum of WX-02-21 in $CD_3OD$

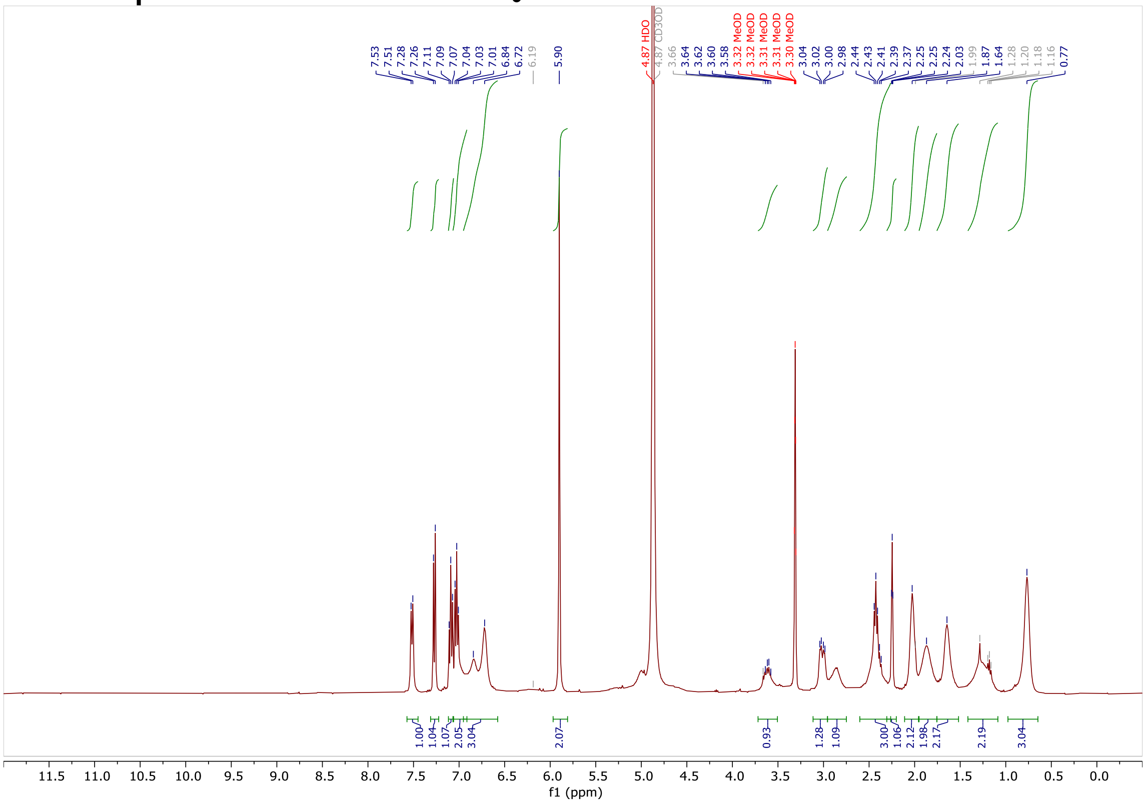

#### $^1H$ NMR spectrum of WX-02-19 in $CD_3OD$

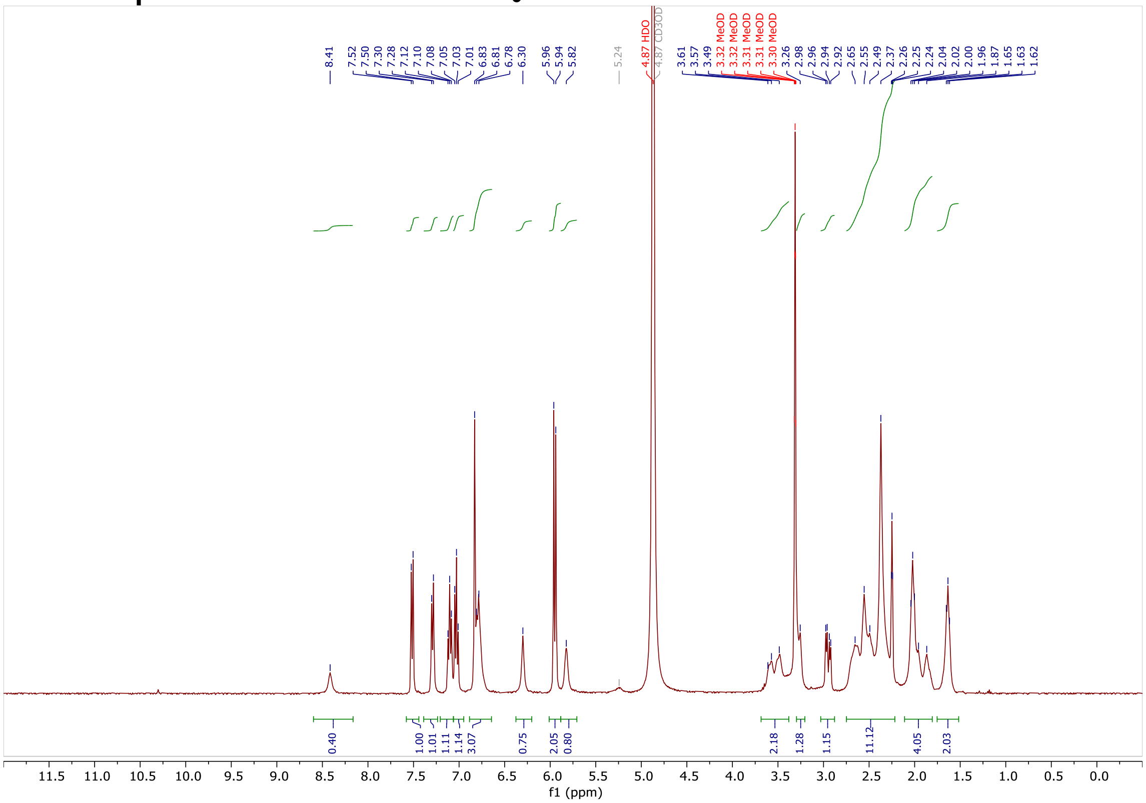

#### $^1H$ NMR spectrum of WX-02-221 in $CD_3OD$

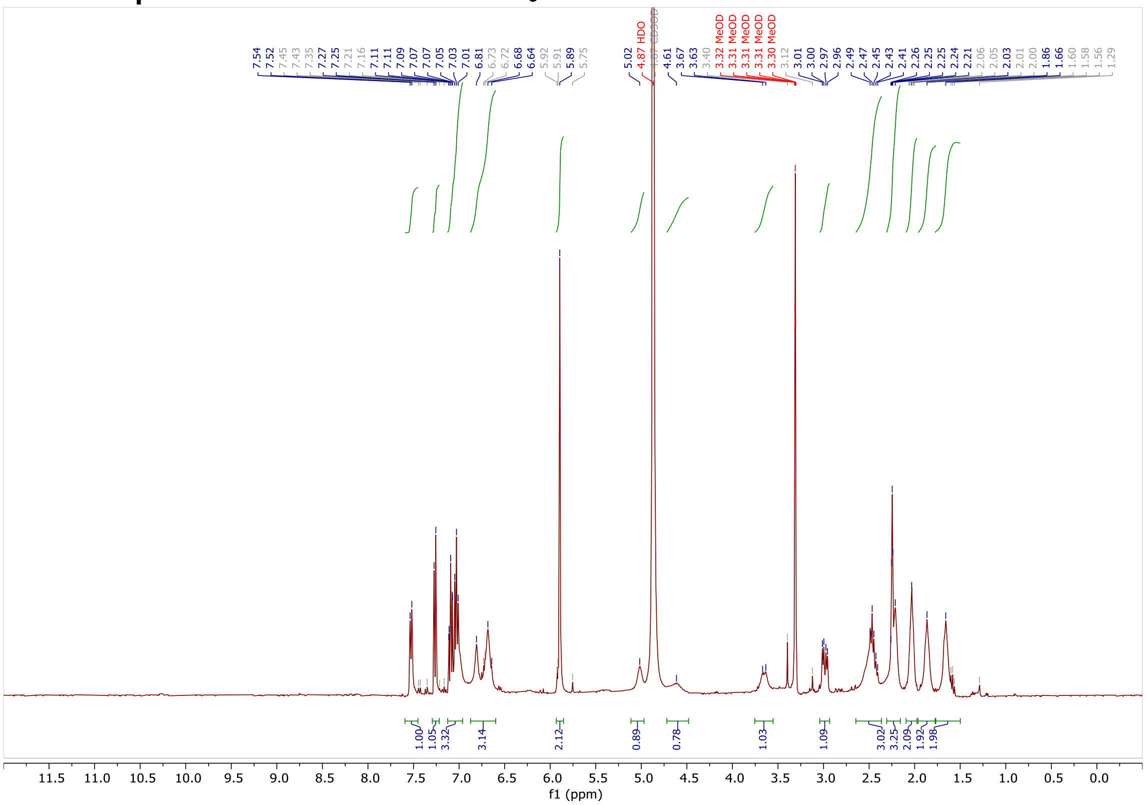

#### $^{1}$H NMR spectrum of WX-03-93 in $CD_3OD$

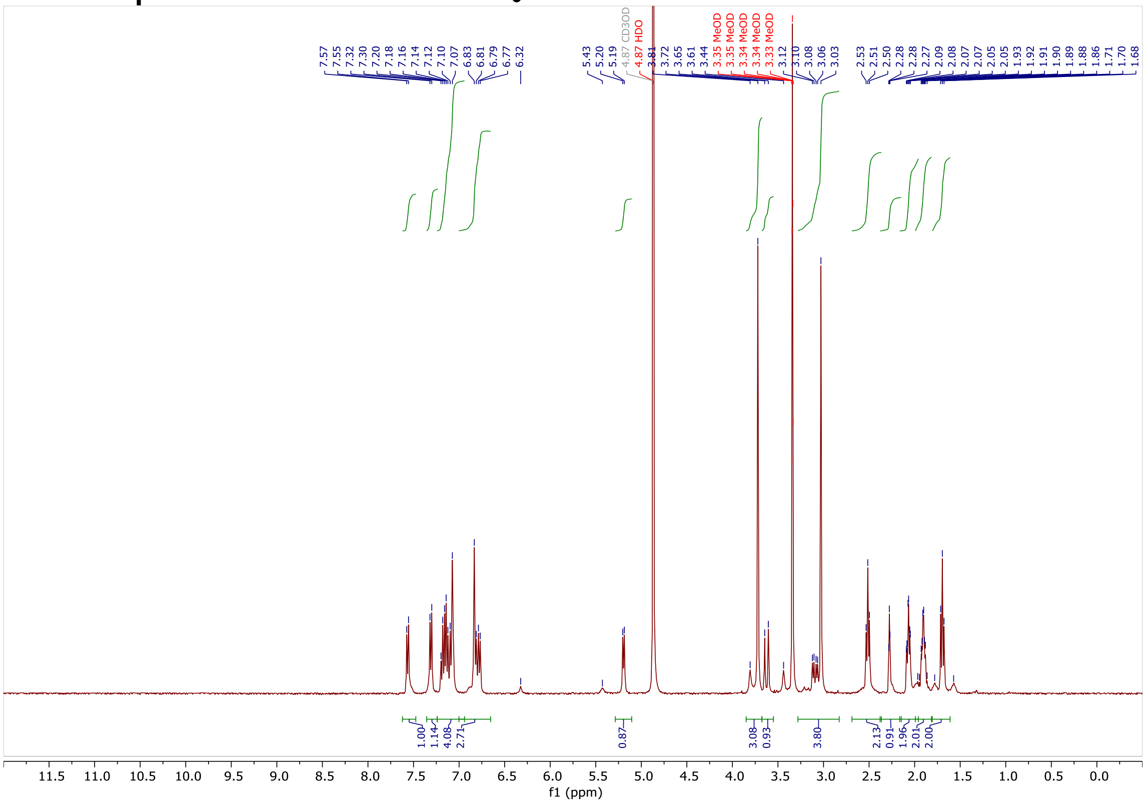

#### $^{1}$H NMR spectrum of WX-03-95 in $CD_3OD$

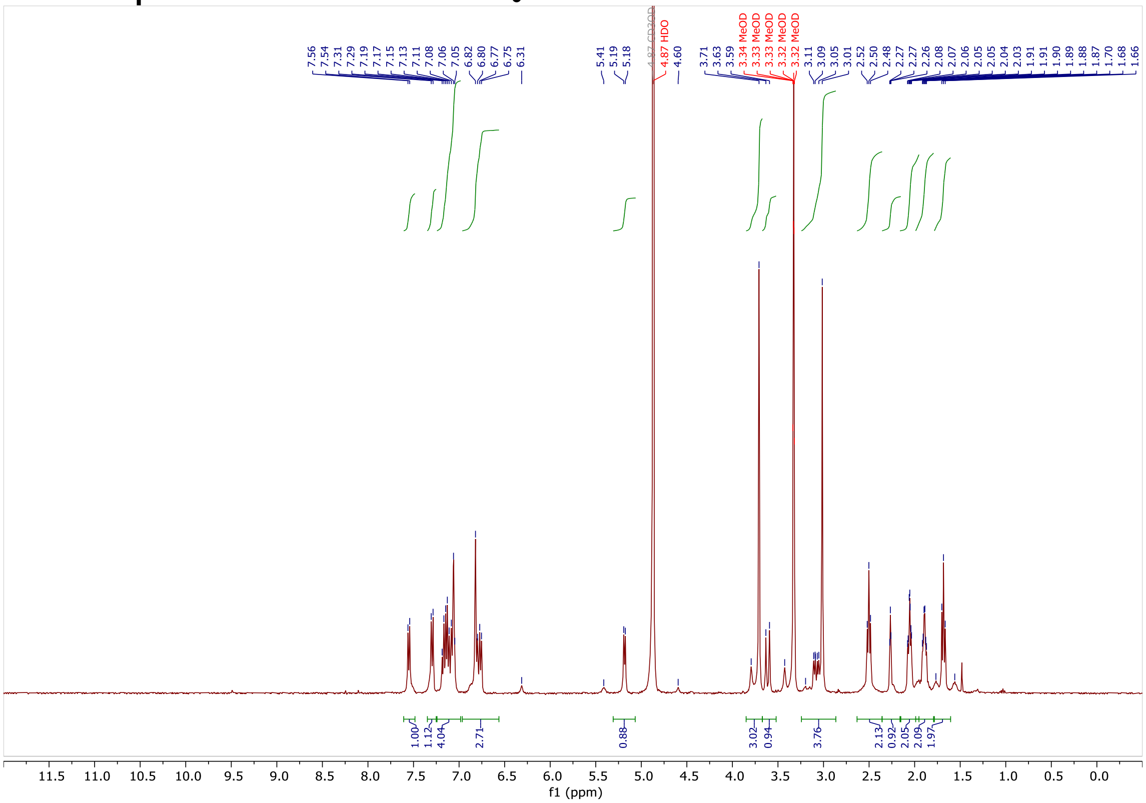

#### $^1$H NMR spectrum of WX-03-97 in $CD_3OD$

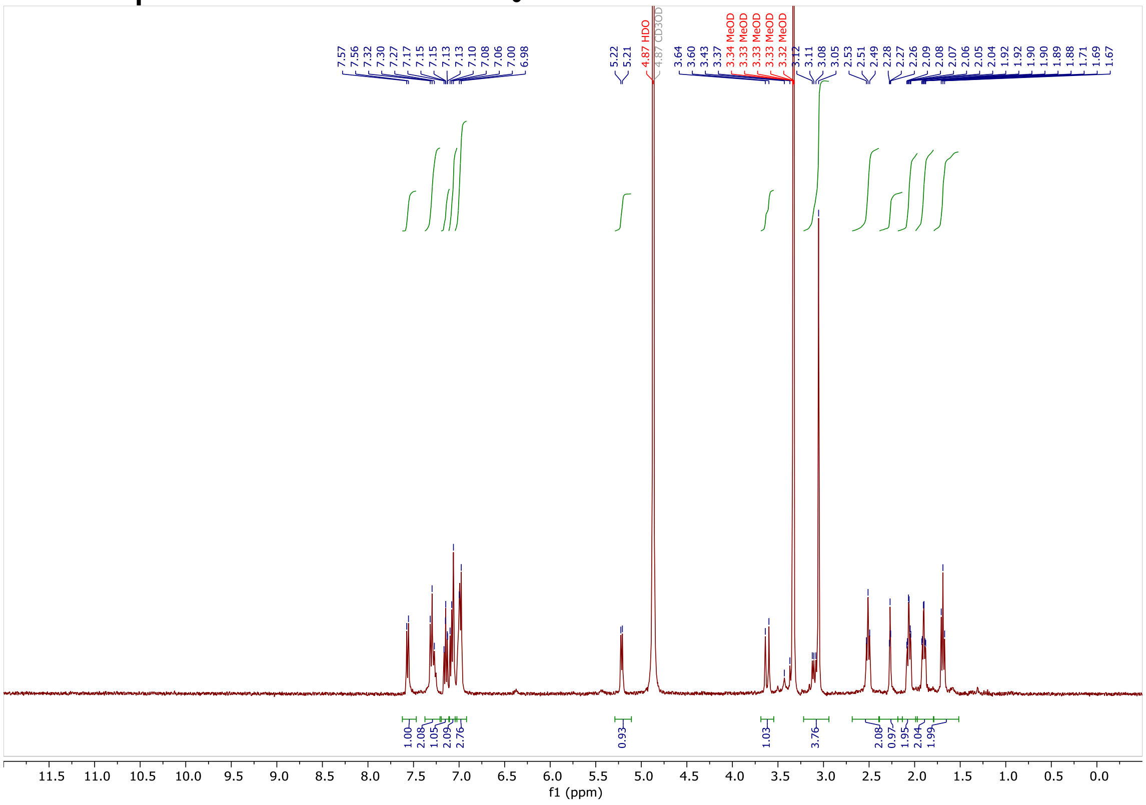

#### $^1$H NMR spectrum of WX-03-99 in $CD_3OD$

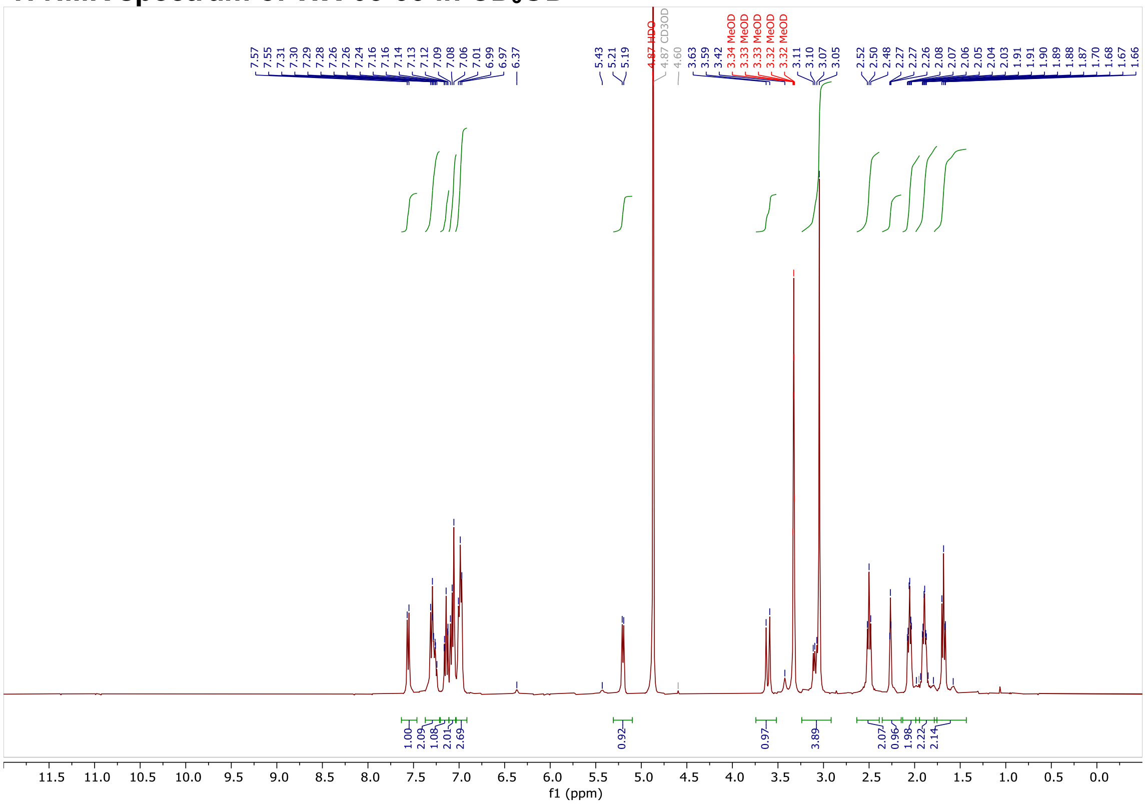

#### $^{1}$H NMR spectrum of DBK-073A in $CDCl_3$

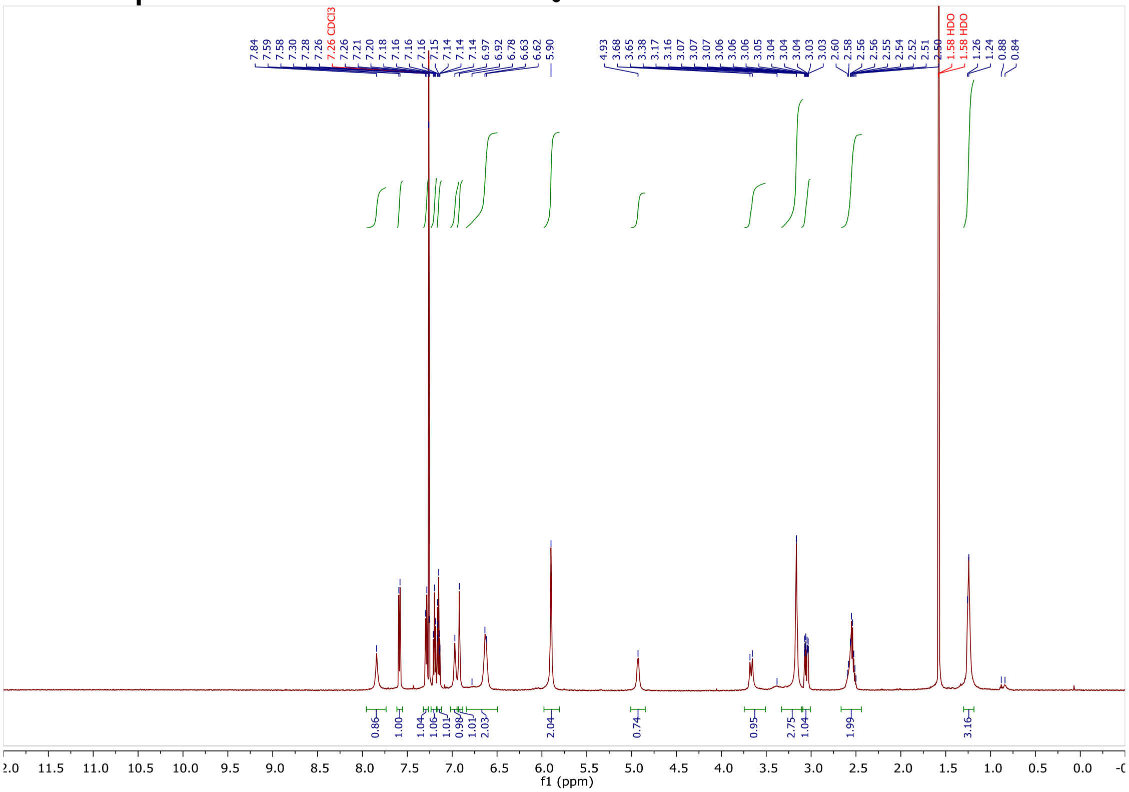

#### $^{1}$H NMR spectrum of DBK-073B in $CDCl_3$

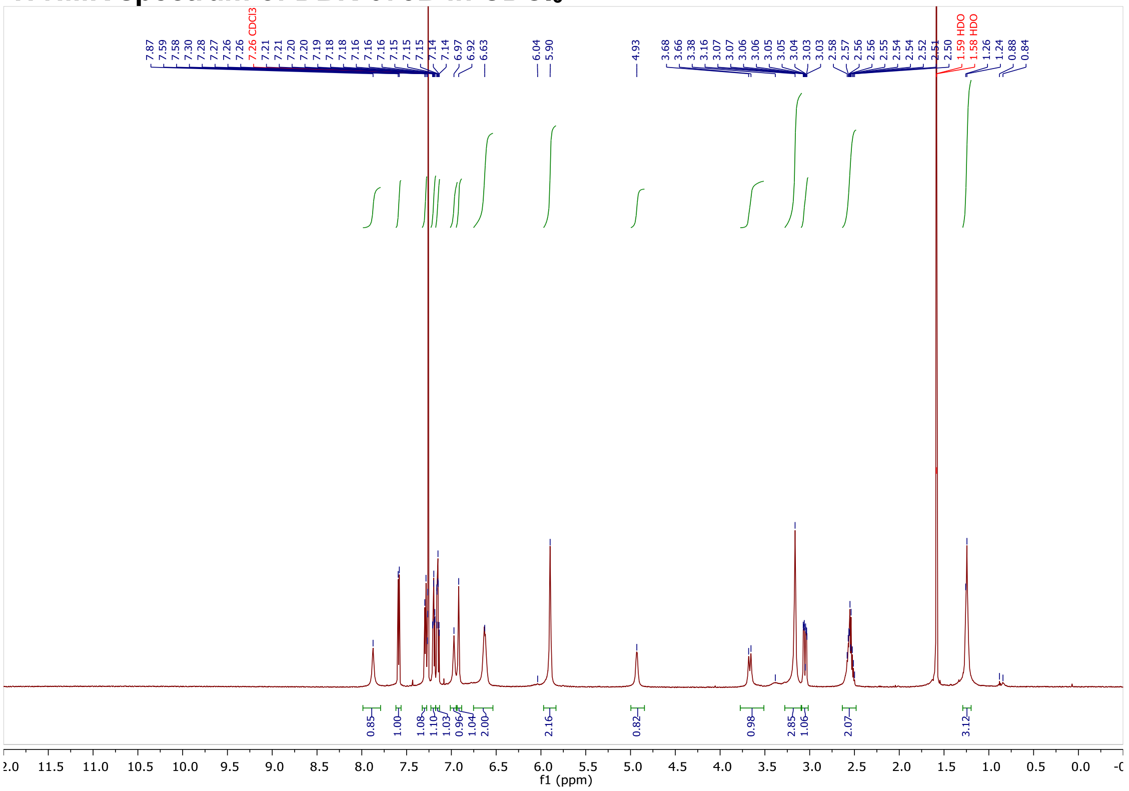

#### $^1H$ NMR spectrum of DO-40A in $CDCl_3$

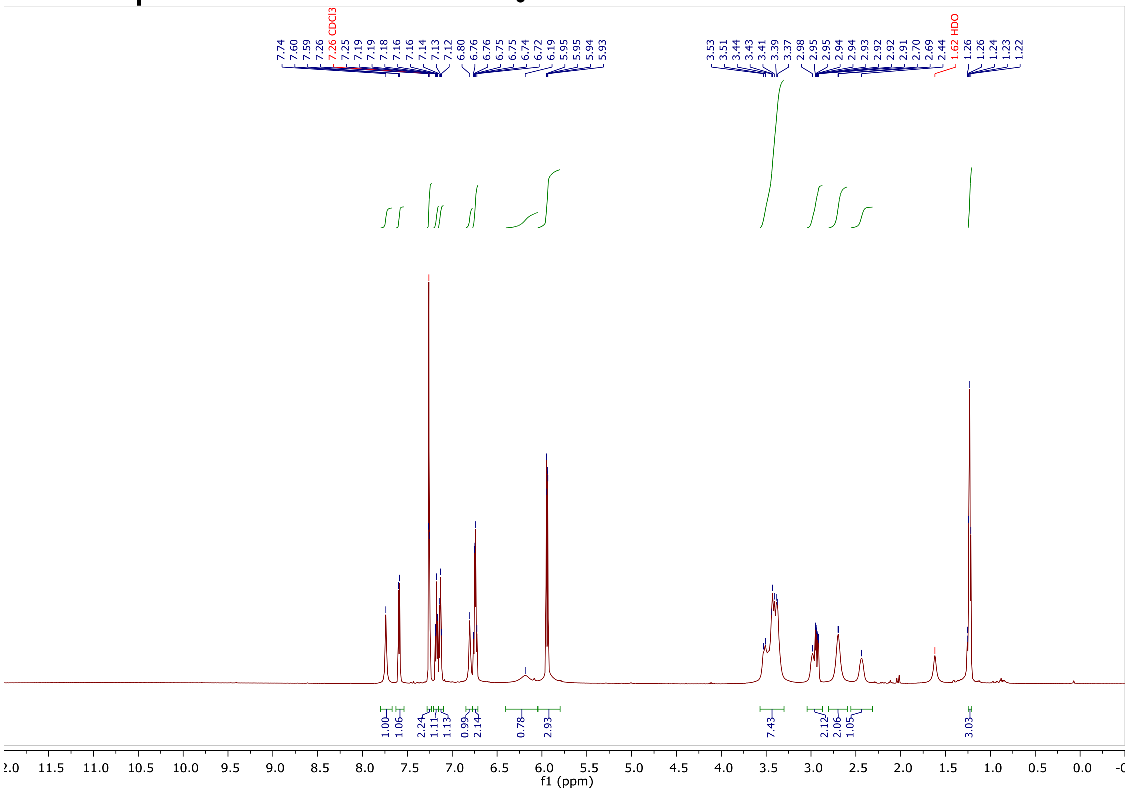

#### $^1H$ NMR spectrum of DO-40B in $CDCl_3$

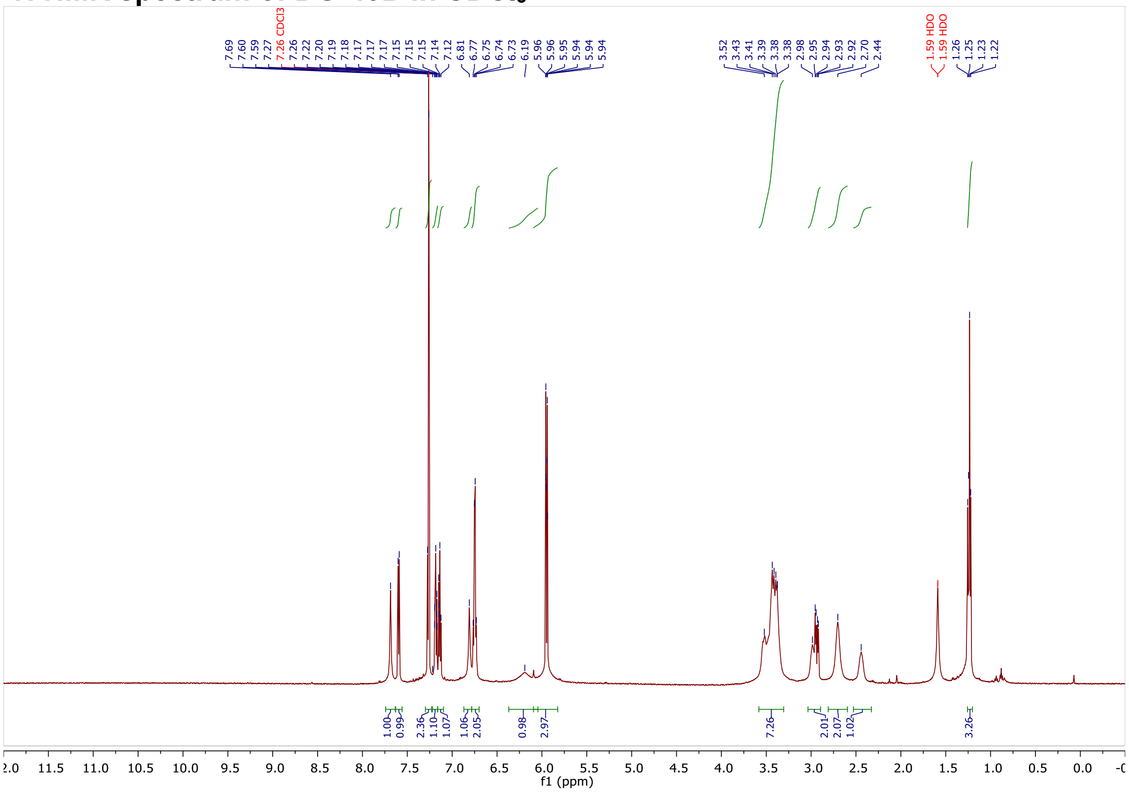

#### $^{1}H$ NMR spectrum of DO-35A in $CDCl_3$

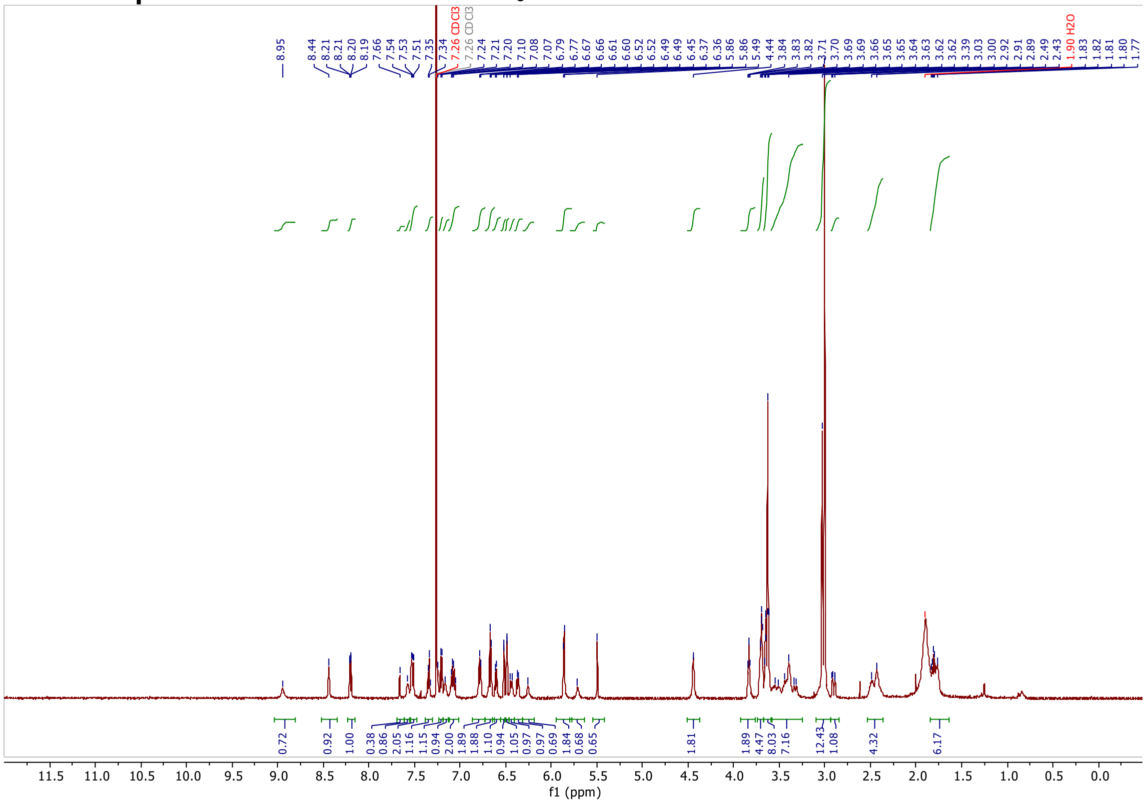

#### $^{1}H$ NMR spectrum of DO-35B in $CDCl_3$

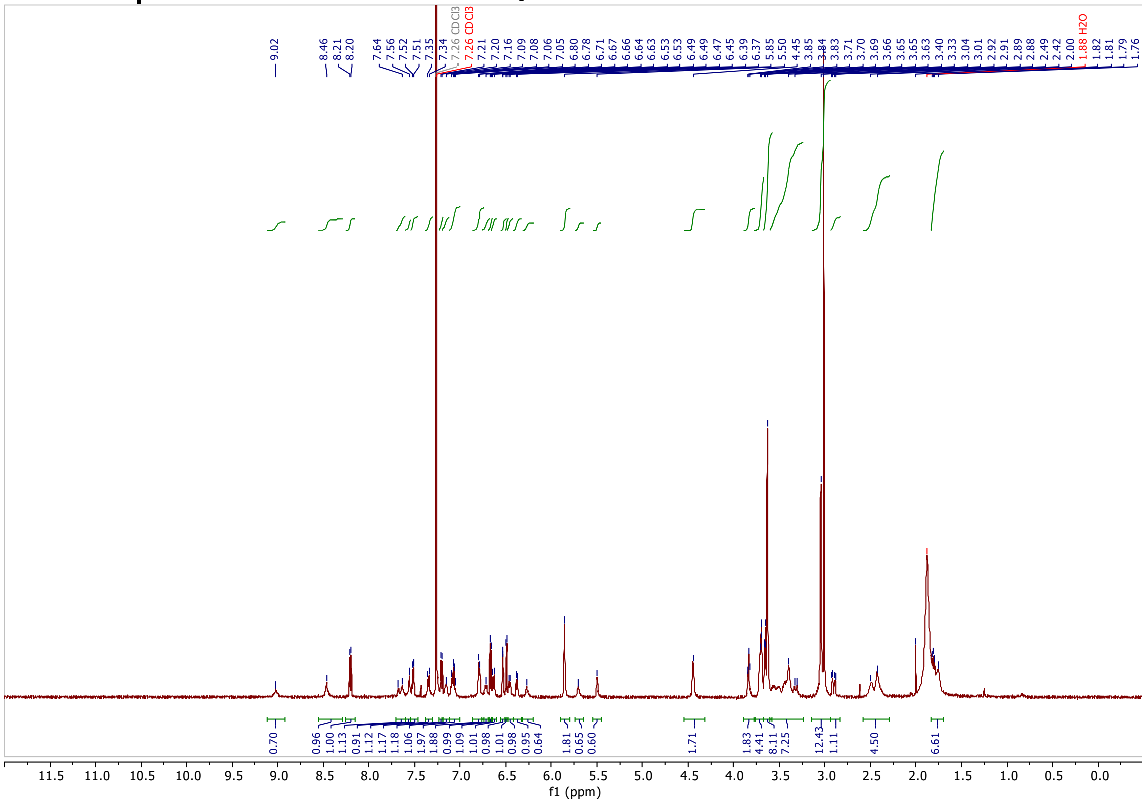

#### $^{1}H$ NMR spectrum of DO-39A in $CDCl_3$

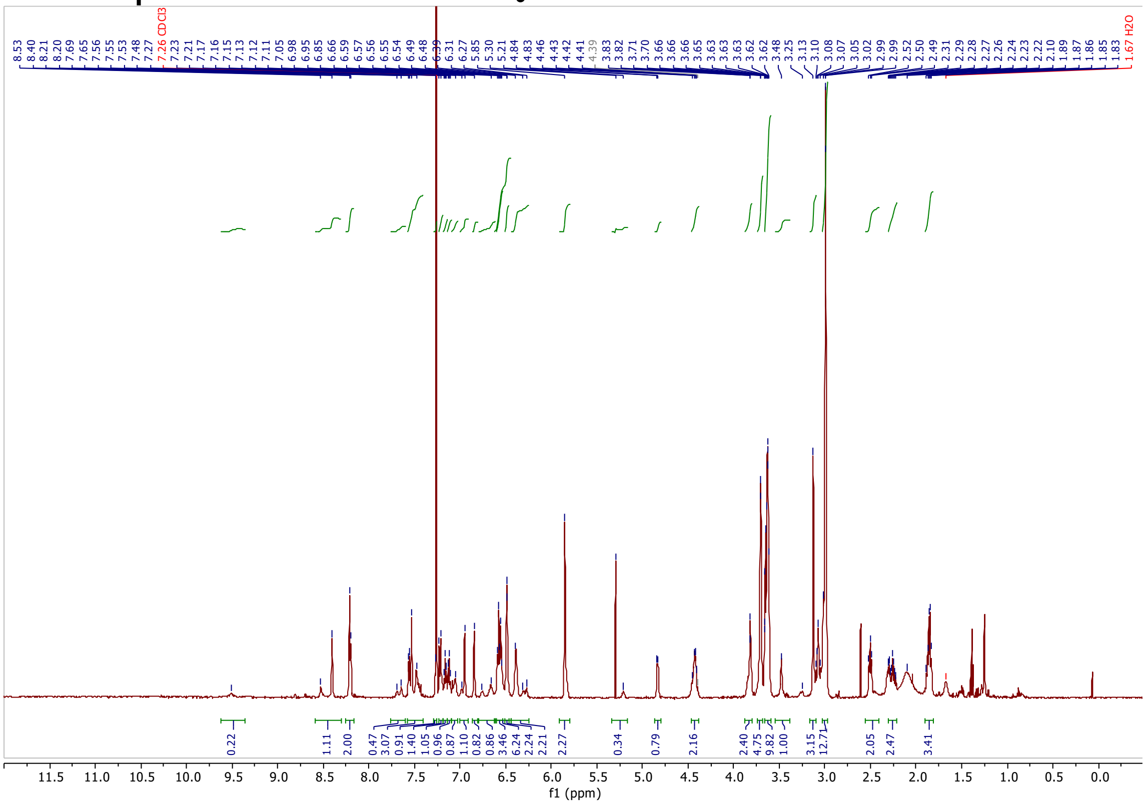

#### $^{1}H$ NMR spectrum of DO-39B in $CDCl_3$

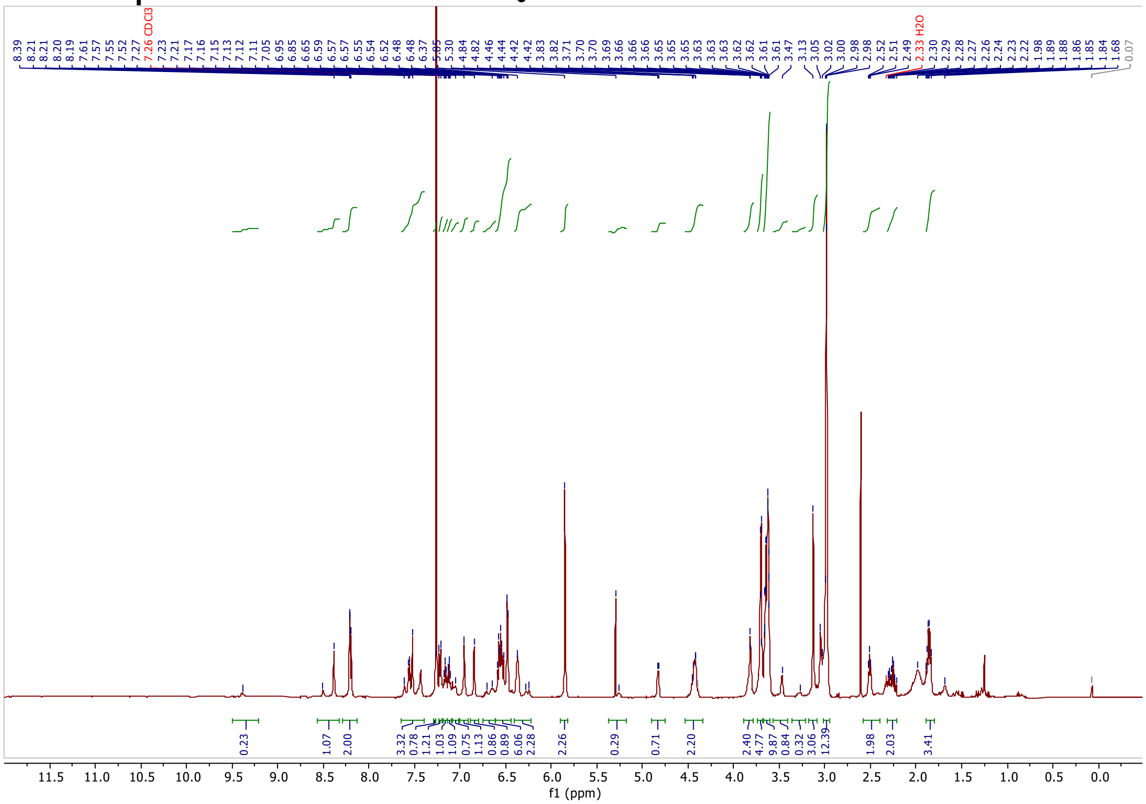
